# *Listeria monocytogenes* utilizes the ClpP1/2 proteolytic machinery for fine-tuned substrate degradation under heat stress

**DOI:** 10.1101/2021.05.25.445618

**Authors:** Dóra Balogh, Konstantin Eckel, Christian Fetzer, Stephan A. Sieber

## Abstract

*Listeria monocytogenes* exhibits two ClpP isoforms (ClpP1/ClpP2) which assemble into a heterooligomeric complex with enhanced proteolytic activity. Herein, we demonstrate that the formation of this complex depends on temperature and reaches a maximum ratio of about 1:1 at heat shock conditions, while almost no complex formation occurred below 4°C. In order to decipher the role of the two isoforms at elevated temperatures, we constructed *L. monocytogenes* ClpP1, ClpP2 and ClpP1/2 knockout strains and analyzed their protein regulation in comparison to the wild type (WT) strain via whole proteome mass-spectrometry (MS) at 37 °C and 42 °C. While Δ*clpP1* strain only altered the expression of very few proteins, Δ*clpP2* and Δ*clpP1/2* strains revealed the dysregulation of many proteins at both temperatures. These effects were corroborated by crosslinking co-immunoprecipitation MS analysis. Thus, while ClpP1 serves as a mere enhancer of protein degradation in the heterocomplex, ClpP2 is essential for ClpX binding and thus functions as a gatekeeper for substrate entry. Applying an integrated proteomic approach combining whole proteome and co-immunoprecipitation datasets, several putative ClpP2 substrates were identified in the context of different temperatures and discussed with regards to their function in cellular pathways such as the SOS response.

## Introduction

*Listeria monocytogenes* is a highly stress resistant pathogenic bacterium that can survive under rapidly changing conditions (Bucur et al., 2018; Radoshevich & Cossart, 2018). In order to cope with different stresses, the cells must detect environmental changes and promptly adjust protein expression as well as turnover in a strictly regulated manner. One characteristic trait of *L. monocytogenes* is its growth at various temperatures ranging from −0.4 to +45 °C posing a major challenge for adapting its cellular physiology (Bucur et al., 2018). For example, heat shock induces the SOS response which is initiated by autocleavage of LexA, the repressor of the SOS genes (Michel, 2005; van der Veen et al., 2007). N- and C-terminal LexA domains (NTD and CTD, respectively) are further digested by bacterial proteases such as ClpXP (see below) to activate the SOS pathway (Cohn et al., 2011; Little & Gellert, 1983; Neher et al., 2003). In *L. monocytogenes*, 28 genes have been identified to be under control of LexA (van der Veen et al., 2010). Most of them are DNA polymerases required for DNA repair. Furthermore, the induction of the SOS pathway inhibits bacterial growth, probably in order to prevent cell division after incomplete DNA replication (Kawai et al., 2003; van der Veen et al., 2010).

In addition to gene regulation, heat stress generates damaged proteins, which need to be efficiently removed by the cellular proteolytic machinery. In bacteria, several proteases are capable of this process. These include caseinolytic protease P (ClpP) which, in concert with its cognate chaperones, digests misfolded protein substrates. ClpX is a hexameric ATP-dependent chaperone which recognizes protein substrates and directs unfolded peptide chains into the tetradecameric barrel of the ClpP serine protease for degradation (Baker & Sauer, 2012). Some bacteria such as *L. monocytogenes* encode two ClpP isoforms (ClpP1 and ClpP2) with yet largely unknown cellular roles (Dahmen et al., 2015; Hall et al., 2017; Mawla et al., 2020; Pan et al., 2019; Zeiler et al., 2011). In *L. monocytogenes* ClpP1 exhibits low sequence homology to ClpP isoforms from other bacteria and is expressed as an inactive heptamer with an impaired catalytic triad. Co-expression with ClpP2, a close homolog of other ClpP isoforms, yields a heterotetradecamer assembly composed of one ClpP1 and one ClpP2 heptamer ring, ClpP1_7_P2_7_, here referred to as ClpP1/2 (Dahmen et al., 2015). In association with ClpX, this heterocomplex exhibits enhanced proteolytic activity in comparison to the corresponding uniform ClpX_6_P2_14_ complex. Structural studies revealed that within this complex ClpP2 serves as a template to force the impaired catalytic triad of ClpP1 into an aligned conformation which enables substrate digestion (Dahmen et al., 2015). Moreover, recent cryo-EM data confirmed that ClpX solely docks via ClpP2 to the heterocomplex as ClpP1 lacks cognate chaperone binding sites (Gatsogiannis et al., 2019). It is thus assumed that ClpP1 is needed by *L. monocytogenes* under certain conditions to enhance proteolytic turnover and clearance of misfolded proteins.

It is hitherto unknown why some bacteria have homotetradecameric and others heterotetradecameric ClpPs. In this study, we revealed the thermosensing ability of ClpP1/2 heterooligomerization and investigated the unique cellular functions of ClpP1 and ClpP2 in *L. monocytogenes*. To achieve this, the phenotypes of Δ*clpP1*, Δ*clpP2* and double knockout (Δ*clpP1/2*) strains were examined in an integrative proteomic approach using mass spectrometry-based whole proteome analysis and co-immunoprecipitation. Our data suggest that ClpP2 plays an important role to mediate substrate recognition of e.g. proteins involved in stress response while ClpP1 is a mere enhancer of proteolytic turnover.

## Results

### ClpP1 and ClpP2 form a heterocomplex at elevated temperatures

Previous transcription analyses showed that both *clpP* genes exhibit up to 7-fold higher expression levels under heat stress (Dahmen et al., 2015; van der Veen et al., 2007), indicating that heterocomplex formation is preferred at high temperatures and might have a specific biological role under these conditions. In line with this observation, heterologous co-expression and purification of *L. monocytogenes* ClpP1/2 in *E. coli* revealed that the heterocomplex is unstable at low temperatures (4 °C) and stable tetradecameric ClpP1/2 could only be obtained when the whole purification process after cell lysis was performed at room temperature (~26 °C, Figure 1a,b). Interestingly, *M. tuberculosis* ClpP1 and ClpP2 also heterooligomerize at elevated temperatures (Leodolter et al., 2015), which suggests that heat sensing could be a conserved biological function of ClpP. To assess whether the temperature-dependent stabilization is a general feature of ClpP1/2 and not a result of the co-expression and purification conditions, we measured heterooligomerization of separately overexpressed and purified ClpP1_7_ and ClpP2_14_ at different temperatures.

**Figure 1.**
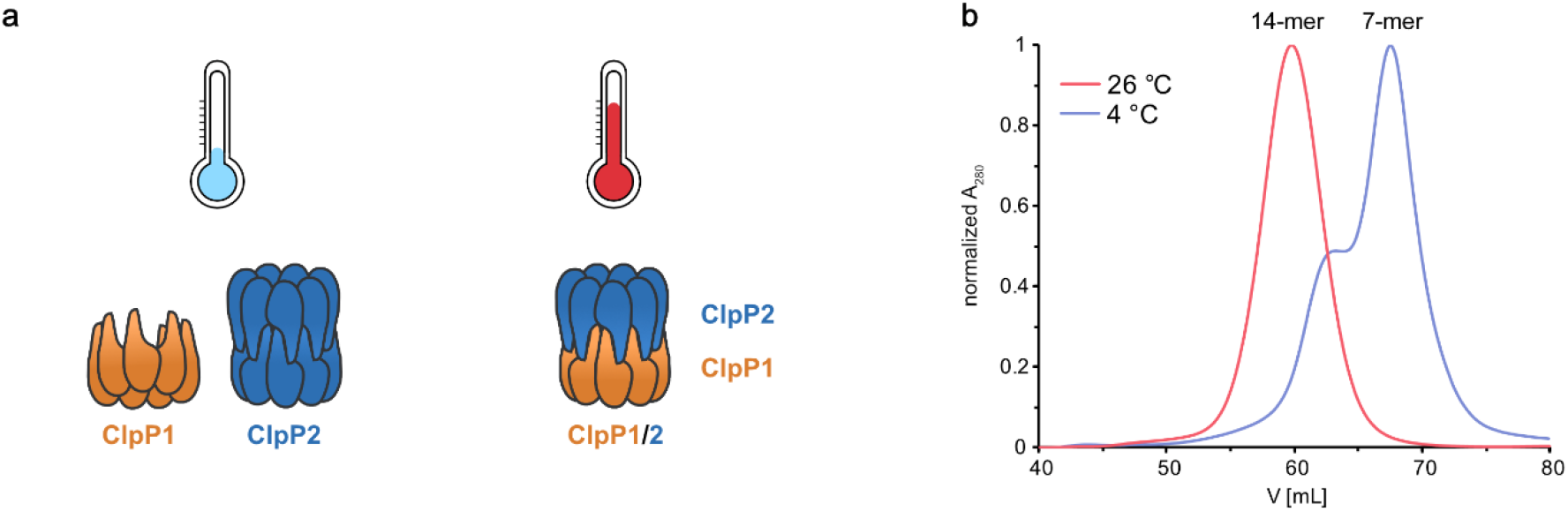
Purification of ClpP1/2 at 4°C and at room temperature. **a** Schematic representation of ClpP1 (orange) and ClpP2 (blue) compositions at different temperatures according to size-exclusion chromatography. **b** Size-exclusion chromatography was performed on a Superdex200pg 16/60 column of co-expressed ClpP1/2 purified at 4 °C and 26 °C. Purifications of *L. monocytogenes* ClpP1/2 at 4 °C yielded a mixture of heptameric ClpP1 and tetradecameric ClpP2 (blue curve with shoulder), whereas a tetradecameric ClpP1/2 heterocomplex was obtained at room temperature (red curve).

For this, equal amounts of both purified enzymes were mixed and incubated at temperatures ranging from 0 °C to 48 °C. The samples were subjected to analytical size-exclusion chromatography (SEC), and the protein composition of the tetradecamer peak was analyzed by intact protein mass spectrometry (ip-MS) (Figure 2a). The ratio of the tetradecamer (ClpP2_14_ and ClpP1/2_14_) and heptamer peaks (ClpP1_7_) differed temperature-dependently with the highest 14-mer amount observed at 42 °C (Figure 2b). Ip-MS analysis revealed an increasing ClpP1 fraction within the tetradecameric complex up to 37-42 °C with a maximum content of about 40-44% (Figure 2c). However, at 48 °C, the 7-mer:14-mer ratio declined to a 1:1 ratio. Accordingly, the ClpP1 partition decreased. As a control, the ClpP1/2 complex assembled at 42 °C was cooled down to 0 °C which resulted in a disassembly of the newly built heterooligomers suggesting that heterocomplex formation is reversible (Figure 2d). In order to rule out the existence of ClpP1_14_ homocomplexes, we incubated ClpP1 at 42 °C. No shift in the chromatogram compared to 0 °C occurred, which implies that ClpP1 is not able build homotetradecamers even under elevated temperatures (Figure 2e).

**Figure 2.**
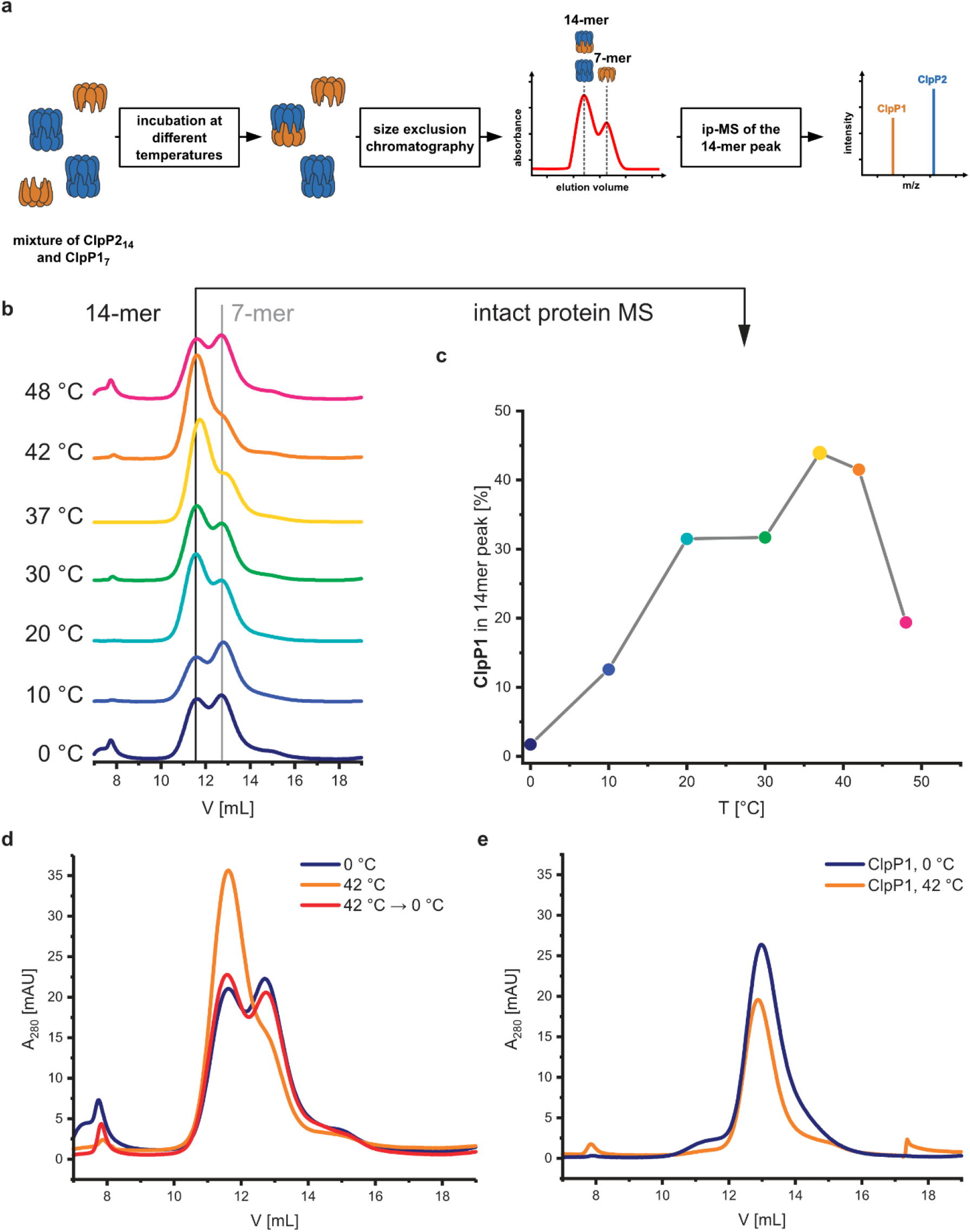
Temperature-dependent formation of the ClpP1/2 heterocomplex. **a** Scheme of the SEC/ip-MS workflow. Orange: ClpP1, blue: ClpP2.**b** Size-exclusion chromatography of ClpP1_7_ and ClpP2_14_ after incubation at the indicated temperatures for 30 min. Black line: tetradecamer, gray line: heptamer. **c** Percentage of ClpP1 in the 14-mer peaks of panel b, measured by intact protein mass spectrometry. **d** Size-exclusion chromatography of ClpP1/2 after incubation at 0 °C for 30 min (blue), 42 °C for 30 min (orange) and 42 °C for 30 min followed by 0 °C for 30 min (red). **e** Size-exclusion chromatography of ClpP1_7_ after incubation at 0 °C for 30 min (blue) and at 42 °C for 30 min (orange).

ClpP1 is not active by itself, however, in association with the heterocomplex it exhibits ten times higher protease activity per subunit compared to the ClpP2 homocomplex (Balogh et al., 2017; Dahmen et al., 2015). In order to assess whether the heterocomplex formation translates to increased protease activity at high temperatures, we monitored the degradation of GFP-SsrA by ClpXP in the presence of an ATP regeneration system (Kim et al., 2000). Using this assay, we compared the protease activity of mixed ClpP1_7_ and ClpP2_14_ to solely ClpP2_14_ at different temperatures. While ClpP1 alone is known to be inactive because of its impaired catalytic triad (Ser98, His123, *Asn172*) and its inability to bind AAA+ chaperones (Balogh et al., 2017; Dahmen et al., 2015; Gatsogiannis et al., 2019; Zeiler et al., 2013), co-incubation with ClpP2 at 37 °C and 42 °C resulted in an elevated proteolytic activity compared to a ClpP2 homocomplex at the same respective temperature (Figure 3). The overall slower kinetics of the GFP degradation at 42 °C are attributed to the low thermal stability of ClpX and the ATP regenerating enzyme creatine kinase (Wu et al., 2011).

**Figure 3.**
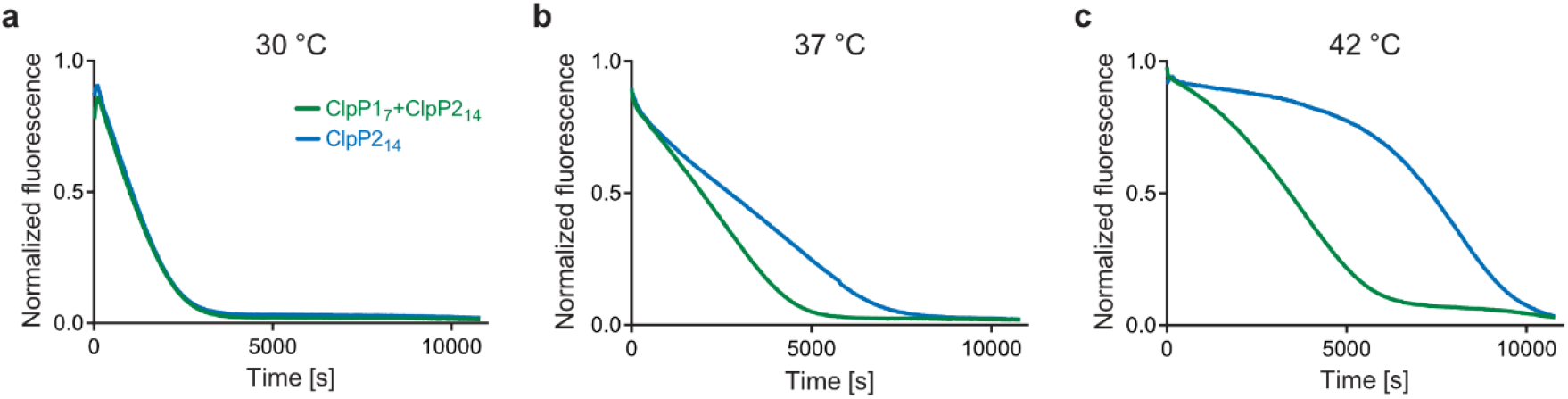
Protease activity of ClpP1_7_ and ClpP2_14_ at different temperatures. ClpP (green line: 0.1 μM ClpP2_14_ and 0.2 μM ClpP1_7_, blue line: 0.1 μM ClpP2_14_) and 0.4 μM ClpX were pre-incubated for 30 min at 30 °C (**a**), 37 °C (**b**) and 42 °C (**c**), subsequently the degradation of 0.4 μM GFP-SsrA was measured. Means of triplicates are shown. The experiments were independently repeated with qualitatively identical results (Figure S3).

### Intracellular heterooligomerization of ClpP1 and ClpP2 under heat stress

Next, we set out to investigate whether temperature-dependent heterooligomerization also occurs in living *L. monocytogenes* as a response to heat stress. For this purpose, we first quantified ClpP1 and ClpP2 levels via Western blot at low and high temperatures to investigate if the previously observed increased expression of both *clpP* genes translates to the protein level (Dahmen et al., 2015).

In order detect ClpP isomers selectively, we inserted a 2×myc tag at the end of the endogenous *clpP* genes. The *L. monocytogenes clpP1(191)*::2×myc and *L. monocytogenes clpP2(199)*::2×myc strains constitutively expressed C-terminally myc-tagged ClpP1 (ClpP1-2×myc) and ClpP2 (ClpP2-2×myc) respectively, which can be visualized with an anti-c-Myc antibody in a western blot. In addition, the myc-tag also allows for co-immunoprecipitation (co-IP) experiments to study the interaction of ClpP1 and ClpP2 *in situ*.

Indeed, we observed strongly increased expression of both isoforms at elevated temperatures compared to 10°C and 20 °C, corroborating previous gene expression studies (Figure S1) (Dahmen et al., 2015). This increase is especially pronounced for ClpP1, since its expression is lower compared to ClpP2 at temperatures < 42 °C. As both isoforms are highly abundant at elevated temperatures, we investigated a potential role of the proteins under heat stress.

Yet, the extremely low expression of ClpP1 at low temperatures represents a challenge for co-IP experiments when studying their temperature-dependent interactions *in situ*, especially when choosing ClpP1 as the bait protein. Despite this limitation, we carried out co-IP experiments at high and low temperatures with an immobilized anti-c-Myc antibody in the presence of a disuccinimidyl sulfoxide (DSSO) crosslinker to stabilize transient protein-protein interactions (Fux et al., 2019). The captured proteins were subjected to a tryptic digest, and the isolated peptides were measured by LC-MS/MS. As expected, when ClpP1-2×myc was used as bait no difference in ClpP2 enrichment could be observed at 42 °C compared to 20 °C (Figure S2a,b). This result is likely attributed to the low abundance of heptameric ClpP1 at 20 °C which under the huge excess of ClpP2 could form sufficient amounts of heterocomplex. In contrast, as ClpP2 is generally abundant, it represents a more robust reference protein for this study. In fact, when ClpP2-2×myc was used as a bait, analysis of the ClpP1 intensities revealed a 6-times higher enrichment at 42 °C compared to 20 °C (Figure S2c,d). Despite this encouraging result, it is difficult to draw general conclusions due to the lack of a reliable ClpP1 expression at low temperatures.

### Phenotypic characterization of *L. monocytogenes* Δ*clpP* mutants

To further investigate the cellular role of ClpP1 and ClpP2, we constructed Δ*clpP1* and Δ*clpP2* single mutants, as well as a Δ*clpP1/2* double knockout strain (Figure 4a, top) in *L. monocytogenes* EGD-e (WT). Growth curves of the mutants show that the single mutants grow at a similar rate to the wild type strain but ΔclpP2 reaches a higher optical density in the stationary phase (Figure 4b, Figure S4). The double mutant ΔclpP1/2 grows substantially slower than all other investigated strains, but shows the highest optical density in the stationary phase.

**Figure 4.**
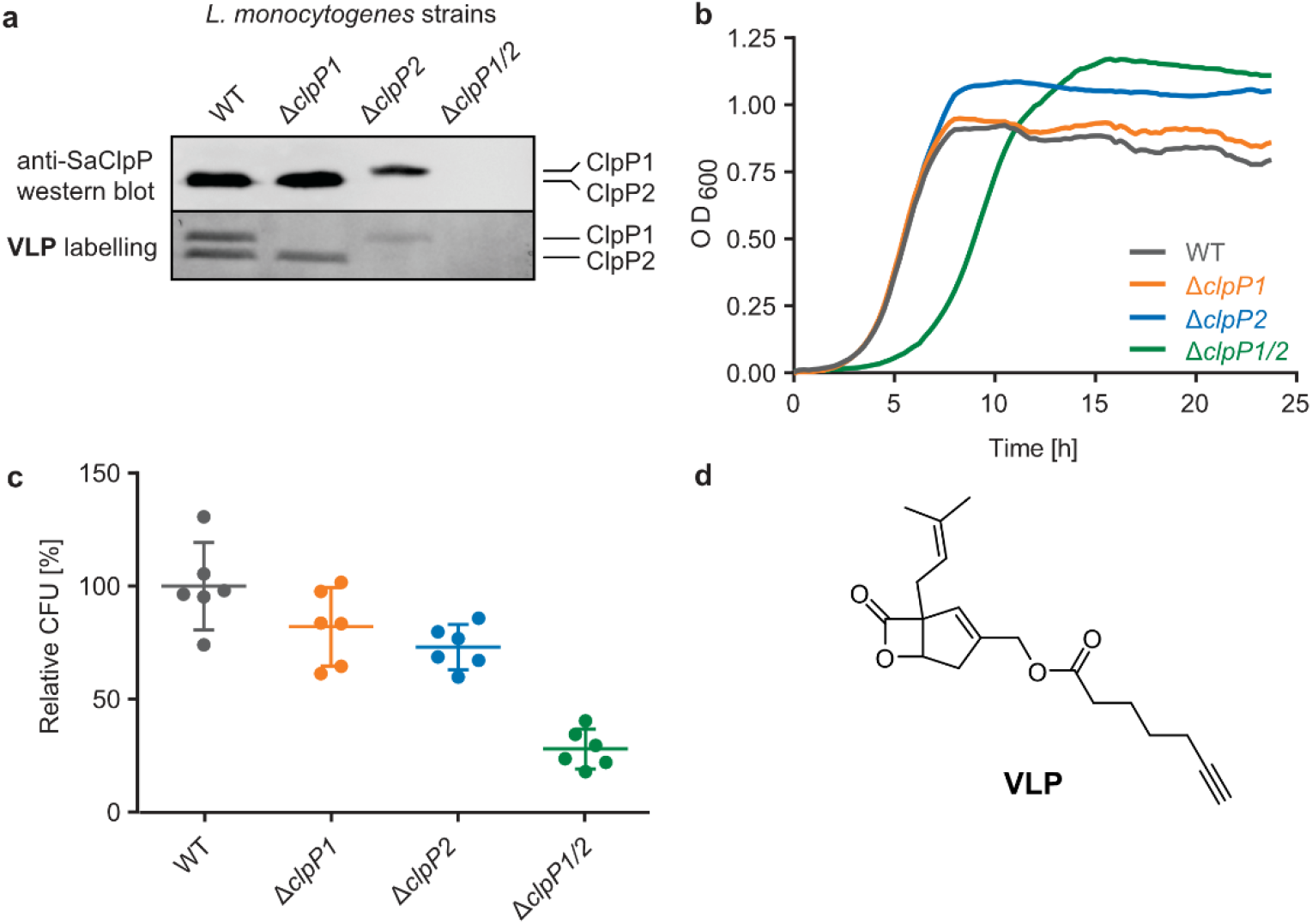
*L. monocytogenes* Δ*clpP* mutants. **a** Validation of the Δ*clpP* mutants by western blot (top) and by fluorescent labelling with vibralactone probe (bottom). **b** Growth curves of the Δ*clpP* mutants in BHI medium at 37 °C. Means of triplicates are shown. The experiment was independently repeated with qualitatively identical results (Figure S4a). **c** Intracellular growth of the Δ*clpP* mutants in murine macrophages. CFUs were determined after 7 h, and normalized to WT as 100% (n = 6, two independent experiments in triplicates were performed, mean ± 95% confidence interval). **d** Structure of the vibralactone probe.

ClpP2 is known to be important for intracellular growth in macrophages and we thus investigated the impact of all mutants in this process as well (Gaillot et al., 2000). Mouse-derived macrophages were infected with *L. monocytogenes* EGD-e (WT), Δ*clpP1*, Δ*clpP2* and Δ*clpP1/2* and colony forming units (CFUs) determined after 7 hours (Figure 4c). All mutants were able to replicate inside the cells, with comparable growth behaviors as observed in medium. Contrary to previous findings (Gaillot et al., 2000), the intracellular growth of Δ*clpP2* was only weakly inhibited which might be attributed to the use of a different strain by Gaillot et al. (*L. monocytogenes* LO28).

We next assessed the *in situ* activity of both ClpPs by labelling the whole *L. monocytogenes* cells with vibralactone probe (VLP) (Figure 4d). Vibralactone is the only known small molecule, which is able to label both ClpP1 and ClpP2 of *L. monocytogenes* by binding to their active site serine (Zeiler et al., 2011). VLP is equipped with a terminal alkyne tag which enables coupling to an azide-functionalized rhodamine dye via copper-catalyzed click chemistry (Huisgen, 1961; Rostovtsev et al., 2002; Tornøe et al., 2002). This way, proteins that covalently bind VLP can be visualized by fluorescence on a polyacrylamide gel. As observed previously, VLP is able to label both ClpP2 and ClpP1 in *L. monocytogenes* EGD-e (Figure 4a, bottom). In line with the lack of proteolytic activity (Dahmen et al., 2015; Zeiler et al., 2013), only a weak ClpP1 band is observed in Δ*clpP2* which may result from some residual binding to the active site. In addition, as expected a strong ClpP2 signal is detected in Δ*clpP1.*

### Whole-proteome analysis of ClpP1 and ClpP2 deletion mutants

ClpP is required for the maintenance and regulation of the proteome by clearing damaged proteins and degrading transcription factors. So far, the specific roles of ClpP1 and ClpP2 in *L. monocytogenes* are elusive. We analyzed whole proteomes of *L. monocytogenes* EGD-e (WT), Δ*clpP1*, Δ*clpP2* and Δ*clpP1/2* grown to early stationary phase at 37 °C and 42 °C to identify proteomic changes upon deletion of one or both proteins.

At 37 °C, the proteome of Δ*clpP1* does not differ markedly from the wild type (Figure 5a, Table S1) but in Δ*clpP2* and in Δ*clpP1/2* many proteins are dysregulated (Figure 5b,c). The dysregulated proteins in Δ*clpP2* and Δ*clpP1/2* are highly overlapping: 89% of the proteins that are upregulated in Δ*clpP1/2* compared to the wild type are also upregulated in Δ*clpP2* and the same applies for 82% of the downregulated proteins in the double mutant (Figure 5d,e). However, a notable difference is the exclusive downregulation of 123 proteins solely in Δ*clpP2* compared to the double mutant. Surprisingly, the different phenotypes of both mutants is not reflected by the respective proteome changes.

**Figure 5.**
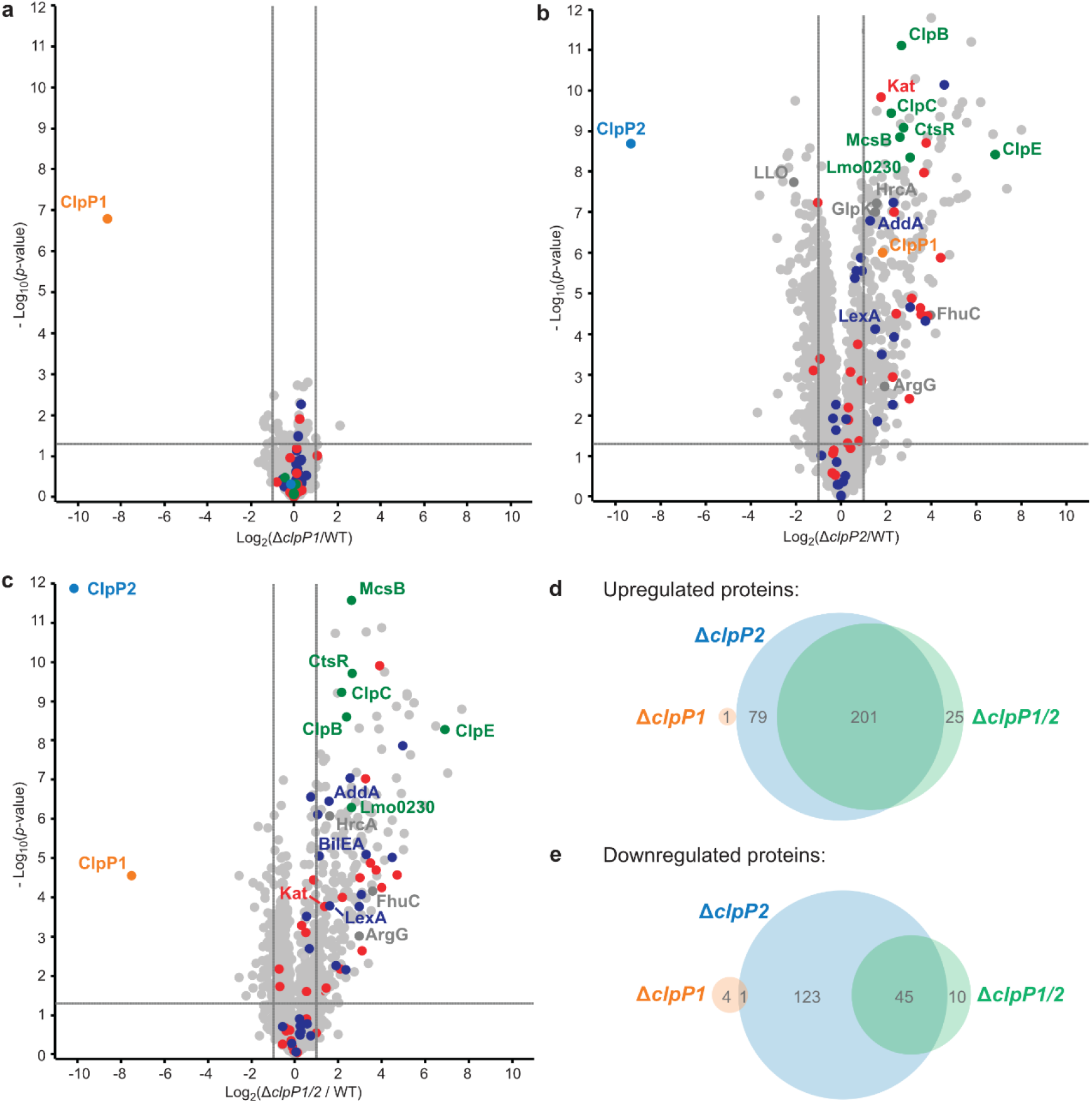
Whole-proteome analysis of the *L. monocytogenes* Δ*clpP* mutants at 37 °C. **a–c** Proteomes of *L. monocytogenes* Δ*clpP1* (**a**), Δ*clpP2* (**b**) and Δ*clpP1/2* (**c**) compared to the WT. Bacterial cultures were grown to stationary phase at 37 °C. − Log_10_ *p*-values from two-sample Student’s *t*-test are plotted against log_2_ ratios of LFQ protein intensities. The vertical grey lines show 2-fold enrichment, the horizontal grey lines show −log_10_ t-test *p*-value = 1.3. Samples were prepared in triplicates in two independent experiments (n = 6). Class III heat shock proteins (green), SOS response proteins (dark blue) and iron-containing proteins (red) are highlighted. Other proteins mentioned in the text are highlighted in dark grey if they are significantly dysregulated in the respective plot. ClpP1 and ClpP2 are shown in orange and blue respectively. **d–e** Venn-diagrams showing the up-(**d**) and downregulated (**d**) proteins in the proteomes of the Δ*clpP* mutants compared to the WT (fold enrichment ≥ 2, −log_10_ t-test *p*-value ≥ 1.3, ClpP1 and ClpP2 excluded).

UniProt keyword and Gene Ontology Biological Process (GOBP) term analyses of the proteomic data were performed with the aGOtool (agotool.org) (Table S3 – Table S10) (Schölz et al., 2015). All proteins detected at 37 °C in the whole proteomes of *L. monocytogenes* EGD-e and all mutants were combined after categorical filtering and used as background. Among the upregulated proteins, the GOBP term “response to stimulus” and “regulation of transcription” was significantly enriched in both Δ*clpP2* and in Δ*clpP1/2* (Table S3). Notably, SOS response-related terms (cellular response to DNA damage stimulus, DNA repair) were specifically enriched only in Δ*clpP1/2*. This indicates that both ClpPs are needed for full regulation of the SOS response in *L. monocytogenes*. Activation of the SOS response inhibits cell division in *L. monocytogenes* (van der Veen et al., 2007) and in *E. coli* (Miller et al., 2004), which rationalizes the observed slower growth of Δ*clpP1/2* compared to the wild type.

Additionally, the class III heat shock proteins (CtsR, McsB, ClpB, ClpC, ClpE and the Lmo0230 protein) were upregulated in both Δ*clpP2* and Δ*clpP1/2*. The class I heat shock proteins were not overexpressed, except for their repressor, HrcA. Most of the class II HSPs (except for GlpK and BilEA, which is also an SOS response protein) and their positive regulator σ^B^ were also not dysregulated. Of the 28 proteins, which have been found in a genome-wide screen for temperature sensitivity (Van Der Veen et al., 2009), only two (ClpB and AddA) were significantly upregulated in Δ*clpP2* and Δ*clpP1/2*. This, and the fact that the class I and II heat shock proteins were not induced, highlights the differences between the stress caused by *clpP2* deletion and heat stress, even though class III heat shock proteins and parts of the SOS response are induced in the mutants lacking *clpP2*. Iron containing and iron-sulfur proteins were also significantly upregulated in Δ*clpP2* and in Δ*clpP1/2*. In *S. aureus*, it has been shown that ClpP degrades damaged iron-sulfur proteins (Flynn et al., 2003; Guillon et al., 2009), which could also be the case in *L. monocytogenes*. Additionally, ClpP has been connected to iron homeostasis and maintaining the oxidative balance inside the cell (Farrand et al., 2015; Frees et al., 2003; Michel et al., 2006).

At 42 °C, in general more proteins are dysregulated than at 37 °C in all whole proteomes. Although there are more proteins dysregulated for Δ*clpP1* at 42 °C compared to 37 °C, there is surprisingly little impact of a ClpP1 deletion on the proteome level (Figure 6a, Table S2). Among the upregulated proteins are the two virulence-associated proteins internalin B (InlB) and listeriolysin O (LLO). Internalin B plays a role in receptor-mediated endocytosis of non-phagocytic cells, whereas listeriolysin O is a pore-forming toxin needed for subsequent vacuole opening to enter the cytosol of infected cells (Radoshevich & Cossart, 2018). In addition, FhuC is upregulated, which is an ABC ATPase involved in the membrane transport of iron(III)hydroxamates in *S. aureus* (Speziali et al., 2006).

**Figure 6.**
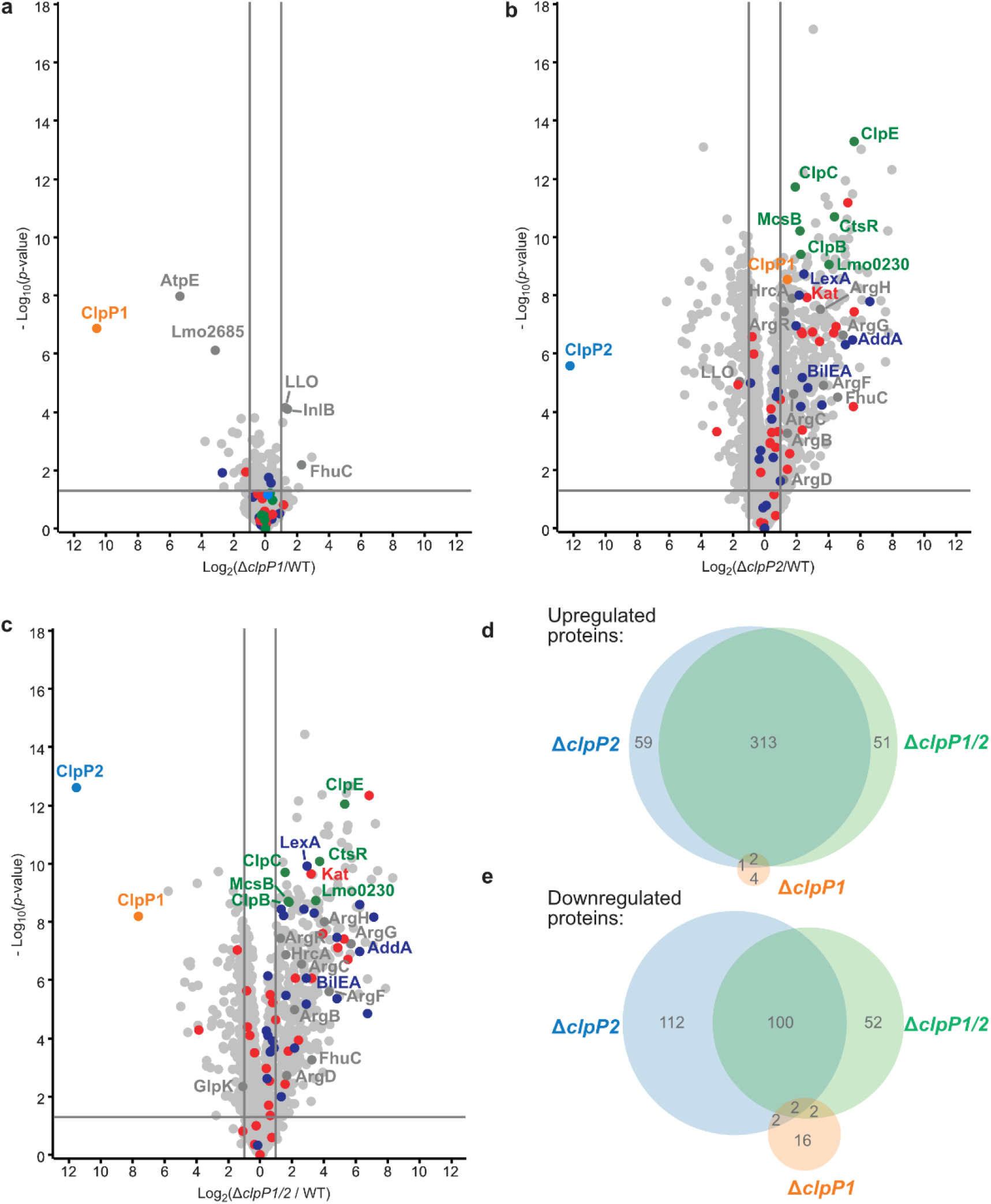
Whole-proteome analysis of the *L. monocytogenes* Δ*clpP* mutants at 42 °C. **a–c** Proteomes of *L. monocytogenes* Δ*clpP1* (**a**), Δ*clpP2* (**b**) and Δ*clpP1/2* (**c**) compared to the WT. Bacterial cultures were grown to stationary phase at 42 °C. − Log_10_ *p*-values from two-sample Student’s *t*-test are plotted against log_2_ ratios of LFQ protein intensities. The vertical grey lines show 2-fold enrichment, the horizontal grey lines show −log_10_ t-test *p*-value = 1.3. Samples were prepared in triplicates in two independent experiments (n = 6). Class III heat shock proteins (green), SOS response proteins (dark blue) and iron-containing proteins (red) are highlighted. Other proteins mentioned in the text are highlighted in dark grey if they are significantly dysregulated in the respective plot. ClpP1 and ClpP2 are shown in orange and blue respectively. **d–e**Venn-diagrams showing the up-(**d**) and downregulated (**d**) proteins in the proteomes of the Δ*clpP* mutants compared to the WT (fold enrichment ≥ 2, −log_10_ t-test *p*-value ≥ 1.3, ClpP1 and ClpP2 excluded).

The most significantly downregulated proteins for Δ*clpP1* at 42 °C are the F-ATPase subunit c (AtpE) and Lmo2685, a component of the phosphotransferase system (PTS). Yet, we identified no protein which is upregulated in Δ*clpP1* and Δ*clpP1/2,* but not in Δ*clpP2* at both temperatures. Since ClpX and most likely other chaperones bind solely to ClpP2 (Gatsogiannis et al., 2019), a deletion of ClpP1 is expected to solely adjust the speed of substrate degradation with little impact on the substrate scope itself. Thus, static proteome analysis may not capture the dynamics of protein digest, as during cell harvest and lysis ClpP2 still retains its activity, which could diminish observed proteome changes between Δ*clpP1* and the wild type.

For Δ*clpP2* and Δ*clpP1/2,* again many more proteins are dysregulated (Figure 6b,c), with 375 and 366 upregulated proteins for either deletion mutant. 216 proteins are downregulated in Δ*clpP2* and 156 proteins are downregulated in Δ*clpP1/2.* Yet, there is still a high overlap between dysregulated proteins of Δ*clpP2* and Δ*clpP1/2*. 86% of proteins upregulated in Δ*clpP1/2* are also upregulated in Δ*clpP2* and the same applies to 65 % of downregulated proteins of Δ*clpP1/2* (Figure 6d,e). Again a notable difference is the downregulation of 112 proteins in Δ*clpP2*, which are not affected in Δ*clpP1/2.* In line with whole proteomes at 37 °C, both deletion mutants show an upregulation of iron- or iron-sulfur-containing proteins and class III heat shock proteins. Interestingly, for Δ*clpP2* at 42 °C we could identify SOS-related GOBP terms (e.g. DNA repair, base-excision repair, cellular response to DNA damage stimulus) to be upregulated, which distinguishes it from the same deletion mutant at 37 °C (Table S7, S8). Yet, the actual term “SOS response” is only significantly upregulated for Δ*clpP1/2* at 42 °C, further supporting the effect of both ClpPs on the SOS response regulation.

In addition, the pyrimidine and especially the UMP de novo biosynthesis is highly downregulated for Δ*clpP2* and Δ*clpP1/2* at both temperatures (Table S9, S10). This downregulation is especially pronounced at 42 °C with nearly every protein of the *pyr* operon affected. Recently, a downregulation of the UMP biosynthesis was also discovered for Δ*clpP* of *S. aureus*, which was subsequently confirmed on the metabolite level (Kirsch et al., 2021). In contrast, the purine biosynthesis is not heavily affected in *L. monocytogenes*, which differs from the *S. aureus* Δ*clpP* proteome.

There are also some notable differences to the 37 °C whole proteomes. A majority of arginine biosynthetic proteins (ArgB, ArgC, ArgD, ArgF, ArgG, ArgH) is upregulated only at 42 °C both for Δ*clpP2* and Δ*clpP1/2*, together with their repressor ArgR.

### Co-immunoprecipitation of ClpP1 and ClpP2

In order to identify specific interaction partners of ClpP1 and ClpP2, we conducted co-immunoprecipitation (co-IP) experiments. The Δ*clpP1* and Δ*clpP2* mutants were grown to stationary phase at 37 °C and 42 °C, respectively, and interacting proteins were covalently crosslinked with DSSO (XL-co-IP). The ClpPs were precipitated with a polyclonal anti-ClpP antibody and binding partners of each ClpP isoform were selectively pulled down. The precipitated proteins were digested with trypsin and analyzed by LC-MS/MS.

451 significantly enriched proteins were found for ClpP1 and 468 for ClpP2 at 37 °C as well as 232 and 145 at 42 °C, respectively (Figure 7). Overall, the co-IP results can be classified into two categories of proteins: a) chaperones and adaptor proteins that engage in a classical protein-protein interaction with ClpP and b) protein substrates that are bound to the chaperones and adaptors and are subsequently digested by ClpP. As it is not possible to assign hit proteins to one of these groups solely by the co-IP data, more in-depth analysis is required.

**Figure 7.**
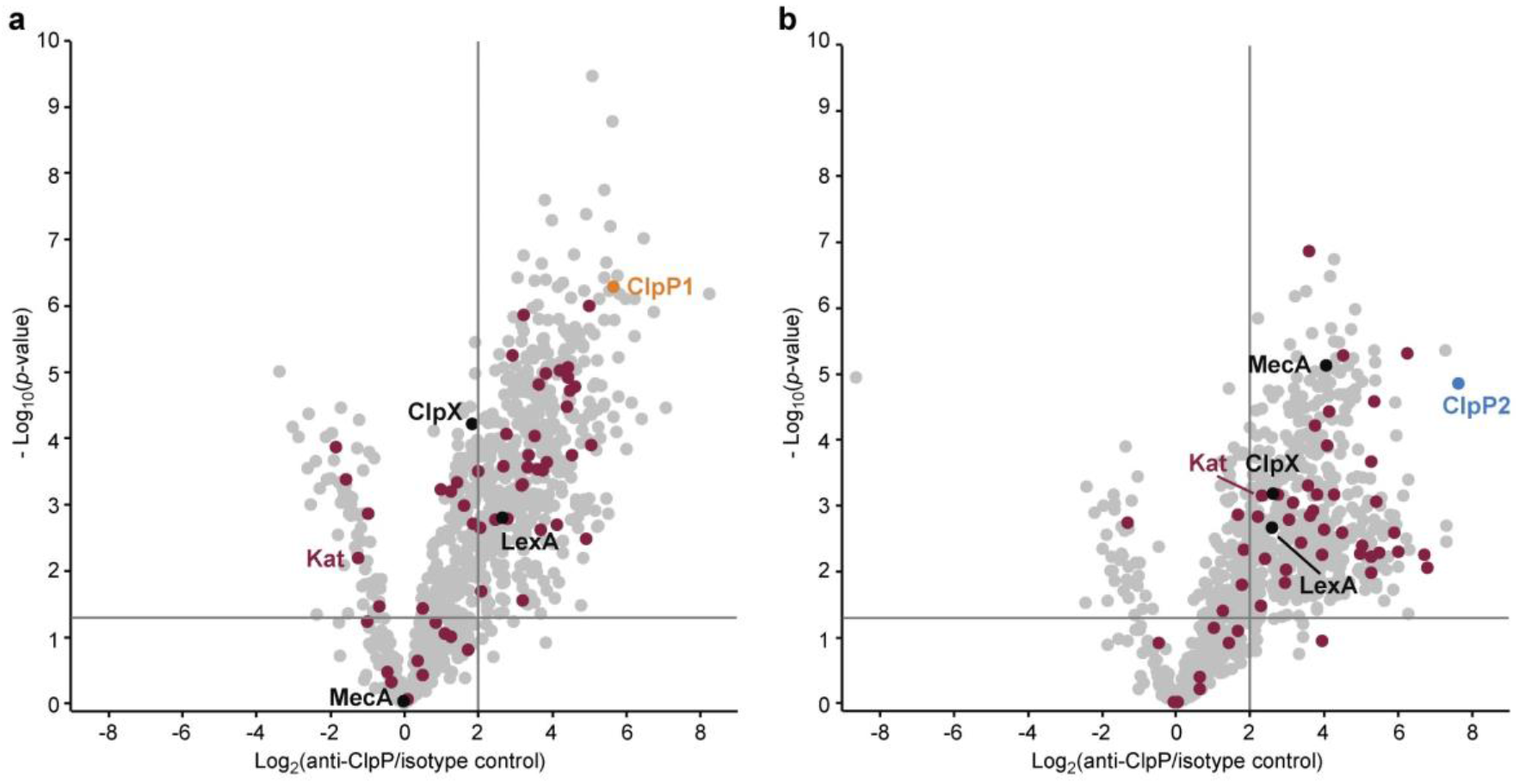
Co-immunoprecipitation of ClpP1 and ClpP2 in *L. monocytogenes* Δ*clpP* mutants. **a, b** Volcano plots of co-IPs with anti-ClpP antibody in *L. monocytogenes* Δ*clpP2* (**a**) and Δ*clpP1* (**b**) at stationary phase (37 °C). − Log_10_ *p*-values from two-sample Student’s *t*-test are plotted against log_2_ ratios of LFQ protein intensities. The vertical grey lines show 4-fold enrichment, the horizontal grey lines show −log_10_ t-test *p*-value *=* 1.3 (n = 4). Oxidoreductases are highlighted with purple. ClpP1 and ClpP2 are shown in orange and blue respectively.

In case of *clpP* deletion, substrates are expected to accumulate. In order to decipher these putative substrates, we searched for common hits between upregulated proteins in Δ*clpP* strains and the co-IPs (Figure 8a). Vice versa, proteins that were only enriched in the co-IP and not in the Δ*clpP* strains were regarded as putative interaction partners. We first evaluated the 37 °C results and then compared these with the 42 °C datasets.

**Figure 8.**
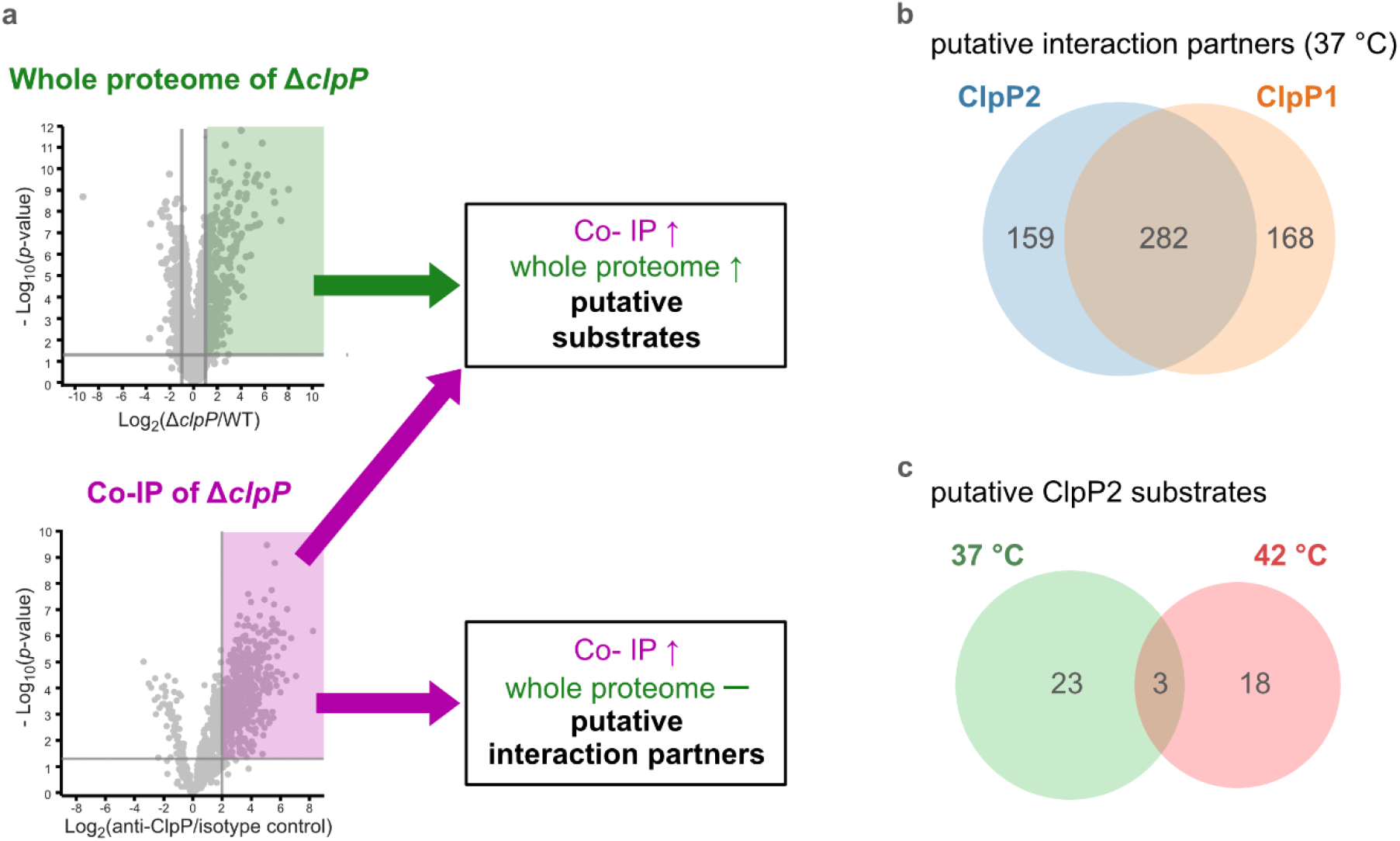
Proteomic analysis of the cellular functions of the ClpP isoforms and identification of putative substrates. **a** Proteins were classified as putative ClpP substrates (see table 1) if they were significantly enriched both in the whole-proteome analysis at 37 °C and/or 42 °C and in the anti-SaClpP co-IP of the respective Δ*clpP* mutants at the same temperature. Proteins that were significantly enriched only in the co-IP were classified as putative interaction partners of ClpP (Table S11, S12). **b** Venn-diagram showing the putative interaction partners of ClpP1 and ClpP2 at 37 °C. **c** Venn-diagram showing the putative substrates of ClpP2 at both temperatures.

Most of the putative interaction partners at 37 °C are common for both proteases (Figure 8b, Table S11). In general, many cellular metabolic terms, including amino acid and nucleobase-related metabolism, are upregulated in this protein group, suggesting an important role of ClpP in the general cellular metabolism.

ClpX was identified as a specific interaction partner of solely ClpP2 which is corroborated by previous structural and activity data demonstrating that ClpP1 lacks the hydrophobic binding pockets needed for association with chaperones (Dahmen et al., 2015; Gatsogiannis et al., 2019). The large number of putative interactors (450 for ClpP1 and 441 for ClpP2) emphasizes that also unspecific binders are among these proteins.

Applying the search criteria (enriched in XL-co-IP as well as in the corresponding deletion strain) we identified 26 putative ClpP2 substrate proteins at 37 °C (Table 1). Among these, analogs of four proteins (MecA, LexA, MurC and catalase) are known ClpP substrates (Feng et al., 2013; Flynn et al., 2003).

**Table 1.** List of putative ClpP2 substrates. *The functions of not annotated proteins were derived from BLAST searches.

| Gene | Uniprot ID | Description |
| --- | --- | --- |
| **Putative substrates at 37 °C** |  |  |
| lmo0485 | Q8Y9P0 | Putative oxidoreductase, iron response* |
| lmo0487 | Q8Y9N8 | Putative hydrolase* |
| lmo0582 (*iap*) | P21171 | Invasion-associated protein p60 |
| lmo0640 | Q8Y993 | Putative oxidoreductase* |
| lmo0823 | Q8Y8S1 | Putative oxidoreductase* |
| lmo0930 | Q8Y8H4 | Putative lactamase* |
| lmo1320 (*polC*) | Q8Y7G1 | PolC-type DNA polymerase III |
| lmo1350 (*gcvPB*) | Q8Y7D3 | Probable glycine dehydrogenase (decarboxylating) subunit 2 |
| lmo1381 (*acyP*) | Q8Y7A7 | Acylphosphatase (pyruvate metabolism) |
| lmo1406 (*pflB*) | Q8Y786 | Pyruvate formate-lyase (pyruvate metabolism) |
| lmo1515 | Q8Y711 | Similar to CymR cysteine metabolism repressor* |
| lmo1538 (*glpK*) | Q8Y6Z2 | Glycerol kinase (glycerol metabolism) |
| lmo1605 (*murC*) | Q8Y6S8 | UDP-N-acetylmuramate-L-alanine ligase |
| lmo1921 | Q8Y5Y2 | Unknown function |
| lmo1932 | Q8Y5X2 | Putative heptaprenyl diphosphate synthase (menaquinone biosynthesis)* |

**Table 1** continued
| Gene | Uniprot ID | Description |
| --- | --- | --- |
| Imo2168 | Q8Y5A1 | Putative lactoylglutathione lyase* |
| Imo2190 (*mecA*) | Q9RGW9 | ClpC adapter protein MecA |
| Imo2205 (*gpmA*) | Q8Y571 | 2,3-Bisphosphoglycerate-dependent phosphoglycerate mutase (glycolysis) |
| Imo2743 (*tal1*) | Q8Y3T8 | Probable transaldolase 1 (pentose phosphate pathway) |
| Imo2755 | Q8Y3S6 | Putative dipeptidyl-peptidase activity* |
| Imo2759 | Q8Y3S3 | Macro domain-containing protein (putative ADP-ribose binding) |
| Imo2785 (*kat*) | Q8Y3P9 | Catalase (H<sub>2</sub>O<sub>2</sub> detoxification) |
| Imo2829 | Q8Y3K6 | Putative nitroreductase* |
| **Putative substrates at both temperatures** |  |  |
| Imo1302 (*lexA*) | Q8Y7H7 | LexA SOS response repressor |
| Imo2182 | Q8Y587 | Putative ferrichrome ABC transporter ATP-binding protein* |
| Imo2526 (*murA*) | Q8Y4C4 | UDP-N-acetylglucosamine 1-carboxyvinyltransferase 1 |
| **Putative substrates at 42 °C** |  |  |
| Imo0227 | Q8YAB9 | tRNA-dihydrouridine synthase |
| Imo0229 (*ctsR*) | Q7AP89 | CtsR (transcription repressor of class III heat shock genes) |
| Imo0231 (*mcsB*) | Q48759 | Arginine Kinase McsB |
| Imo0454 | Q8Y9R9 | Putative MoxR family ATPase* |
| Imo0608 | Q8Y9C4 | Putative multidrug ABC transporter * |
| Imo0785 | Q8Y8V7 | Transcriptional Regulator ManR |
| Imo1293 (*glpD*) | Q8Y7I4 | Glycerol-3-phosphate dehydrogenase |
| Imo1387 | Q8Y7A2 | Putative pyrrolysine-5-carboxylate reductase* |
| Imo1475 (*hrcA*) | P0DJM4 | HrcA (heat-inducible transcription repressor A) |
| Imo1631 (*trpD*) | Q8Y6Q3 | Anthranilate phosphoribosyltransferase |
| Imo1713 (*mreB*) | Q8Y6H3 | Cell shape-determining protein MreB |
| Imo1813 | Q8Y684 | L-serine deaminase |
| Imo1881 | Q8Y621 | Putative 5'-3'-exonuclease* |
| Imo2267 (*addA*) | Q8Y511 | ATP-dependent helicase/nuclease subunit A |
| Imo2352 | Q8Y4T0 | Putative LysR family transcriptional regulator* |
| Imo2489 (*uvrB*) | Q8Y4F5 | UvrABC system protein B, excision nuclease |
| Imo2552 (*murZ*) | Q8Y4A2 | UDP-N-acetylglucosamine 1-carboxyvinyltransferase 2 |
| Imo2712 | Q8Y3W7 | Putative gluconate kinase (Pentose phosphate pathway)* |

Six of the 26 classified substrates are oxidoreductases and four other proteins are associated with oxidative stress (LexA, Lmo1515 CymR analog, Lmo2168 putative lactoylglutathione lyase and Lmo2182 ferrichrome ABC transporter) suggesting that ClpP plays a crucial role in redox homeostasis in *L. monocytogenes*, similar to *S. aureus* ClpP (Farrand et al., 2015; Michel et al., 2006). For example, LexA, the repressor of the SOS regulon, is a known ClpP target in *E. coli* and in *S. aureus* (Cohn et al., 2011; Flynn et al., 2003). During the activation of the SOS response, LexA undergoes autocleavage and the N- and C-terminal domains are separated (Michel, 2005). Consequently, the ClpX recognition sequence gets exposed and NTD (in some organisms also the CTD) is degraded by ClpXP (Cohn et al., 2011). While we were unable to detect any peptides that stretch across the autocleavage site, the fact that many SOS response proteins were upregulated in both Δ*clpP2* and Δ*clpP1/2* suggests that cleaved LexA accumulates, which can only weakly bind to the SOS box.

Interestingly, the number of overall and enriched proteins identified via XL-co-IP largely dropped at 42 °C (Figure S5, Table S12). Of the putative interaction partners the majority was identified for ClpP1 (123 for ClpP2 and 230 for ClpP1) and fewer were mutual for both isoforms (Figure S5c). A large fraction in this protein group again falls into general metabolic pathways. Yet, some minor differences such as DNA replication being overrepresented are notable.

In search for ClpP2 substrates at this temperature, we identified 21 putative proteins (Table 1). Interestingly, only three of those proteins are also substrate candidates at 37 °C (LexA, MurA, Lmo2182 ferrichrome ABC transporter) with 18 additional proteins being substrate candidates solely at 42 °C (Figure 8c). This indicates that the substrate scope is adapted with changing conditions like temperature. There are five additional proteins among those that have been previously described as ClpP substrates in other bacteria, namely the two heat transcriptional regulators HrcA and CtsR, ClpC adaptor protein McsB, DNA damage repair protein UvrB and glycerol-3-phosphate dehydrogenase GlpD (Feng et al., 2013; Flynn et al., 2003). HrcA and CtsR regulate the expression of class I and III heat shock proteins, while UvrB is an integral part of the SOS response (Kisker et al., 2013; Nair et al., 2000; Schulz & Schumann, 1996; van der Veen et al., 2010). In addition, we identified AddAB helicase/nuclease subunit A as putative ClpP2 substrate, which is also an SOS response protein (van der Veen et al., 2010). Other identified putative substrates at 42 °C are involved in cell wall synthesis, cell shape (MurA, MurZ, MreB) and metabolic processes (e.g. GlpD, Lmo1387 pyrrolysin-5-carboxylate reductase, TrpD, Lmo1813 L-serine deaminase, Lmo2712 gluconate kinase). Thus, in summary 44 putative ClpP2 substrates were identified in this work. At both temperatures, nearly no putative ClpP1 substrates could be identified with this approach in our datasets (except for FhuC at 42 °C). This lack of substrates was expected as ClpP1 is only active in complex with ClpP2.

With many identified dysregulated proteins and putative substrates being related to oxidative stress, we finally investigated the ability of the Δ*clpP* mutants to grow in medium supplemented with H_2_O_2_ (Figure 9). Surprisingly, only Δ*clpP1/2* could grow in the presence of 100 ppm H_2_O_2_. This is in line with the observation that SOS-related GOBP terms were significantly upregulated only in Δ*clpP1/2* but not in Δ*clpP2* and Δ*clpP1 at 37 °C.* Thus, this indicates that a constitutively upregulated SOS response system readily protects cells from H_2_O_2_ in the Δ*clpP1/2* strain.

**Figure 9.**
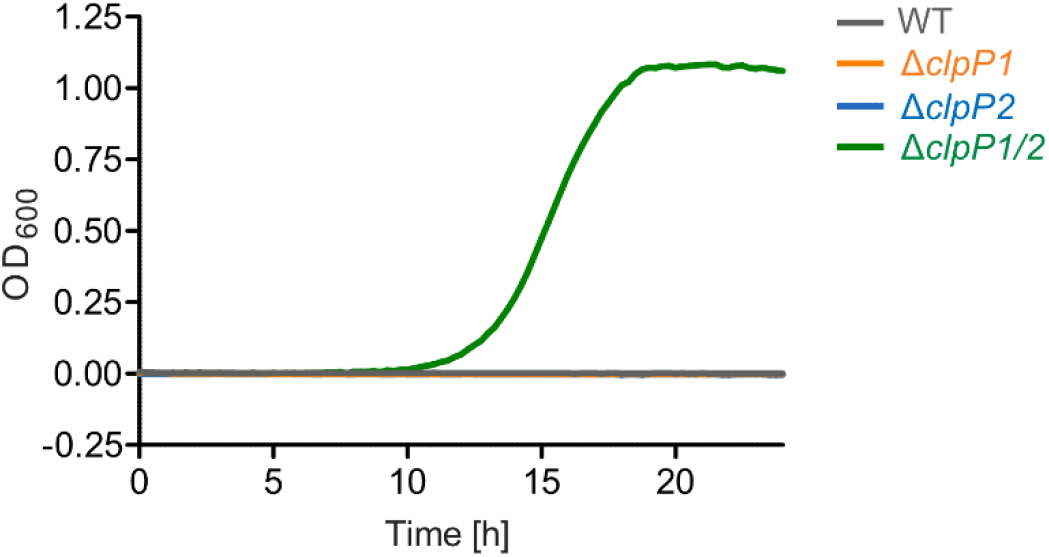
*L. monocytogenes* Δ*clpP1/2* is resistant against oxidative stress. Growth curves of the Δ*clpP* mutants in the presence of 100 ppm H_2_O_2_ (BHI medium, 37 °C). Note that the WT strain and the single *clpP* knockouts show no growth under these conditions. Means of triplicates are shown. The experiment was independently repeated with qualitatively identical results (data not shown).

## Discussion

ClpP is a conserved heat shock protein in bacteria and in eukaryotic organelles. Some organisms have more than one *clpP* gene, but the role of multiple ClpPs in these organisms is not well understood. In bacteria, it is known that two different ClpPs are able to form heterocomplexes to become active, tune the cleavage specificity or enhance the activity of the homocomplexes (Dahmen et al., 2015; Li et al., 2016; Mawla et al., 2020; Pan et al., 2019). Here we examined the biological role of ClpP1/2 heterocomplex formation in *L. monocytogenes* and the specific physiological functions of both ClpPs.

We showed that ClpP1 and ClpP2 do not bind to each other at 10 °C, and under these conditions ClpP2 is a homotetradecamer and ClpP1 an inactive heptamer. At high temperatures, especially above 37 °C, the ClpP1/2 heterocomplex is formed displaying enhanced substrate turnover. We suspected that this trait is important for modulation of ClpP proteolytic activity and is therefore crucial for stress response and virulence regulation. In order to study this effect in intact *L. monocytogenes* cells, we performed MS-based co-IP experiments at various temperatures. We observed enhanced ClpP1 binding to the bait ClpP2 at 42 °C as compared to 20 °C, which indeed indicates that temperature affects intracellular heterooligomer formation. However, the analysis of ClpP1 as bait was challenged due to its low abundance at 20 °C. Thus, further research is needed to investigate the exact conditions under which heterooligomerization takes place *in situ* and elucidate whether other factors such as binding partners or post-translational modifications can modulate ClpP1/2 complex formation.

With the aim of dissecting the physiological functions of each ClpP isoforms, we constructed single and double *clpP* deletion mutants in *L. monocytogenes* EGD-e. Phenotypic assays showed decreased growth of Δ*clpP1/2* in culture medium and in macrophages. MS-based whole proteome analysis demonstrated that the deletion of *clpP1* only caused minimal changes in the proteome while Δ*clpP2* and Δ*clpP1/2* mutants differed greatly from the wild type. These results highlight the predominant role of ClpP1 as an enhancer of catalytic turnover which is unable to process substrates by itself. In Δ*clpP2* and Δ*clpP1/2* mutants, class III heat shock proteins and a subset of the SOS response proteins as well as iron-containing proteins were upregulated. These results suggest that ClpP plays an important role in the regulation of oxidative stress response, which is in line with the results of transcriptomic analysis of the *S. aureus* Δ*clpP* mutant (Michel et al., 2006). Furthermore, the upregulated SOS response predominantly observed in the Δ*clpP1/2* mutant led to a strong H_2_O_2_ resistance for this strain.

We conducted co-IPs in the single mutants with anti-ClpP antibody in order to identify specific ClpP1 and ClpP2 substrates and interaction partners. Combined analysis of the co-IP and whole proteome data at two temperatures led to the identification of 44 putative ClpP2 substrates and ~700 putative ClpP1 and ClpP2 interaction partners. To a large extent the putative interaction partners are shared between ClpP1 and ClpP2 and many of them are involved in the general cellular metabolism, including amino acid and nucleic acid metabolism. ClpP might have an indirect effect on proteostasis via the interactions with these proteins. A large fraction of the identified ClpP2 substrates are related to transcriptional regulation, cell wall synthesis, cellular metabolism and oxidative stress, including LexA, corroborating the upregulation of SOS response proteins in the whole proteome.

In summary, we found that the ClpP1/2 heterocomplex in *L. monocytogenes* acts as a thermometer which assembles at elevated temperatures and revealed ClpP’s role in coping with heat-induced stress. Studying ClpP heterocomplex formation in other organisms under varying conditions might reveal that thermosensitivity is a general feature of ClpPs in bacteria carrying more than one *clpP* genes. This study and initial data from *M. tuberculosis* (Leodolter et al., 2015), showing a temperature-dependent, reversible assembly of a ClpP1/2 heterocomplex without an activator peptide, point in this direction.

## Acknowledgements

This work was performed within the framework of SFB 1035 (German Research Foundation DFG, Sonderforschungsbereich 1035, Projektnummer 201302640, project A09). We thank Mona Wolff and Katja Bäuml for technical support as well as Dr. Stuart Ruddell for critical proofreading of the manuscript. We are grateful to Prof. Dr. Thilo M. Fuchs and Dr. Jakob Schardt for their help with cloning in *L. monocytogenes*. We thank Dr. David Lyon (UZH, Institute of Molecular Life Sciences, Switzerland) for support concerning the aGOtool.

## Methods

### Protein overexpression and purification

ClpP2 was obtained as described previously (Zeiler et al., 2013). In short, expression constructs with C-terminal Strep-tag II were cloned in pET301 plasmids, over-expressed in *E. coli* BL21(DE3) and purified by affinity chromatography and gel filtration. ClpP1 was kindly provided by Dr. Maria Dahmen (Dahmen et al., 2015). Co-expressed ClpP1/2 was obtained as described previously (Gatsogiannis et al., 2019). Creatine kinase (10 127 566 001) was purchased from Roche (Roche Diagnostics GmbH, Mannheim, Germany).

### Analytical size-exclusion chromatography followed by intact protein mass spectrometry

544 nmol ClpP1_7_ (1:1 ClpP1:ClpP2 monomeric ratio) and/or 272 nmol ClpP2_14_ were incubated for 30 min at the indicated temperatures (0 – 48 °C) in ClpP-GF buffer (20 mM MOPS, 100 mM KCl, 5% glycerol, pH 7.0) in a final volume of 100 μL. The samples were loaded on a pre-equilibrated Superdex 200 10/300 gel filtration column (GE Healthcare, Chicago, United States) connected to an ÄKTA Purifier 10 system (GE Healthcare) and eluted with 1 CV ClpP-GF buffer. 200 μL fractions were collected. UV absorption was recorded at 280 nm. The oligomerization state was determined by comparison of the elution volumes to the calibration curve of the column (Gel Filtration Calibration Kit, GE Healthcare). The fraction corresponding to the tetradecamer peak was analyzed by intact protein mass spectrometry. Measurements were carried out on a Dionex Ultimate 3000 HPLC system (Thermo Fisher Scientific, Waltham, United States) coupled to a Thermo LTQ-FT Ultra mass-spectrometer (Thermo Fisher Scientific) with electrospray ionisation source (spray voltage 4.0 kV, tube lens 110 V, capillary voltage 48 V, sheath gas 60 a.u., aux gas 10 a.u., sweep gas 0.2 a.u.). 5 μL were desalted with a MassPREP desalting cartridge (Waters, Milford, United States). The mass spectrometer was operated in positive mode collecting full scans at high resolution (R =200 000) from m/z =600 to m/z =2000. Collected data was deconvoluted using the Thermo Xcalibur Xtract algorithm (Thermo Fisher Scientific).

The experiments with a mixture of ClpP1_7_ and ClpP2_14_ at 20 °C and at 42 °C were repeated with qualitatively identical results. Plots were made with Microcal OriginPro 2018 (OriginLab Corporation, Northampton, United States).

### Protease assay

Protease assays were carried out in flat bottom black 96-well plates in a final volume of 60 μL. 0.1 μM ClpP2_14_ or a mixture of 0.2 μM ClpP1_7_ and 0.1 μM ClpP2_14_ (1:1 ClpP1:ClpP2 monomeric ratio), ClpX_6_ (0.4 μM) and ATP regeneration mix (4 mM ATP, 16 mM creatine phosphate, 20 U/mL creatine kinase) were pre-incubated for 30 min at the indicated temperatures (30 °C, 37 °C and 42 °C) in PZ buffer (25 mM HEPES, 200 mM KCl, 5 mM MgCl_2_, 1 mM DTT, 10% glycerol, pH 7.6). 0.4 μM eGFP-LmSsrA substrate was added and fluorescence (λ_ex_ = 485 nm, λ_em_ = 535 nm) was measured at the respective temperatures with an infinite M200Pro plate reader (Tecan, Männedorf, Switzerland). Data were recorded in triplicates. The measurements were independently repeated with qualitatively identical results. Protease activity was determined by linear regression using Microsoft Excel and plots were made with GraphPad Prism 6 (GraphPad, San Diego, United States).

### Cloning of *L. monocytogenes* mutants

#### Generation of *L. monocytogenes clpP1(191)*::2×myc and *L. monocytogenes clpP2(199)*:: 2×myc

##### Construction of pLSV101_clpP-2×myc shuttle vectors

Ca. 1000 base pairs upstream and downstream from the C-terminus of *clpP1* were amplified by PCR using the A–B and C–D primer pairs from Table 2 (Phusion polymerase, GC buffer, New England Biolabs, Ipswich, United States). For *clpP2*, ca. 700 bp upstream and downstream were amplified using the A–B and C–CD primer pairs from Table 2 (Phusion polymerase, GC buffer, New England Biolabs). The 2×myc tag was added to the B primers as overhangs. The PCR products were purified with E.Z.N.A. Cycle Pure Kit (Omega Bio-tek, Norcross, United States). The AB fragments were digested with SalI-HF (New England Biolabs) and BglII (Promega, Madison, United States), the CD fragments were digested with BglII (Promega) and BamHI (New England Biolabs) and the empty pLSV101 vector was digested with SalI-HF and BamHI-HF (New England Biolabs). The digested DNAs were purified with E.Z.N.A. Gel Extraction Kit (Omega Bio-tek) after agarose gel electrophoresis. The AB and CD fragments were ligated with T4 DNA ligase (New England Biolabs) (1:1 molar ratio, 15 °C, overnight). The ligated fragments were amplified by PCR (Phusion polymerase, HF buffer, New England Biolabs) using the clpP1_A–clpP1D and clpP2_A– clpP2_CD primer pairs (Table 2). The PCR products were purified with E.Z.N.A. Gel Extraction Kit (Omega Bio-tek) after agarose gel electrophoresis. The ABCD fragments were digested with SalI-HF and BamHI-HF (New England Biolabs) and dephosphorylated with Antarctic phosphatase (New England Biolabs). The fragments were purified with E.Z.N.A. Gel Extraction Kit (Omega Bio-tek) after agarose gel electrophoresis. The fragments were ligated into the pLSV101 vector (1:1 and 3:1 molar ratios) with T4 DNA ligase (New England Biolabs) (10 °C for 30 s and 30 °C for 30 s alternating overnight). The ligated vectors were transformed into chemically competent *E. coli* TOP10. *E. coli* containing pLSV101 was grown with 200 μg/mL erythromycin. Colonies were tested with colony PCR using pLSV101_seq fwd and rev primers (Table 2). The vectors were purified from positive colonies with NucleoSpin Plasmid EasyPure, Mini kit (MACHEREY-NAGEL, Düren, Germany) (elution with ddH_2_O) and sequenced by Sanger sequencing with A and D primers.

**Table 2.**
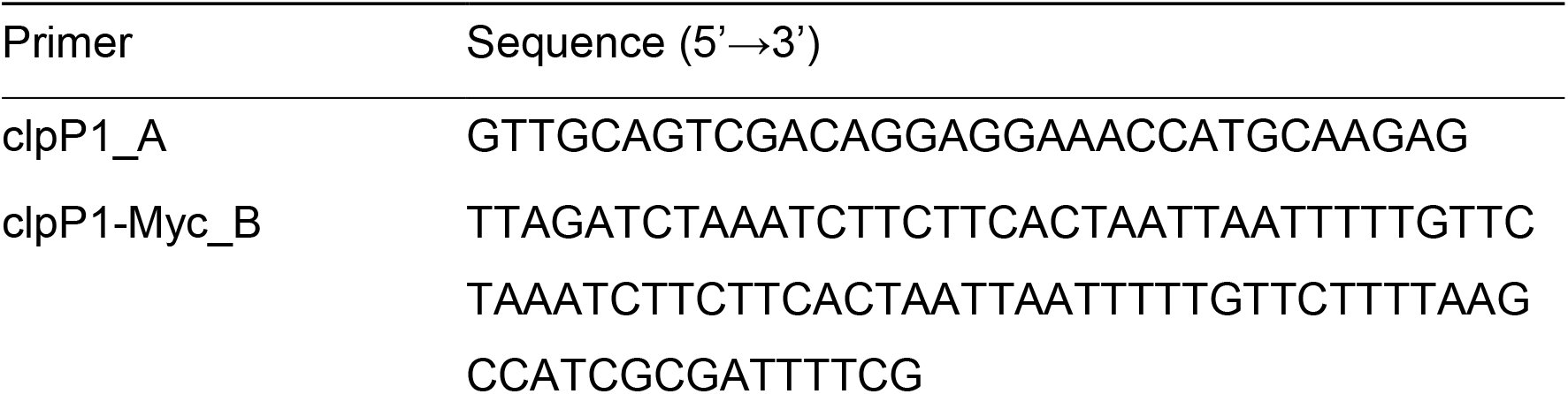
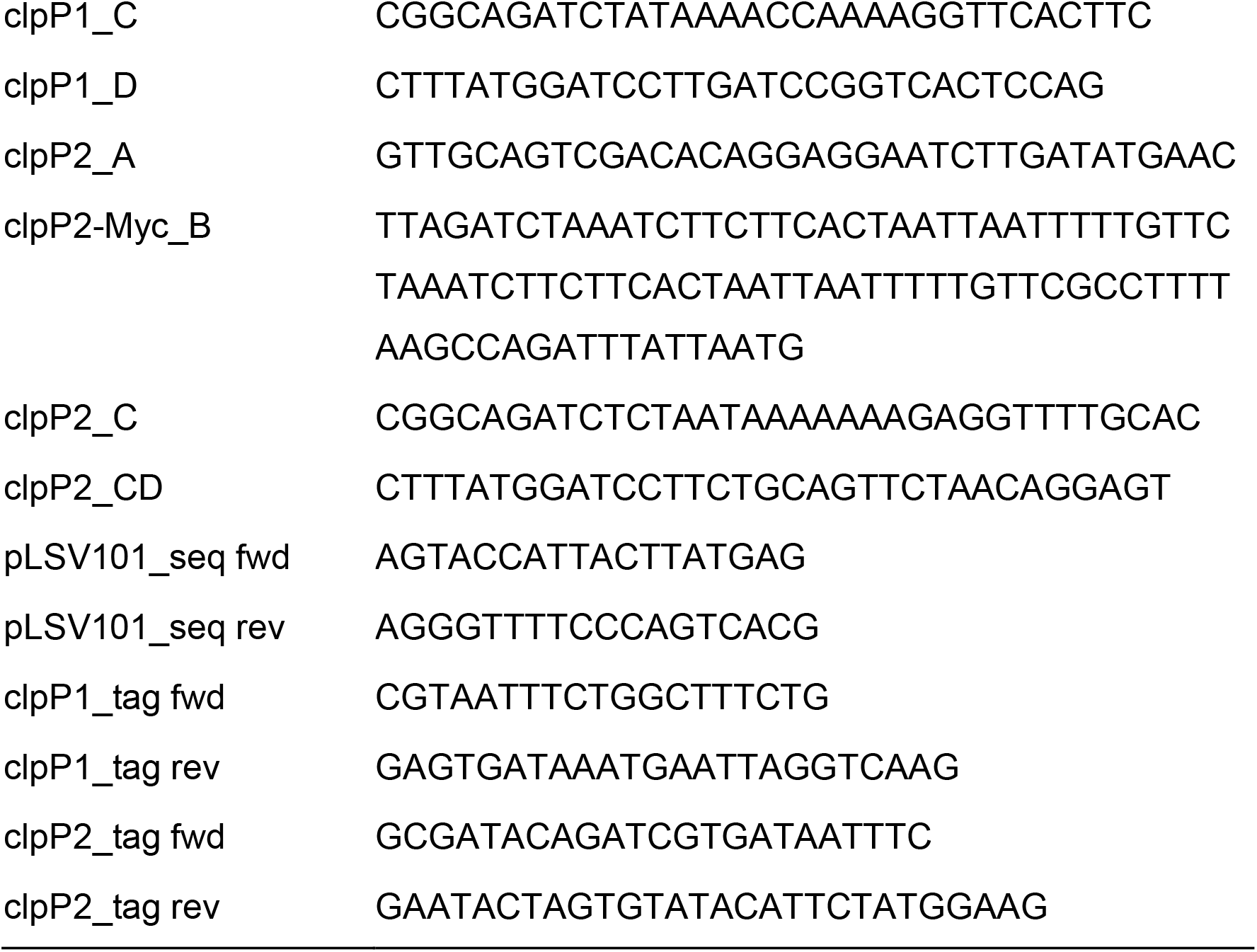
List of primers used for the genomic insertion of 2×myc tag into *L. monocytogenes*.

*Preparation of electrocompetent L. monocytogenes* 200 mL BHI medium (7.5 g/L brain infusion, 1 g/L peptone, 10 g/L heart infusion, 5 g/L NaCl, 2.5 g/L Na_2_HPO_4_, 2 g/L glucose, pH 7.4) was inoculated to an initial OD_600_ of 0.05 with an overnight culture of *L. monocytogenes* EGD-e. The culture was grown to OD_600_ = 0.5 at 37 °C, 200 rpm. 5 μg/mL penicillin G was added, and the bacteria were incubated at 37 °C, 200 rpm for 15 min and on ice without shaking for 10 min. The cells were harvested (4000 g, 10 min, 4 °C) and washed with 30 mL ice-cold SMHEM medium (952 mM saccharose, 3.5 mM MgCl_2_, 7 mM HEPES, pH 7.2). The pellet was resuspended in 2 mL cold SMHEM medium. 100 μL aliquots were prepared and shock-frozen in liquid N_2_ and stored at −80 °C.

##### Transformation into L. monocytogenes

Electrocompetent *L. monocytogenes* EGD-e aliquots were thawed on ice and 1 μg plasmid was added. The cells were transferred into ice-cold 2 mm electroporation cuvettes (Bio-Rad, Hercules, United States) and electroporated (2500 V, 200 Ω, 25 μF, exponential decay, time constant < 4 ms) using Gene Pulser Xcell (Bio-Rad). 1 mL BHI medium + 0.5 mM saccharose was added and the cells were incubated at 30 °C for 4 h and plated on BHI agar plates containing 10 μg/mL erythromycin. The plates were incubated at 30 °C for 3 days.

*Homologous recombination and colony selection* 2.5 mL BHI medium with 10 μg/mL erythromycin was inoculated with single colonies after transformation. 10^−2^ and 10^−6^ dilutions were plated on BHI + 10 μg/mL erythromycin agar plates and incubated at 42 °C for 2 days. Colony PCR (OneTaq polymerase, New England Biolabs) with the respective primer pairs clp_A–pLSV101_seq rev and pLSV101_seq fwd–clp_D (Table 2) was performed to check the genomic integration of the fragments. Positive colonies were subcultivated several times in 3 mL BHI medium without antibiotic at 30 °C (200 rpm). 10^−6^ dilutions were plated on BHI agar plates (37 °C, overnight). Single colonies were picked and transferred to BHI agar plates with and without 10 μg/mL erythromycin (37 °C, overnight). Erythromycin-sensitive strains were tested with colony PCR (OneTaq DNA polymerase, New England Biolabs) using the clpP_tag fwd and rev primer pair (Table 2) to check for integration of the 2×myc tag into the genome.

#### Generation of *L. monocytogenes* Δ*clpP1*

##### Construction of pMAD_ΔclpP1 shuttle vector

A pMAD shuttle vector derivative was used to introduce a deletion of *clpP1* (Arnaud et al., 2004). Approx. 1000 bp upstream (clpP1_KO_A and clpP1_KO_B, Table 3) region of clpP1 was amplified by PCR (GC buffer, Phusion polymerase, New England Biolabs) using isolated *L. monocytogenes* EGD-e DNA as template. The PCR product was purified (Cycle Pure Kit, E.Z.N.A., Omega Bio-tek) and digested with MluI and NcoI (Promega, standard protocol). pMAD plasmid was also digested with MluI and NcoI and dephosphorylated by addition of TSAP (Promega, streamlined restriction digestion protocol) for 20 min. After restriction digest products were purified (MicroElute DNA Clean-Up Kit, E.Z.N.A., Omega Bio-tek). Ligation into pMAD vector was conducted using T4 DNA Ligase (Promega, standard protocol) overnight at 8 °C and a vector:insert ratio of 1:6. The ligation product (pMAD-AB) was chemically transformed into *E. coli* TOP10 cells and plated onto LB agar containing ampicillin. Accordingly, a 1000 bp downstream (clpP1_KO_C and clpP1_KO_D, Table 3) region of clpP1 was amplified by PCR (GC buffer, Phusion polymerase, New England Biolabs) using isolated *L. monocytogenes* EGD-e DNA as template. The PCR product was purified (Cycle Pure Kit, E.Z.N.A., Omega Bio-tek) and digested with MluI and BamHI (Promega, standard protocol). pMAD-AB plasmid was also digested with MluI and BamHI and dephosphorylated by addition of TSAP (Promega, streamlined restriction digestion protocol) for 20 min. After restriction digest products were purified (MicroElute DNA Clean-Up Kit, E.Z.N.A., Omega Bio-tek). Ligation into pMAD-AB vector was conducted using T4 DNA Ligase (Promega, standard protocol) overnight at 8 °C and a vector:insert ratio of 1:6. Insertion of the desired construct was tested after plasmid extraction (Plasmid Mini Kit I, E.Z.N.A., Omega Bio-tek) by analytical restriction digest and sequencing (pMAD-seq-for and pMAD-seq-rev, Table 3).

**Table 3.**
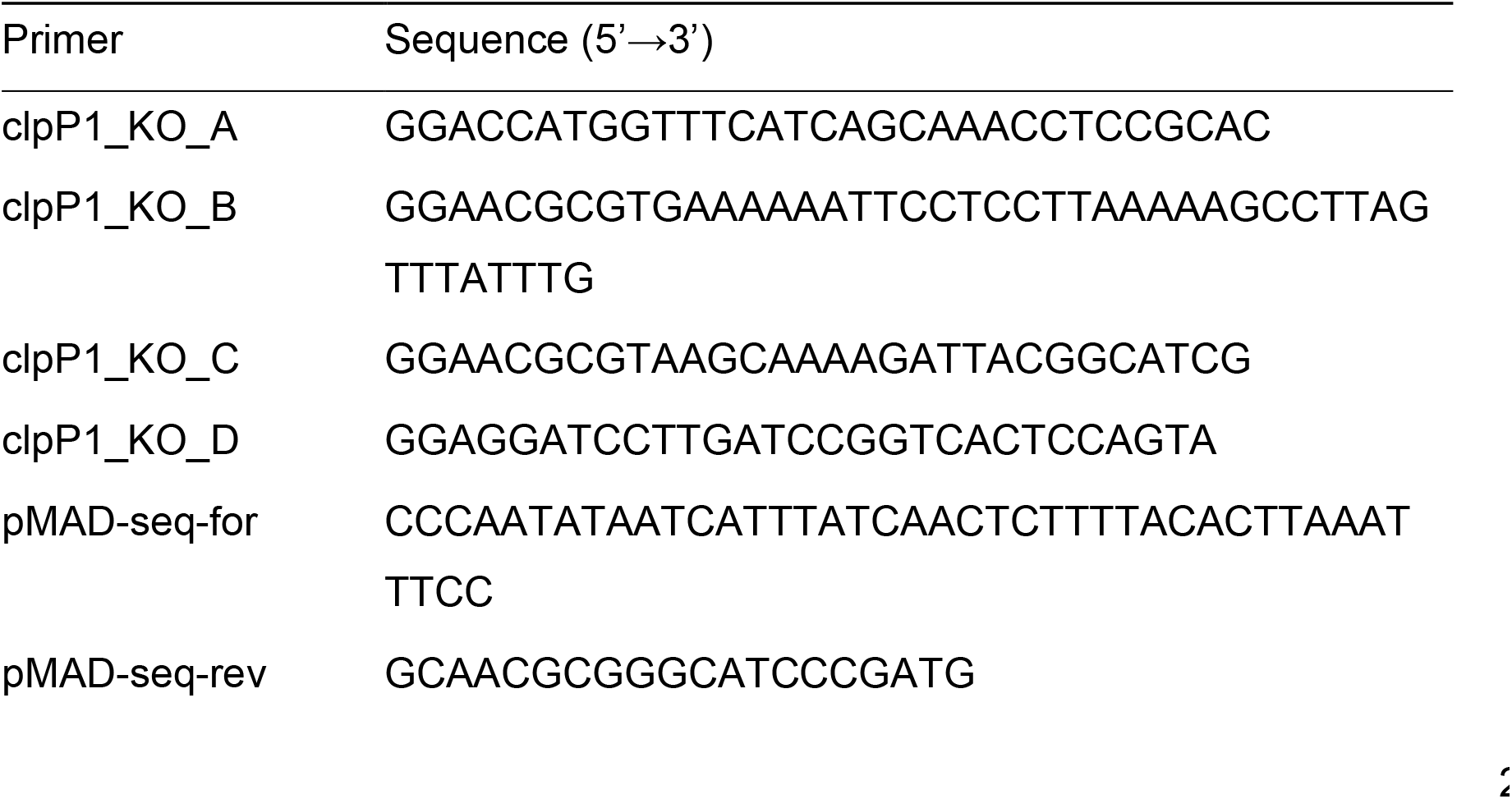
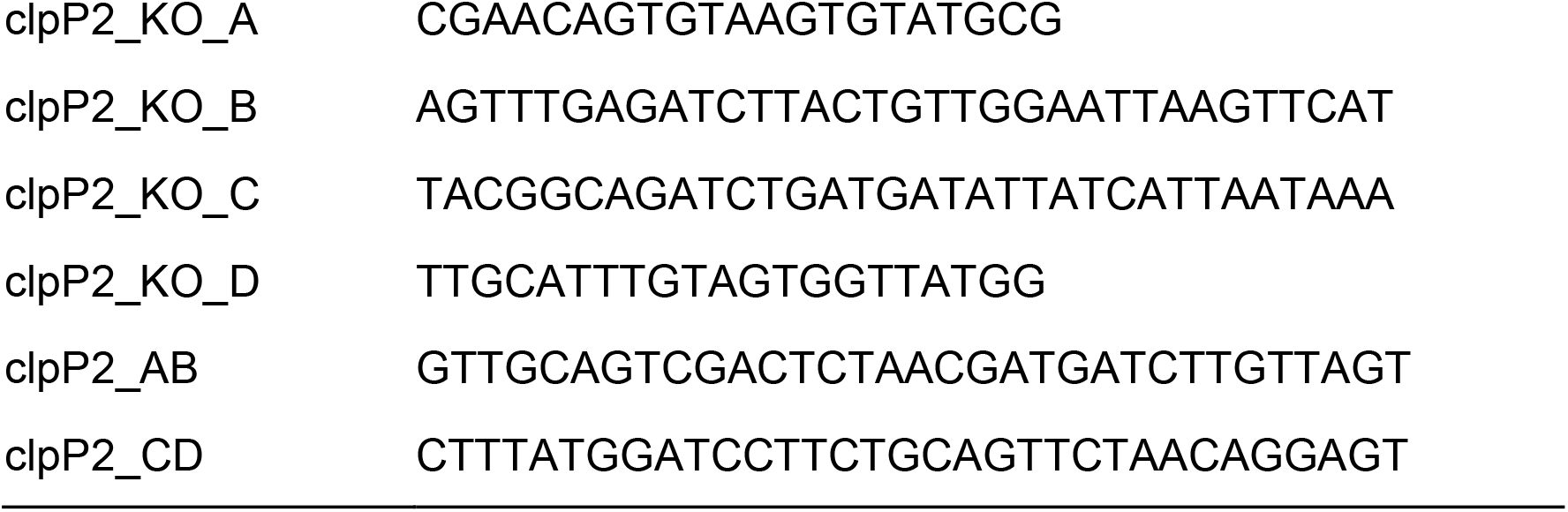
List of primers used for the construction *L. monocytogenes clpP* deletion mutants.

*Preparation of electrocompetent L. monocytogenes* 100 mL of BM medium (10 g/L soy peptone, 5 g/L yeast extract, 5 g/L NaCl, 1 g K_2_HPO_4_ × 3 H_2_O, 1 g/L glucose, pH 7.4–7.6) were inoculated with 1 mL (1:100) from a *L. monocytogenes* EGD-e overnight culture and incubated at 37 °C until an OD_600_ of 0.5 was reached. Cells were centrifuged (5000 g, 15 min, 4 °C) and washed three times with cold 10% glycerol (sterile): 1.) 100 mL; 2.) 50 mL; 3.) 25 mL. The pellet was resuspended in 400 μL cold 10% glycerol and 75 μL aliquots were frozen in liquid nitrogen and stored at −80 °C.

##### Transformation into L. monocytogenes

Electrocompetent *L. monocytogenes* was thawed at room temperature (RT) and incubated for 10 min with > 1 μg plasmid. The suspension was transferred into a 0.1 cm electroporation cuvette (Bio-Rad) and electroporated (exponential, 25 μF, 1 kV, 400 Ω) using a Gene Pulser Xcell (Bio-Rad). Immediately after the pulse 1 mL pre-warmed BM medium was added and incubated at 30 °C for 90 min. The cell suspension was streaked onto BM agar containing selective antibiotic + X-gal and incubated until colonies were visible.

##### Selection protocol – pMAD

After successful transformation into *L. monocytogenes* EGD-e, indicated by blue colonies, single colonies were picked and incubated overnight at 30 °C in the presence of 1 μg/mL erythromycin. 10 mL BM medium were inoculated 1:1000 from the overnight culture and incubated 2 h at 30 °C and 6 h at 42 °C. 100 μL diluted cultures (10^−2^ to 10^−6^) were plated onto BM agar (containing 1 μg/mL erythromycin and 100 μg/mL X-gal) and incubated at 42 °C until colonies with blue coloration were visible (enrichment of single crossover). Ten light blue colonies were picked and incubated (together) in 10 mL BM medium at 3 °C for 8 h followed by overnight incubation at 42 °C. 10 mL BM medium were inoculated 1:1000 from this overnight culture and grown for 4 h at 30 °C and additional 4 h at 42 °C. 100 μL of diluted cultures (10^−2^ to 10^−6^) were plated onto BM agar containing X-gal and incubated at 4 °C. White colonies were picked and streaked onto BM agar containing erythromycin and X-gal and onto BM agar containing only X-gal. Plates were incubated at 30 °C and erythromycin susceptible colonies further analyzed by colony PCR followed by analytical restriction digest and sequencing. For colony PCR small parts of colonies were resuspended in 50 μL sterile water and 1 μL thereof was used in PCR reactions with an initial denaturation step for 10 min (95 °C).

#### Generation of *L. monocytogenes* Δ*clpP2* andΔ*clpP1/2*

##### Construction of pLSV101_ΔclpP2 shuttle vector

A construct derived from the mutagenesis vector pLSV101 was used for *clpP2* deletion (pLSV101 was kindly provided by Prof. Dr. Thilo M. Fuchs) (Joseph et al., 2006). Ca. 1000 base pairs upstream and downstream from the *clpP2* gene were amplified by PCR using the A–B and C–D primer pairs from table 3 (Phusion polymerase, GC buffer, New England Biolabs). The PCR products were purified with E.Z.N.A. Gel Extraction Kit (Omega Bio-tek) after agarose gel electrophoresis. The fragments were digested with BglII (Promega) and purified with E.Z.N.A. Cycle Pure Kit (Omega Bio-tek). The AB and CD fragments were ligated with T4 DNA ligase (New England Biolabs) (1:1 molar ratio, 15 °C, overnight). The ligated fragment was amplified by PCR (Phusion polymerase, HF buffer) using the AB–CD primer pair (Table 3). The PCR product was purified with E.Z.N.A. Cycle Pure Kit (Omega Bio-tek). The insert and the empty pLSV101 vector were digested with SalI-HF and BamHI-HF (New England Biolabs) and purified with E.Z.N.A. Gel Extraction Kit (Omega Bio-tek) after agarose gel electrophoresis. The fragment was ligated into the pLSV101 vector (3:1 molar ratio) with T4 DNA ligase (New England Biolabs) (16 °C overnight). The ligated vector was transformed into chemically competent *E. coli* TOP10. *E. coli* containing pLSV101_ΔclpP2 was grown with 300 μg/mL erythromycin. Colonies were tested with colony PCR using pLSV101_seq fwd and rev primers (Table 2). The vectors were purified with E.Z.N.A. Plasmid Mini Kit I (Omega Bio-tek) from positive colonies (elution with ddH_2_O) and sequenced by Sanger sequencing with pLSV101_seq fwd and rev primers.

##### Transformation into L. monocytogenes

Electrocompetent *L. monocytogenes* EGD-e and Δ*clpP1* cells were prepared as described above. Aliquots of electrocompetent cells were thawed on ice and 2 or 5 μg plasmid was added. The cells were transferred into ice-cold 2 mm electroporation cuvettes (Bio-Rad) and electroporated (2500 V, 200 Ω, 25 μF, exponential decay, time constant ~ 4 ms) using Gene Pulser Xcell (Bio-Rad). 1 mL warm BHI medium was added and the cells were incubated at 30 °C for 6 h under shaking at 200 rpm and plated on BHI agar plates with 10 μg/mL erythromycin. The plates were incubated at 30 °C for 5 days.

*Homologous recombination and colony selection* 2.5 mL BHI medium with 10 μg/mL erythromycin was inoculated with single colonies after transformation. 10^−2^ and 10^−5^ dilutions were plated on BHI + 10 μg/mL erythromycin agar plates and incubated at 42 °C for 2 days. Colony PCR (OneTaq polymerase, New England Biolabs) with the primer pairs clp2_KO_A–pLSV101_seq rev and pLSV101_seq fwd–clpP2_KO_D (see Tables 2 and 3) was performed to check the genomic integration of the fragments. Positive colonies were subcultivated several times in 2.5 mL BHI medium without antibiotic at 30 °C (200 rpm). 10^−6^ dilutions were plated on BHI agar plates (RT, 3 days). Single colonies were picked and transferred to BHI agar plates with and without 10 μg/mL erythromycin (37 °C, overnight). Erythromycin-sensitive strains were tested with colony PCR (OneTaq DNA polymerase, New England Biolabs) using the clpP2_KO_A and clpP2_KO_D primer pair (Table 3) to check for *clpP2* deletion.

#### Western blot

5 mL BHI medium was inoculated with *L. monocytogenes* EGD-e, Δ*clpP1*, Δ*clpP2* and Δ*clpP1/2* strains. An amount of cells corresponding to 200 μL of OD_600_ = 20 of each mutant was harvested (4000 g, 10 min, 4 °C). The cells were lysed by ultrasonication (3×20 s, 75%, cooled on ice during breaks). 2× Laemmli buffer was added and 20 μL sample was separated by SDS-PAGE (12.5% polyacrylamide, 150 V, 2.5 h). The proteins from the polyacrylamide gel were transferred to a methanol-soaked PVDF membrane (Bio-Rad) in a Trans-Blot SD semi-dry western blot cell (Bio-Rad) using blotting buffer (48 mM Trizma, 39 mM glycine, 0.04% SDS, 20% methanol) (10 V, 1 h). The membrane was blocked with 5% milk powder in PBS-T (0.5% Tween-20 in PBS) for 1 h at RT and subsequently incubated with rabbit polyclonal anti-ClpP antibody (custom-made, raised against *S. aureus* ClpP, 2 mg/mL, 1:1000 dilution) in PBS-T + 5% milk powder (4 °C, overnight). The membrane was washed three times with PBS-T (15 min, RT) and incubated with Pierce Goat anti-Rabbit poly-HRP secondary antibody (1:10 000, Thermo Fisher Scientific) in PBS-T + 5% milk powder (1 h, RT). The membrane was washed three times with PBS-T (15 min, RT) and chemiluminescence was detected after 10 min incubation with freshly prepared Clarity Western ECL Substrate (Bio-Rad) with a LAS-4000 gel scanning station (Fujitsu Life Sciences, Tokyo, Japan).

For detection of myc-tagged ClpP1 and ClpP2, pellets of *L. monocytogenes clpP1(191)*::2×myc and *L. monocytogenes clpP2(199)*::2×myc corresponding to 1 ml OD_600_=20 were prepared as described in the MS-based co-immunoprecipitation section without crosslinking. The pellets were resuspended in 200 μL 0.4% SDS-PBS and lyzed by sonication (3x 20s, 75%, cooled on ice during breaks). Protein concentration was determined using a BCA assay (Roti-Quant universal, Carl Roth GmbH + Co. KG, Karlsruhe, Germany), all samples were adjusted to 4 mg/mL with 2× Laemmli buffer and 20 μL of the samples were separated by SDS-PAGE (12.5% polyacrylamide, 150 V, 2.5 h). Protein transfer and detection was performed as described above, with a 1:5000 dilution of an anti-c-Myc antibody (rabbit polyclonal, ab152146, 1 mg/mL, Abcam, Cambridge, United Kingdom) in PBS-T + 5% milk powder used as primary antibody and the membrane was stained with Ponceau S.

#### Fluorescent labelling

25 mL BHI medium was inoculated with *L. monocytogenes* EGD-e, Δ*clpP1*, Δ*clpP2* and Δ*clpP1/2* from a day culture to an initial OD_600_ of 0.05. The culture was grown to early stationary phase and an amount corresponding to 800 μL OD_600_ = 20 was harvested (4000 g, 4 °C, 10 min). The cells were washed with 1 mL PBS (4000 g, 4 °C, 5 min). The pellets were resuspended in 800 μL PBS and aliquots of 250 μL were prepared. 2.5 μL 5 mM vibralactone probe (or 5 mM D3 or DMSO as controls) from a DMSO stock was added to all strains (2 h, RT). The cells were centrifuged (4000 g, 5 min, 4 °C), the supernatant was discarded, and the pellets were washed with 1 mL PBS (4000 g, 5 min, 4 °C). The pellets were stored at −80 °C until further usage. The cells were resuspended in 250 μL PBS and transferred to 2 mL tubes containing 0.5 mL inlets filled with glass beads of 0.5 mm diameter. The cells were lysed using 2× program #2 in Precellys 24 tissue homogenizer (Bertin Instruments, Montigny-le-Bretonneux, France) coupled to liquid N_2_-cooled Cryolys (flow rate set to level I during shaking, level 0 during breaks). 200 μL of the lysates were pipetted into microcentrifuge tubes and the insoluble fractions were separated (10 000 g, 30 min, 4 °C). Click reagents [2 μL 5 mM rhodamine azide, 2 μL 15 mg/mL TCEP, 6 μL 1.67 mM tris((1-benzyl-4-triazolyl)methyl)amine ligand and 2 μL 50 mM CuSO4] were added to 88 μL of the supernatant and the reactions were incubated in the dark for 1 h at RT. 2×Laemmli buffer was added and the samples were stored at −20 °C until further usage. 50 μL of the samples were separated by SDS-PAGE (12.5% polyacrylamide, 150 V, 3 h) and fluorescence was detected with LAS-4000 gel scanning station (Fujitsu Life Sciences).

#### Growth curves of *L. monocytogenes* mutants

In the inner wells of a transparent flat-bottom 96-well plate, 200 μL BHI medium (if required, supplemented with 100 ppm H_2_O_2_) were inoculated to a starting OD_600_ of 0.01 with overnight cultures of *L. monocytogenes* EDG-e and its mutants (Δ*clpP1*, Δ*clpP2* and Δ*clpP1/2*) or left sterile for blank measurements. The outer wells of the plate were filled with 200 μL BHI medium but were not measured. The plate was covered with a transparent lid and was incubated at 37 °C with 5 s shaking every 15 min in an infinite M200Pro plate reader (Tecan). OD_600_ was measured every 15 min for 24 h. Data was recorded in triplicates and at least two independent experiments were conducted with qualitatively identical results. Plots were made with GraphPad Prism 6.

#### Intracellular growth assay

##### Cultivation of the J774A.1 cell line

J774A.1 murine macrophage-like cells were grown in tissue culture flasks with hydrophobic surface for suspension cells in DMEM/FCS (DMEM high glucose medium (Sigma-Aldrich, St. Louis, United States) supplemented with 2 mM glutamine and 10% heat-deactivated FCS). The flasks were incubated at 37 °C under 5% CO_2_. The cells were splitted into new flasks every 2–3 days to ca. 5×10^4^ cells/cm^2^. For detachment, cells were washed twice with TEN buffer (40 mM Tris-HCl, 150 mM NaCl, 1 mM EDTA, pH 7.4) and incubated with Accutase solution (Sigma-Aldrich) at 37 °C for 30 min.

*Intracellular growth assay* 10^5^ J774A.1 cells in 100 μL DMEM/FCS were pipetted into the inner wells of a flat-bottom 96-well plate. The outer wells were filled with 150 μL sterile PBS. The plates were incubated overnight at 37 °C under 5% CO_2_. On the next day, DMEM/FCS was inoculated with *L. monocytogenes* EDG-e, Δ*clpP1*, Δ*clpP2* and Δ*clpP1/2* overnight cultures to 10^3^ CFU/μL. The J774A.1 cells were washed with 150 μL PBS and 100 μL bacterial suspension was added (multiplicity of infection = 0.5). The plate was incubated on ice for 15 min and at 37 °C for 15 min. The cells were washed three times with 200 μL PBS. 150 μL DMEM/FCS supplemented with 10 μL gentamycin was added to kill extracellular bacteria. The plates were incubated at 37 °C under 5% CO_2_. After 7 h, the cells were washed three times with 200 μL PBS, and lysed with 2× 100 μL 0.05% Triton X-100 in ddH_2_O (1 min, RT). Dilution series were prepared from the lysates and plated on BHI agar plates. The agar plates were incubated at 37 °C for 2 days until colonies were counted. Data was recorded in triplicates and two independent experiments were performed. Plots were made with GraphPad Prism 6.

#### Whole-proteome analysis

*Cultivation of L. monocytogenes* 3×5 mL BHI medium (3 technical replicates) were inoculated 1:100 with overnight cultures of *L. monocytogenes* EGD-e, Δ*clpP1*, Δ*clpP2* and Δ*clpP1/2*. The first day culture was grown to an OD_600_ of ca. 0.5 at 37 °C under shaking at 200 rpm. For the second day culture, 3×5 mL BHI medium was inoculated with the first day culture to a starting OD_600_ of 0.05 and incubated at 37 °C or 42 °C under shaking at 200 rpm. After reaching early stationary phase, 1.5 mL of the cultures were harvested (4000 g, 10 min, 4 °C). The pellet was washed with 1 mL PBS and stored at −80 °C until further usage. Two biological replicates were generated.

##### Cell lysis and protein precipitation

Bacteria were resuspended in 150 μL lysis buffer (1% Triton X100, 0.5% SDS, 1 tablet cOmplete EDTA-free protease inhibitor cocktail (Roche Diagnostics GmbH, Mannheim, Germany) in 10 mL PBS) and lysed by ultrasonication (5×20 s, 80%, on ice during breaks). Cell debris was pelletized (5000 g, 10 min, 4 °C) and the supernatant was sterile filtered through a 0.2 μm pore size PTFE filter. Protein concentration was determined using a BCA assay (Roti-Quant universal, Carl Roth GmbH + Co. KG), all samples were adjusted to the same volume and concentration (ca. 1 mg/mL) and transferred to protein low-bind microcentrifuge tubes (Eppendorf, Hamburg, Germany). To precipitate the proteins, 4× sample volume acetone (−80 °C) was added and the samples were stored at −80 °C overnight. The samples were centrifuged at 21 000 g at 4 °C for 15 min and the supernatant was discarded. The pellet was resuspended in 500 μL methanol (−80 °C) with ultrasonication (10 s, 10%). After centrifugation at 21 000 g and at 4 °C for 15 min, the supernatant was discarded and the pellet was air-dried.

*Sample preparation for LC-MS/MS* 200 μL X buffer (7 M urea, 2 M thiourea, 20 mM HEPES, pH 7.5) was added and the pellet was resuspended by ultrasonication (10 s, 10%). The samples were reduced by the addition of 0.2 μL 1 M DTT (45 min, RT, 450 rpm), alkylated with 2 μL 0.55 M iodoacetamide (IAA) (30 min, RT, 450 rpm) and the reaction was quenched with 0.8 μL 1 M DTT (30 min, RT, 450 rpm). The samples were pre-digested with 0.5 μg/μL LysC (2 h, RT, 450 rpm). For the tryptic digest, 600 μL 50 mM triethylammonium bicarbonate (TEAB) buffer and 0.5 μg/μL trypsin (sequencing grade, modified, Promega) was added (overnight, 37 °C, 450 rpm). The pH was set to < 3 with 10 μL formic acid (FA). The samples were desalted on a Sep-Pak C18 50 mg column (Waters) using gravity flow. The columns were equilibrated with 1 mL MeCN, 0.5 mL 80% MeCN + 0.5% FA and 3× 1 mL 0.1% trifluoroacetic acid (TFA). The samples were loaded on the column and washed with 2× 1 mL 0.1% TFA and with 250 μL 0.5% FA. The peptides were eluted with 3× 250 μL MeCN, 0.5 mL 80% MeCN + 0.5% FA using vacuum in the last step. The solvents were removed under vacuum at 30 °C and the samples were resuspended in 1% FA (volume set to 2 μg/μL protein concentration), with pipetting up and down, 15 min ultrasonication in water bath and vortexing. The samples were filtered through a 0.2 μm pore size centrifugal filter.

##### LC-MS/MS

Samples were analyzed by LC-MS/MS using an UltiMate 3000 nano HPLC system (Thermo Fisher Scientific) equipped with an Acclaim C18 PepMap100 75 μm ID × 2 cm trap and an Aurora C18 separation column (75 μM ID x 25 cm, Ionopticks, Fitzroy, Australia) coupled to an Orbitrap Fusion (Thermo Fisher Scientific). Whole proteome and anti-ClpP XL-co-IP experiments performed at 37 °C were analyzed with the same setup, but with an Acclaim Pepmap RSLC C18 separation column (75 μm ID × 50 cm) in an EASY-spray setting. Injected samples were loaded on the trap column with a flow rate of 5 μL/min with 0.1% TFA buffer and then transferred onto the separation column at a flow rate of 0.4 μL/min (0.3 μL/min in case an Acclaim Pepmap RSLC C18 separation column was used). Samples were separated using a 152 min gradient (buffer A: H2O with 0.1% FA, buffer B: MeCN with 0.1% FA, gradient: 5% buffer B for 7 min, from 5% to 22% buffer B in 105 min, then to 32% buffer B in 10 min, to 90% buffer B in 10 min and hold at 90% buffer B for 10 min, then to 5% buffer B in 0.1 min and hold 5% buffer B for 9.9 min). Peptides were ionized using a nanospray source at 1.7–1.9 kV and a capillary temperature of 275 °C. The instrument was operated in a top speed data dependent mode with a cycle time between master scans of 3 s. MS full scans were performed in the orbitrap with quadrupole isolation at a resolution of R = 120 000 and an automatic gain control (AGC) ion target value of 2×10^5^ in a scan range of 300–1500 m/z with a maximum injection time of 50 ms. Internal calibration was performed using the ion signal of fluoranthene cations (EASY-ETD/IC source). Dynamic exclusion time was set to 60 s with a mass tolerance of 10 ppm (low/high). Precursors with intensities higher than 5×10^3^ and charge states 2–7 were selected for fragmentation with HCD (30%). MS2 scans were recorded in the ion trap operating in a rapid mode with an isolation window of 1.6 m/z. The AGC target was set to 1×10^4^ with a maximum injection time of 35 ms (100 ms in case of the temperature-dependent co-IP with anti-c-Myc antibody) and the “inject ions for all available parallelizable time” was enabled.

##### Data analysis

MS raw data were analyzed with MaxQuant 1.6.5.0 and default settings were used, except for the following: label-free quantification and match between runs were activated. All replicates for one condition (n = 6) were set as one fraction. The UniProt database of *L. monocytogenes* EGD-e proteins (taxon ID: 169963, downloaded on 25.01.2019) was searched. Data was further analyzed with Perseus 1.6.2.3. The rows “only identified by site”, “potential contaminants” and “reverse” were filtered and the data were log_2_-transformed. Replicates were grouped and filtered to at least 4 valid values per at least one group. Missing values were imputed for the total matrix from normal distribution. Two-sample Student’s *t*-tests were performed with default settings. Iron containing proteins were searched for with the UniProt Keyword “Iron”. SOS response proteins were identified from van der Veen et al (van der Veen et al., 2010). UniProt keyword and GOBP term analyses were performed with aGOtool (agotool.org) (Schölz et al., 2015). Proteins with a fold change of ≥ 2 (upregulated) or ≤ −2 (downregulated) and a −log_10_ t-test *p*-value ≥ 1.3 were set as foreground. “compare_samples” was selected as enrichment method with majority protein IDs from the wild type whole proteome used as background. A p-value cutoff of 0.05 was set and overrepresented terms as well as multiple testing per category was used with no GO term subset. Terms associated with only one proteins as well as redundant parent terms were filtered.

### MS-based co-immunoprecipitation

#### Temperature-dependent co-IP with anti-c-Myc antibody

*Cultivation of L. monocytogenes with c-Myc-tagged clpP* 30 mL BHI medium were inoculated 1:100 with overnight cultures of *L. monocytogenes clpP1(191)*::2×myc and *L. monocytogenes clpP2(199)*::2×myc. The first day culture was grown to an OD_600_ of ca. 0.5 at 37 °C under shaking at 200 rpm. For the second day culture, 4×100 mL BHI medium was inoculated with the first day cultures to a starting OD_600_ of 0.05. 2 flasks per condition were incubated at 20 °C and at 42 °C under shaking at 200 rpm. After reaching early stationary phase, an amount of bacteria corresponding to 4×1 mL OD_600_ = 20 per flask was harvested (4000 g, 5 min, 4 °C) and washed with 1 mL PBS. The pellets were resuspended in 1 mL PBS and 2 mM DSSO was added (20 μL from a 100 mM DMSO stock). DSSO was kindly provided by Dr. Vadim Korotkov and Dr. Pavel Kielkowski and synthesized as described previously (Fux et al., 2019). The bacteria were incubated with the crosslinker for 30 min at 20 °C or 42 °C under shaking at 200 rpm. The reaction was quenched by washing twice with 50 mM Tris-HCl (pH 8.0) and the pellets were stored at −80 °C until further usage.

##### Cell lysis and co-IP

Bacteria were resuspended in 800 μL co-IP lysis buffer (50 mM Tris-HCl, 150 mM NaCl, 5% glycerol, pH 7.4) and 120 μg lysozyme was added. The samples were incubated at 37 °C under shaking at 1400 rpm for 1 h. Afterwards, 8 μL 10% NP-40 solution was added and the bacteria were lysed by ultrasonication (5×30 s, 80%, on ice during breaks). The insoluble fraction was pelletized (10 000 g, 30 min, 4 °C) and the supernatant was sterile filtered through a 0.2 μm PTFE filter. Protein concentration was determined using a BCA assay (Roti-Quant universal, Carl Roth GmbH + Co. KG). 30 μL Protein A/G agarose beads (Thermo Fisher Scientific) were transferred to protein low-bind microcentrifuge tubes (Eppendorf) and washed with 1 mL co-IP wash buffer (50 mM Tris-HCl, 150 mM NaCl, 5% glycerol, 0.05% NP-40, pH 7.4) and centrifuged for 1 min at 1000 g at 4 °C. 500 μg proteome (in 500 μL) and either 1 μL anti-c-Myc antibody (rabbit polyclonal, ab152146, 1 mg/mL, Abcam) or 0.4 μL isotype control (2.5 mg/mL, Cell Signaling Technology, Danvers, United States) were added. The samples were incubated at 4 °C for 3 h under constant rotation. The supernatant was removed after centrifugation (1000 g, 1 min, 4 °C), and the beads were washed twice with 1 mL co-IP wash buffer. The detergent was removed by washing the beads twice with co-IP lysis buffer.

##### Sample preparation for LC-MS/MS

The samples were reduced and digested in 25 μL co-IP digest buffer (50 mM Tris-HCl, 5 ng/μL trypsin (sequencing grade, modified, Promega), 2 M urea, 1 mM DTT, pH 8.0) at 25 °C under shaking at 600 rpm for 30 min. For alkylation, 100 μL 50 mM Tris-HCl, 2 mM urea, 5 mM IAA (pH 8.0) was added (25 °C, 600 rpm, 30 min). The digestion was completed overnight at 37 °C under shaking at 600 rpm. The pH was set to < 3 with 0.75 μL FA. The samples were desalted on double layer C18-stage tips (Empore disk-C18, Agilent Technologies, Santa Clara, United States). The stage tips were equilibrated with 70 μL methanol and 3× 70 μL 0.5% FA. The samples were loaded and washed with 3× 70 μL 0.5% FA. The peptides were eluted with 3× 30 μL 80% MeCN + 0.5% FA. The solvents were removed under vacuum at 30 °C and the samples were resuspended in 27 μL 1% FA with pipetting up and down, 15 min ultrasonication in water bath and vortexing. The samples were filtered through a 0.2 μm pore size centrifugal filter. LC-MS/MS measurement was conducted as described for the whole proteome analysis.

##### Data analysis

MS raw data were analyzed with MaxQuant 1.6.10.43. and default settings were used, except for the following: label-free quantification and match between runs were activated, *N*-acetylation modification was deactivated. All replicates for one condition (n = 4) were set as one fraction. The UniProt database of *L. monocytogenes* EGD-e proteins (taxon ID: 169963, downloaded on 21.10.2019.) was searched. Data was further analyzed with Perseus 1.6.10.43. The rows “only identified by site”, “potential contaminants” and “reverse” were filtered and the data were log2-transformed. Replicates were grouped and filtered to at least 3 valid values per at least one group. Missing values were imputed for the total matrix from normal distribution. Two-sample Student’s *t*-tests were performed with default settings.

### Co-IP with anti-clpP antibody

20 mL BHI medium was inoculated 1:100 with overnight cultures of *L. monocytogenes* Δ*clpP1* and Δ*clpP2*. The first day culture was grown to an OD_600_ of ca. 0.5 at 37 °C under shaking at 200 rpm. For the second day culture, 50 mL BHI medium was inoculated with the first day cultures to a starting OD_600_ of 0.05 and incubated at 37 °C or 42 °C under shaking at 200 rpm. After reaching early stationary phase, an amount of bacteria corresponding to 2×1 mL OD_600_ = 20 per replicate was harvested (4000 g, 5 min, 4 °C) and washed with 1 mL PBS. The pellets were resuspended in 1 mL PBS and 2 mM DSSO was added (20 μL from a 100 mM DMSO stock). The bacteria were incubated with the crosslinker for 30 min at 37 °C or 42 °C and under shaking at 200 rpm. The reaction was quenched by washing twice with 50 mM Tris-HCl (pH 8.0) and the pellets were stored at −80 °C until further usage. Four replicates from independent overnight cultures were generated for each experiment.

Cell lysis, co-IP and sample preparation were conducted as described for the temperature-dependent co-IP with anti-c-Myc antibody, except that either 5 μL anti-ClpP antibody (custom-made, polyclonal, raised against *S. aureus* ClpP in rabbit, 2 mg/mL) or 4 μL isotype control (2.5 mg/mL, Cell Signaling Technology) were used. 300 μg proteome was used in case of the 42 °C XL-co-IP. LC-MS/MS measurements and data analysis was done as described for the temperature-dependent co-IP with anti-c-Myc antibody. Oxidoreductases were searched for with the UniProt Keyword “Oxidoreductase”.

