## Supporting Information for "*Listeria monocytogenes* utilizes the ClpP1/2 proteolytic machinery for fine-tuned substrate degradation under heat stress"

**Important Note**

The mass spectrometry proteomics data will be deposited to the ProteomeXchange Consortium via the PRIDE (Perez-Riverol et al., 2019) partner repository upon final publication of the manuscript.

1. **Supplementary Figures**
2. **Supplementary Tables**
3. **Supplementary References**

### 1. Supplementary Figures

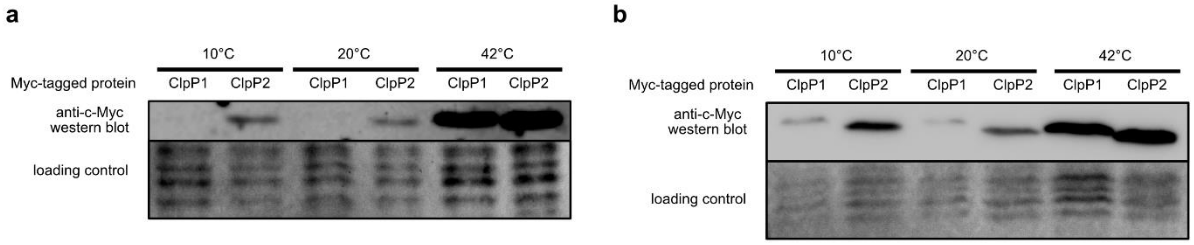

**Figure S1 Increased expression of ClpP1 and ClpP2 in *L. monocytogenes* at elevated temperatures. a** Western Blot of *L. monocytogenes clpP1::2×myc* and *clpP2::2×myc* cell lysates grown to stationary phase at 10 °C, 20 °C and 42 °C with an anti-c-Myc antibody. The membrane was stained using Ponceau S as loading control. The experiment was independently repeated with qualitatively similar results (**b**).

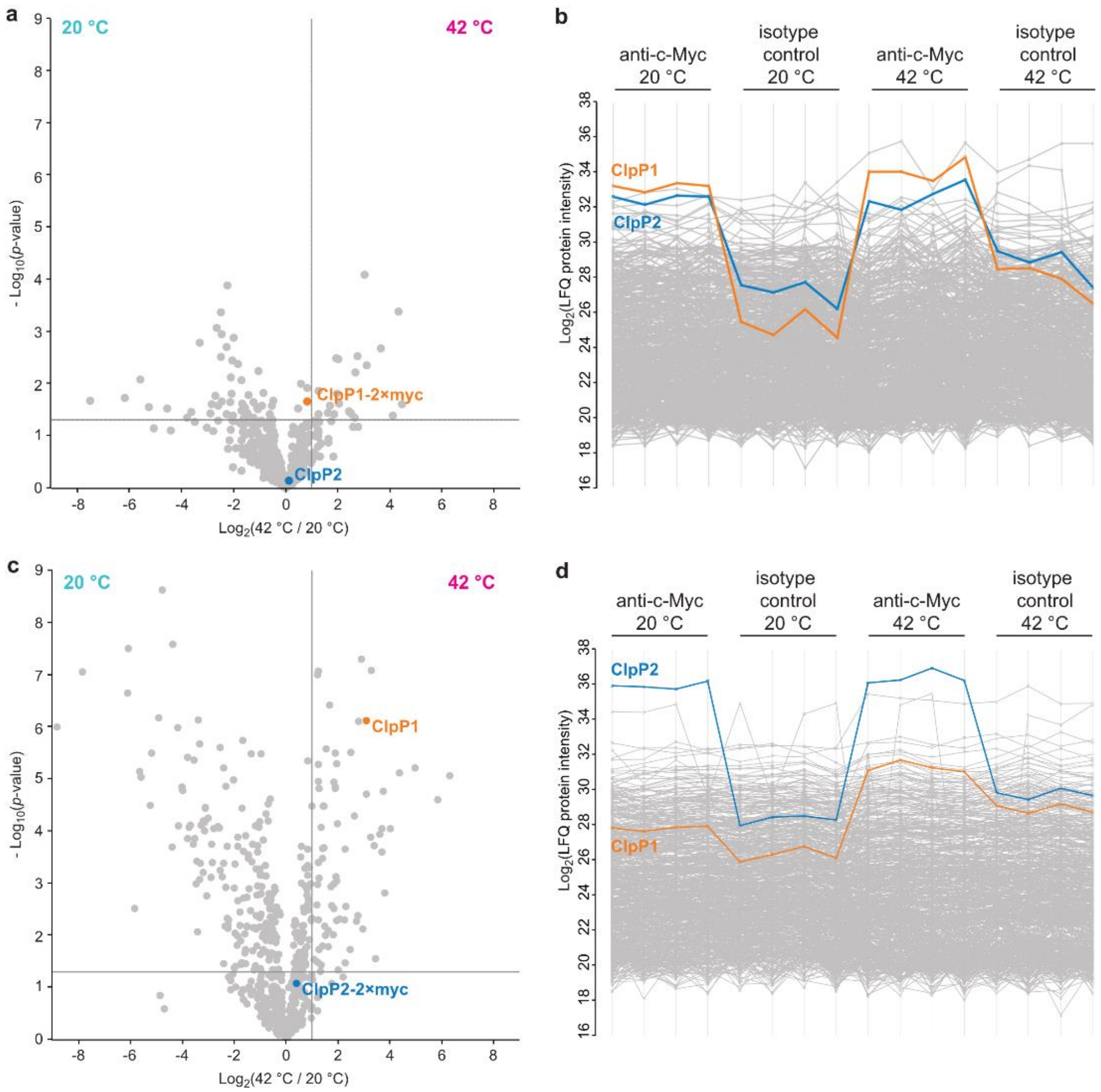

**Figure S2 Intracellular heterooligomerization of ClpP1 and ClpP2 in *L. monocytogenes* demonstrated by crosslinking co-immunoprecipitation. a, b** Volcano plots of co-IPs using ClpP1-2×myc (**a**) and ClpP2-2×myc (**c**) as baits after growing the *L. monocytogenes* cultures at 20 °C and

42 °C to stationary phase. − $\log_{10}$ *p*-values from two-samples Student's *t*-test are plotted against $\log_2$ ratios of label-free quantification (LFQ) protein intensities. The vertical grey lines show 2-fold enrichment at 42 °C compared to 20 °C, the horizontal grey lines show $-\log_{10}$ t-test *p*-value = 1.3 (n = 4). **b, d** Profile plots showing the $\log_2$ LFQ intensities of all measured proteins across all replicates after missing value imputation of the ClpP1-2×myc (**b**) and ClpP2-2×myc (**d**) XL-co-IP. ClpP1 and ClpP2 are highlighted with orange and blue, respectively. The lower number of enriched proteins obtained with the Myc-tagged ClpPs compared to the endogenous ClpPs using a polyclonal anti-ClpP antibody (Figure 7) could be attributed to the heterologous C-terminal 2×myc tag which is in close proximity to the hydrophobic pockets of ClpP and thus interfering with chaperone binding.

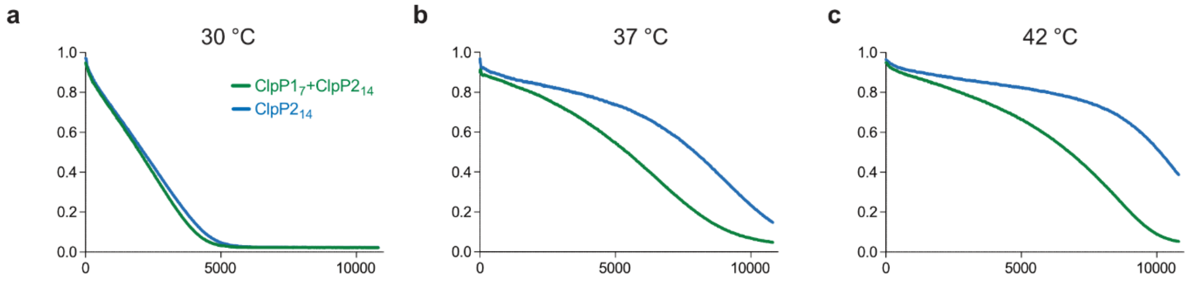

**Figure S3 Replicate measurements of ClpP1$_7$ and ClpP2$_{14}$ protease activity at different temperatures.** ClpP (green line: 0.1 µM ClpP2$_{14}$ and 0.2 µM ClpP1$_7$, blue line: 0.1 µM ClpP2$_{14}$) and 0.4 µM ClpX were pre-incubated for 30 min at 30 °C (**a**), 37 °C (**b**) and 42 °C (**c**), subsequently the degradation of 0.4 µM GFP-SsrA was measured. Means of triplicates are shown.

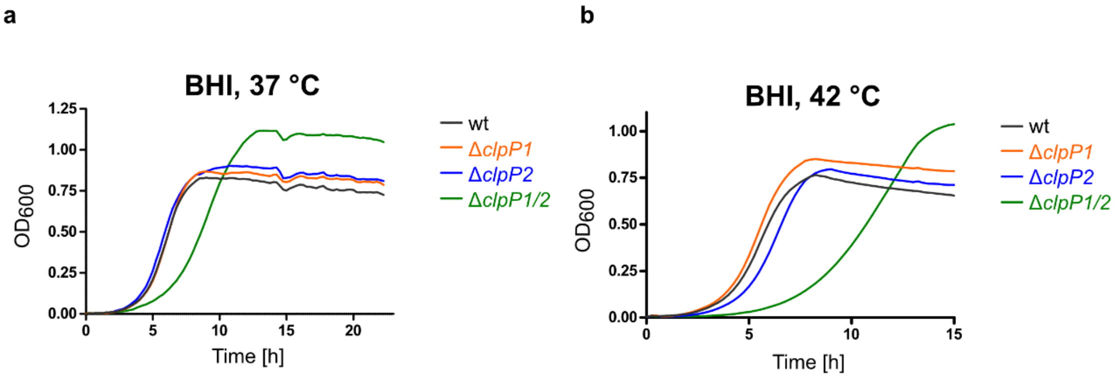

**Figure S4 Growth curves of *Listeria monocytogenes* EGD-e (WT) and Δ*clpP* mutants. a** Replicate growth curves of the ΔclpP mutants in BHI medium at 37 °C **b** Growth curves of the ΔclpP mutants in BHI medium at 42 °C. Means of triplicates are shown.

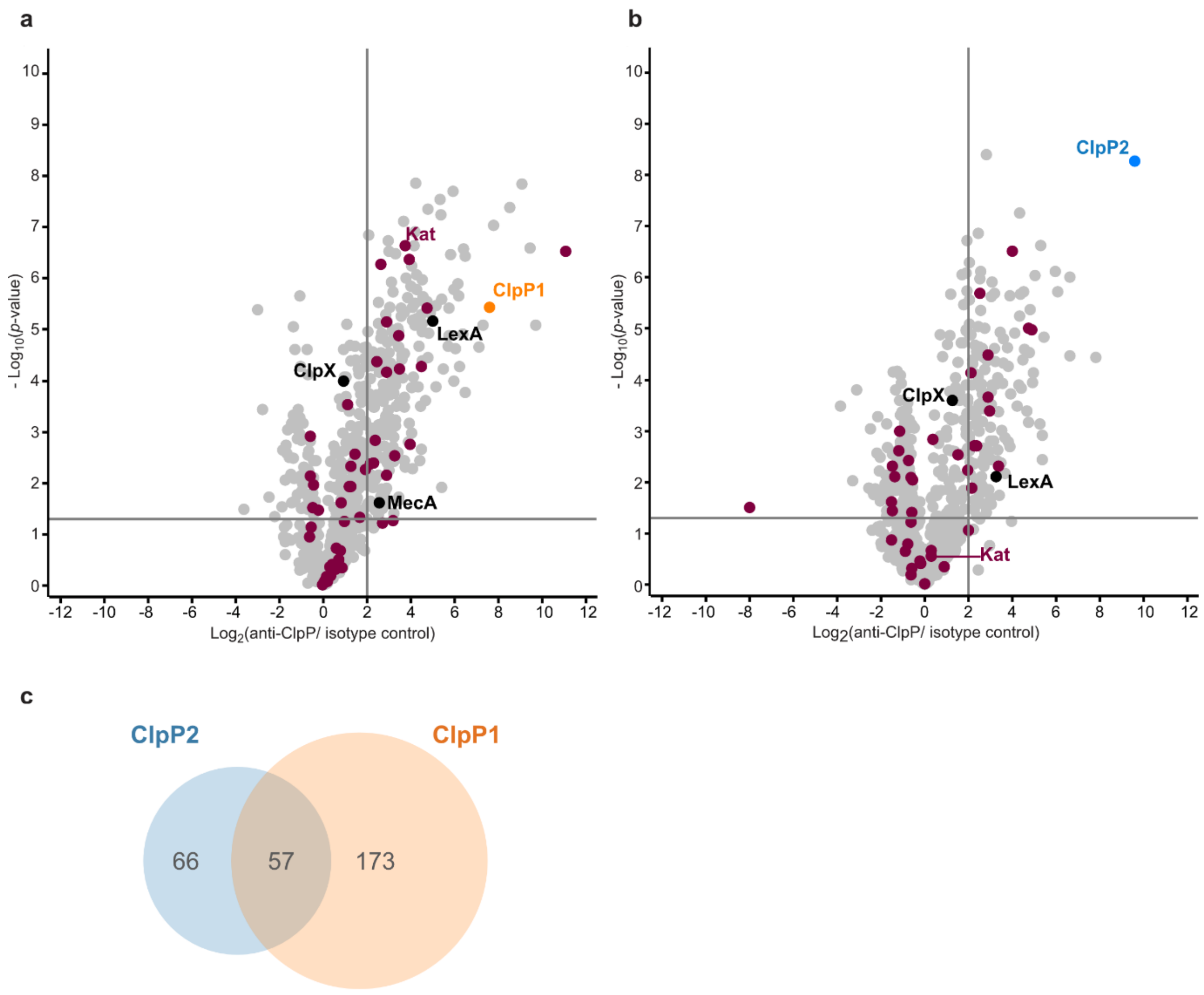

**Figure S5 Co-immunoprecipitation of ClpP1 and ClpP2 in *L. monocytogenes* Δ*clpP* mutants at 42 °C. a, b** Volcano plots of crosslinking co-IPs with anti-ClpP antibody in *L. monocytogenes* Δ*clpP2* (**a**) and Δ*clpP1* (**b**) at stationary phase (42 °C). − $\log_{10}$ *p*-values from two-sample Student's *t*-test are plotted against $\log_2$ ratios of LFQ protein intensities. The vertical grey lines show 4-fold enrichment, the horizontal grey lines show $-\log_{10}$ t-test *p*-value = 1.3 (n = 4). Oxidoreductases are highlighted with purple. ClpP1 and ClpP2 are shown in orange and blue respectively. **c** Venn diagram of putative interaction partners of ClpP1 and ClpP2 at 42 °C.

#### 2. Supplementary Tables

**Table S1** Dysregulated proteins in Δ*clpP1* whole proteome at 37 °C in reference to wildtype control.

| Gene name | Uniprot ID | $\log_2$ ratio (Δ*clpP1*/WT) | $-\log_{10}$ t-test p-value (Δ*clpP1*/WT) |
|---|---|---|---|
| **Upregulated proteins (37 °C)** | | | |
| lmo2768 | Q8Y3R5 | 2.10 | 1.75 |
| lmo2326 | Q8Y4V4 | 1.01 | 1.45 |

| | | | |
|---|---|---|---|
| **Downregulated proteins (37 °C)** | | | |
| lmo0488 | Q8Y9N7 | -1.00 | 1.37 |
| lmo2044 | Q8Y5L4 | -1.19 | 2.00 |
| lmo2516 | Q8Y4D4 | -1.35 | 1.72 |
| lmo1189 | Q8Y7T2 | -1.38 | 1.88 |
| ispD | Q8YAB5 | -1.41 | 1.82 |
| clpP | Q8Y7Y1 | -8.63 | 6.80 |

**Table S2** Dysregulated proteins in Δ*clpP1* whole proteome at 42 °C in reference to wildtype control.

| Gene name | Uniprot ID | $log_2$ ratio (Δ*clpP1*/WT) | $-log_{10}$ t-test p-value (Δ*clpP1*/WT) |
|---|---|---|---|
| **Upregulated proteins (42 °C)** | | | |
| lmo0375 | Q8Y9Z2 | 2.93 | 2.45 |
| fhuC | Q8Y5U5 | 2.25 | 2.18 |
| lmo1307 | Q8Y7H3 | 2.07 | 1.42 |
| inlB | P25147 | 1.40 | 4.10 |
| lmo2755 | Q8Y3S6 | 1.24 | 1.66 |
| hly | P13128 | 1.24 | 4.14 |
| spl | Q7AP49 | 1.08 | 3.64 |
| **Downregulated proteins (42 °C)** | | | |
| lmo2573 | Q8Y482 | -1.08 | 1.99 |
| lmo0953 | Q8Y8F1 | -1.09 | 1.66 |
| lmo2795 | Q8Y3P0 | -1.12 | 1.57 |
| lmo2088 | Q8Y5H4 | -1.17 | 2.26 |
| lmo0273 | Q8YA86 | -1.20 | 1.50 |
| nifJ | Q8Y8R6 | -1.24 | 1.94 |
| ung2 | Q8Y7P6 | -1.32 | 1.65 |
| cspD | Q92AD0 | -1.43 | 1.96 |
| lmo0913 | Q8Y8I9 | -1.52 | 3.79 |
| lmo0887 | Q8Y8L1 | -1.53 | 2.08 |
| lmo1642 | Q8Y6P2 | -1.63 | 2.04 |
| lmo0612 | Q8Y9C0 | -1.66 | 1.43 |
| lmo1868 | Q8Y632 | -1.79 | 3.25 |
| msrA | Q8Y640 | -1.99 | 1.91 |
| lmo0927 | Q8Y8H6 | -2.07 | 3.33 |

| | | | |
|---|---|---|---|
| lmo0253 | Q8YAA1 | -2.32 | 2.62 |
| lmo2258 | Q8Y520 | -2.62 | 1.43 |
| addB | Q8Y510 | -2.72 | 1.91 |
| lmo1505 | Q8Y721 | -2.77 | 2.92 |
| lmo2685 | Q927F6 | -3.17 | 6.12 |
| lmo2805 | Q8Y3N0 | -3.82 | 3.01 |
| atpE | Q8Y4B7 | -5.41 | 7.99 |
| clpP | Q8Y7Y1 | -10.65 | 6.87 |

**Table S3** Significantly enriched GOBP terms of upregulated proteins in Δ*clpP2* and Δ*clpP12* whole proteomes compared to WT at 37 °C identified with aGOtool (Schölz et al., 2015). Terms unique for either whole proteome are highlighted in bold.

| **Δ*clpP2*/WT, 37 °C** | | | | **ΔclpP12/WT, 37 °C** | | | |
|---|---|---|---|---|---|---|---|
| **Rank** | **s value** | **-log10 (p-value)** | **Description (GOBP)** | **Rank** | **s value** | **-log10 (p-value)** | **Description (GOBP)** |
| 1 | 0.074 | 1.90 | regulation of transcription, DNA-templated | 1 | 0.259 | 3.65 | response to stimulus |
| 2 | 0.069 | 1.82 | response to stimulus | 2 | 0.183 | 3.20 | cellular response to stimulus |
| 3 | 0.049 | 1.61 | cellular response to stimulus | 3 | 0.163 | 3.07 | response to stress |
| 4 | 0.036 | 1.62 | response to chemical | 4 | 0.138 | 2.54 | regulation of transcription, DNA-templated |
| 5 | 0.032 | 1.32 | response to stress | 5 | 0.102 | 2.45 | **cellular response to stress** |
| 6 | 0.027 | 1.63 | response to inorganic substance | 6 | 0.101 | 2.48 | **cellular response to DNA damage stimulus** |
| 7 | 0.027 | 1.63 | cellular response to chemical stimulus | 7 | 0.088 | 2.32 | **DNA repair** |
| 8 | 0.020 | 1.43 | response to metal ion | 8 | 0.087 | 2.15 | **DNA metabolic process** |
| 9 | 0.018 | 1.44 | **cellular response to iron ion** | 9 | 0.054 | 1.41 | **regulation of cellular process** |
| 10 | 0.018 | 1.33 | glucose metabolic process | 10 | 0.053 | 1.88 | response to chemical |
| | | | | 11 | 0.031 | 1.69 | **alpha-amino acid catabolic process** |
| | | | | 12 | 0.031 | 1.69 | glucose metabolic process |
| | | | | 13 | 0.029 | 1.59 | response to inorganic substance |
| | | | | 14 | 0.029 | 1.59 | cellular response to chemical stimulus |
| | | | | 15 | 0.027 | 1.43 | **DNA recombination** |
| | | | | 16 | 0.027 | 1.43 | **monosaccharide metabolic process** |
| | | | | 17 | 0.019 | 1.33 | response to metal ion |
| | | | | 18 | 0.017 | 1.50 | **threonine catabolic process** |

| | | | | | | | |
|---|---|---|---|---|---|---|---|
| | | | | 19 | 0.014 | 1.33 | **tetrahydrofolate interconversion** |

**Table S4** Significantly enriched UniProt keywords of upregulated proteins in Δ*clpP2* and Δ*clpP12* whole proteomes compared to WT at 37 °C identified with aGOtool (Schölz et al., 2015). Terms unique for either whole proteome are highlighted in bold.

| **ΔclpP2/WT, 37 °C** | | | | **ΔclpP12/WT, 37 °C** | | | |
|---|---|---|---|---|---|---|---|
| **Rank** | **s value** | **$-log_{10}$ (p-value)** | **Description (Keywords)** | **Rank** | **s value** | **$-log_{10}$ (p-value)** | **Description (Keywords)** |
| 1 | 0.183 | 4.44 | Iron-sulfur | 1 | 0.245 | 4.47 | Iron |
| 2 | 0.168 | 3.83 | Iron | 2 | 0.242 | 4.89 | Iron-sulfur |
| 3 | 0.132 | 3.75 | 4Fe-4S | 3 | 0.214 | 4.63 | 4Fe-4S |
| 4 | 0.036 | 1.44 | Transcription regulation | 4 | 0.050 | 1.62 | Transcription regulation |
| 5 | 0.014 | 1.48 | Organic radical | 5 | 0.046 | 1.79 | **DNA repair** |
| | | | | 6 | 0.044 | 1.42 | **Oxidoreductase** |
| | | | | 7 | 0.020 | 1.71 | Organic radical |
| | | | | 8 | 0.020 | 1.48 | **Heme** |

**Table S5** Significantly enriched GOBP terms of downregulated proteins in Δ*clpP2* and Δ*clpP12* whole proteomes compared to WT at 37 °C identified with aGOtool (Schölz et al., 2015). Terms unique for either whole proteome are highlighted in bold.

| **ΔclpP2/WT, 37 °C** | | | | **ΔclpP12/WT, 37 °C** | | | |
|---|---|---|---|---|---|---|---|
| **Rank** | **s value** | **$-log_{10}$ (p-value)** | **Description (GOBP)** | **Rank** | **s value** | **$-log_{10}$ (p-value)** | **Description (GOBP)** |
| 1 | 0.275 | 2.84 | **organonitrogen compound biosynthetic process** | 1 | 0.246 | 3.70 | 'de novo' UMP biosynthetic process |
| 2 | 0.162 | 3.23 | **nitrogen compound transport** | 2 | 0.078 | 2.30 | secondary alcohol metabolic process |
| 3 | 0.129 | 1.72 | **cellular biosynthetic process** | 3 | 0.049 | 1.56 | arginine biosynthetic process |
| 4 | 0.119 | 2.07 | **transport** | 4 | 0.049 | 1.56 | amino acid transport |
| 5 | 0.115 | 1.60 | **biosynthetic process** | 5 | 0.049 | 1.56 | pyrimidine nucleobase biosynthetic process |
| 6 | 0.114 | 2.26 | **organic substance transport** | 6 | 0.039 | 1.32 | **proteolysis** |
| 7 | 0.110 | 1.56 | **organic substance biosynthetic process** | | | | |
| 8 | 0.105 | 1.55 | **organonitrogen compound metabolic process** | | | | |

| | | | |
|---|---|---|---|
| 9 | 0.070 | 2.47 | **branched-chain amino acid biosynthetic process** |
| 10 | 0.070 | 2.72 | 'de novo' UMP biosynthetic process |
| 11 | 0.067 | 1.43 | **cellular macromolecule biosynthetic process** |
| 12 | 0.064 | 1.57 | **peptide metabolic process** |
| 13 | 0.063 | 1.59 | **translation** |
| 14 | 0.058 | 2.38 | **isoleucine biosynthetic process** |
| 15 | 0.054 | 1.81 | **ribonucleoside monophosphate biosynthetic process** |
| 16 | 0.046 | 1.84 | **peptide transport** |
| 17 | 0.045 | 1.98 | **pathogenesis** |
| 18 | 0.045 | 1.70 | **cell wall macromolecule biosynthetic process** |
| 19 | 0.044 | 2.17 | **valine biosynthetic process** |
| 20 | 0.043 | 1.64 | **peptidoglycan-based cell wall biogenesis** |
| 21 | 0.042 | 1.31 | **cellular amino acid biosynthetic process** |
| 22 | 0.042 | 1.33 | **alpha-amino acid biosynthetic process** |
| 23 | 0.040 | 2.01 | amino acid transport |
| 24 | 0.038 | 1.60 | **peptidoglycan metabolic process** |
| 25 | 0.030 | 1.50 | **peptidoglycan biosynthetic process** |
| 26 | 0.027 | 1.51 | **protein transport** |
| 27 | 0.024 | 1.41 | **cellular localization** |
| 28 | 0.019 | 1.35 | arginine biosynthetic process |
| 29 | 0.019 | 1.35 | **threonine biosynthetic process** |
| 30 | 0.019 | 1.35 | pyrimidine nucleobase biosynthetic process |
| 31 | 0.015 | 1.43 | **methionine transport** |
| 32 | 0.015 | 1.43 | **pyridoxal phosphate biosynthetic process** |
| 33 | 0.015 | 1.43 | secondary alcohol metabolic process |

**Table S6** Significantly enriched UniProt keywords of downregulated proteins in in Δ*clpP2* and Δ*clpP12* whole proteomes compared to WT at 37 °C identified with aGOtool (Schölz et al., 2015). Terms unique for either whole proteome are highlighted in bold.

| **ΔclpP2/WT, 37 °C** | | | | **ΔclpP12/WT, 37 °C** | | | |
|---|---|---|---|---|---|---|---|
| **Rank** | **s value** | **$-\log_{10}$ (p-value)** | **Description (Keywords)** | **Rank** | **s value** | **$-\log_{10}$ (p-value)** | **Description (Keywords)** |
| 1 | 0.814 | 5.45 | Membrane | 1 | 1.213 | 5.02 | Transmembrane helix |
| 2 | 0.778 | 5.42 | **Transmembrane** | 2 | 1.143 | 4.74 | Membrane |
| 3 | 0.708 | 5.12 | Transmembrane helix | 3 | 0.323 | 2.74 | Transport |
| 4 | 0.685 | 6.06 | Transport | 4 | 0.221 | 3.38 | Pyrimidine biosynthesis |
| 5 | 0.533 | 5.08 | **Cell membrane** | 5 | 0.133 | 1.74 | Signal |
| 6 | 0.498 | 5.25 | Signal | 6 | 0.058 | 1.78 | **Peptidoglycan-anchor** |
| 7 | 0.174 | 3.17 | **Ribosomal protein** | 7 | 0.053 | 1.66 | Serine protease |
| 8 | 0.094 | 2.83 | **Lipoprotein** | 8 | 0.053 | 1.66 | **Cell wall** |
| 9 | 0.075 | 2.60 | **Palmitate** | 9 | 0.049 | 1.56 | **Decarboxylase** |
| 10 | 0.058 | 2.38 | Pyrimidine biosynthesis | | | | |
| 11 | 0.054 | 2.23 | **Branched-chain amino acid biosynthesis** | | | | |
| 12 | 0.045 | 1.98 | **Virulence** | | | | |
| 13 | 0.036 | 1.86 | **Protein transport** | | | | |
| 14 | 0.028 | 1.42 | **Cell wall biogenesis/degradation** | | | | |
| 15 | 0.028 | 1.80 | **Threonine biosynthesis** | | | | |
| 16 | 0.024 | 1.63 | **Amino-acid transport** | | | | |
| 17 | 0.022 | 1.32 | **Peptidoglycan synthesis** | | | | |
| 18 | 0.021 | 1.48 | Serine protease | | | | |

**Table S7** Significantly enriched GOBP terms of upregulated proteins in Δ*clpP2* and Δ*clpP12* whole proteomes compared to WT at 42 °C identified with aGOtool (Schölz et al., 2015). Terms unique for either whole proteome are highlighted in bold.

| **ΔclpP2/WT, 42 °C** | | | | **ΔclpP12/WT, 42 °C** | | | |
|---|---|---|---|---|---|---|---|
| **Rank** | **s value** | **$-\log_{10}$ (p-value)** | **Description (GOBP)** | **Rank** | **s value** | **$-\log_{10}$ (p-value)** | **Description (GOBP)** |
| 1 | 0.092 | 2.28 | regulation of transcription, DNA-templated | 1 | 0.159 | 3.11 | regulation of transcription, DNA-templated |
| 2 | 0.089 | 2.22 | regulation of nucleobase-containing compound metabolic process | 2 | 0.155 | 3.03 | regulation of nucleobase-containing compound metabolic process |
| 3 | 0.078 | 2.10 | response to stimulus | 3 | 0.152 | 2.94 | **regulation of nitrogen compound metabolic process** |
| 4 | 0.076 | 2.25 | response to stress | 4 | 0.151 | 2.96 | **regulation of cellular macromolecule biosynthetic process** |

| | | | | | | | |
|---|---|---|---|---|---|---|---|
| 5 | 0.056 | 2.03 | cellular response to DNA damage stimulus | 5 | 0.147 | 2.89 | **regulation of primary metabolic process** |
| 6 | 0.054 | 1.83 | cellular response to stimulus | 6 | 0.142 | 2.84 | **regulation of gene expression** |
| 7 | 0.054 | 1.63 | regulation of macromolecule metabolic process | 7 | 0.138 | 2.77 | **regulation of cellular metabolic process** |
| 8 | 0.054 | 1.86 | DNA metabolic process | 8 | 0.138 | 3.15 | response to stress |
| 9 | 0.050 | 1.86 | cellular response to stress | 9 | 0.134 | 2.72 | regulation of macromolecule metabolic process |
| 10 | 0.042 | 1.75 | DNA repair | 10 | 0.104 | 2.45 | response to stimulus |
| 11 | 0.020 | 1.59 | arginine biosynthetic process | 11 | 0.093 | 2.67 | cellular response to DNA damage stimulus |
| 12 | 0.014 | 1.34 | **base-excision repair** | 12 | 0.088 | 2.45 | DNA metabolic process |
| 13 | 0.012 | 1.45 | **base-excision repair, AP site formation** | 13 | 0.084 | 2.47 | cellular response to stress |
| | | | | 14 | 0.074 | 2.17 | cellular response to stimulus |
| | | | | 15 | 0.073 | 2.36 | DNA repair |
| | | | | 16 | 0.053 | 1.52 | **regulation of biological process** |
| | | | | 17 | 0.029 | 1.76 | **response to external stimulus** |
| | | | | 18 | 0.025 | 1.64 | **cellular response to extracellular stimulus** |
| | | | | 19 | 0.023 | 1.63 | **SOS response** |
| | | | | 20 | 0.021 | 1.64 | arginine biosynthetic process |
| | | | | 21 | 0.020 | 1.37 | **DNA recombination** |
| | | | | 22 | 0.014 | 1.40 | **nucleotide-excision repair** |

**Table S8** Significantly enriched UniProt keywords of upregulated proteins in Δ*clpP2* and Δ*clpP12* whole proteomes compared to WT at 42 °C identified with aGOtool (Schölz et al., 2015). Terms unique for either whole proteome are highlighted in bold.

| **ΔclpP2/WT, 42 °C** | | | | **ΔclpP12/WT, 42 °C** | | | |
|---|---|---|---|---|---|---|---|
| **Rank** | **s value** | **$-\log_{10}$ (p-value)** | **Description (Keywords)** | **Rank** | **s value** | **$-\log_{10}$ (p-value)** | **Description (Keywords)** |
| 1 | 0.076 | 2.87 | Iron-sulfur | 1 | 0.049 | 2.23 | Iron-sulfur |
| 2 | 0.058 | 2.52 | 4Fe-4S | 2 | 0.047 | 2.23 | 4Fe-4S |
| 3 | 0.045 | 1.94 | Iron | 3 | 0.038 | 1.41 | **DNA-binding** |
| 4 | 0.040 | 1.89 | DNA repair | 4 | 0.038 | 1.75 | Iron |
| 5 | 0.031 | 1.42 | **Transcription regulation** | 5 | 0.033 | 1.68 | DNA repair |
| 6 | 0.028 | 1.31 | **Transcription** | 6 | 0.024 | 1.79 | Arginine biosynthesis |
| 7 | 0.023 | 1.74 | Arginine biosynthesis | 7 | 0.019 | 1.77 | **Excision nuclease** |
| | | | | 8 | 0.019 | 1.77 | **DNA excision** |

**Table S9** Significantly enriched GOBP terms of downregulated in Δ*clpP2* and Δ*clpP12* whole proteomes compared to WT at 42 °C identified with aGOtool (Schölz et al., 2015). Terms unique for either whole proteome are highlighted in bold.

| **ΔclpP2/WT, 42 °C** | | | | **ΔclpP12/WT, 42 °C** | | | |
|---|---|---|---|---|---|---|---|
| **Rank** | **s value** | **$-\log_{10}$ (p-value)** | **Description (GOBP)** | **Rank** | **s value** | **$-\log_{10}$ (p-value)** | **Description (GOBP)** |
| 1 | 0.693 | 5.87 | **cellular macromolecule biosynthetic process** | 1 | 0.163 | 4.11 | 'de novo' UMP biosynthetic process |
| 2 | 0.574 | 6.00 | **translation** | 2 | 0.115 | 2.01 | transport |
| 3 | 0.562 | 5.83 | **peptide metabolic process** | 3 | 0.094 | 2.05 | **cellular catabolic process** |
| 4 | 0.443 | 4.77 | **cellular protein metabolic process** | 4 | 0.084 | 2.32 | **organic hydroxy compound metabolic process** |
| 5 | 0.433 | 4.86 | **amide biosynthetic process** | 5 | 0.076 | 1.75 | organic substance transport |
| 6 | 0.328 | 3.38 | **cellular macromolecule metabolic process** | 6 | 0.072 | 2.08 | pyrimidine-containing compound metabolic process |
| 7 | 0.321 | 3.92 | **cellular amide metabolic process** | 7 | 0.058 | 1.81 | **ribonucleoside monophosphate biosynthetic process** |
| 8 | 0.320 | 3.13 | **organonitrogen compound metabolic process** | 8 | 0.050 | 2.03 | **cellular carbohydrate catabolic process** |
| 9 | 0.314 | 3.27 | **organonitrogen compound biosynthetic process** | 9 | 0.050 | 2.03 | pathogenesis |
| 10 | 0.301 | 3.58 | **gene expression** | 10 | 0.049 | 2.22 | **glycerol catabolic process** |
| 11 | 0.258 | 3.01 | **cellular nitrogen compound biosynthetic process** | 11 | 0.045 | 1.46 | **small molecule catabolic process** |
| 12 | 0.196 | 2.42 | **macromolecule metabolic process** | 12 | 0.044 | 2.05 | pyrimidine nucleobase biosynthetic process |
| 13 | 0.166 | 1.96 | **cellular process** | 13 | 0.042 | 1.48 | **carbohydrate transport** |
| 14 | 0.163 | 2.09 | **cellular biosynthetic process** | 14 | 0.037 | 1.63 | nucleobase metabolic process |
| 15 | 0.162 | 2.07 | **organic substance biosynthetic process** | 15 | 0.032 | 1.39 | **cellular carbohydrate metabolic process** |
| 16 | 0.092 | 3.31 | 'de novo' UMP biosynthetic process | 16 | 0.032 | 1.46 | **alcohol metabolic process** |
| 17 | 0.090 | 1.40 | **cellular metabolic process** | 17 | 0.031 | 1.83 | 'de novo' pyrimidine nucleobase biosynthetic process |
| 18 | 0.089 | 3.00 | **pyrimidine ribonucleotide biosynthetic process** | 18 | 0.024 | 1.36 | macromolecule localization |
| 19 | 0.077 | 1.37 | **cellular nitrogen compound metabolic process** | 19 | 0.017 | 1.45 | **pyridoxal phosphate biosynthetic process** |
| 20 | 0.076 | 1.60 | **carbohydrate derivative metabolic process** | 20 | 0.017 | 1.45 | **lactose metabolic process** |
| 21 | 0.067 | 1.63 | transport | 21 | 0.017 | 1.45 | **D-tagatose 6-phosphate catabolic process** |

| | | | | | | | |
|---|---|---|---|---|---|---|---|
| 22 | 0.066 | 2.58 | pathogenesis | 22 | 0.017 | 1.45 | **positive regulation of catalytic activity** |
| 23 | 0.064 | 2.16 | pyrimidine-containing compound metabolic process | | | | |
| 24 | 0.061 | 2.17 | **pyrimidine-containing compound biosynthetic process** | | | | |
| 25 | 0.061 | 2.43 | macromolecule localization | | | | |
| 26 | 0.051 | 1.54 | organic substance transport | | | | |
| 27 | 0.050 | 1.91 | **cell wall macromolecule biosynthetic process** | | | | |
| 28 | 0.047 | 1.44 | **cellular component biogenesis** | | | | |
| 29 | 0.047 | 1.83 | **peptidoglycan-based cell wall biogenesis** | | | | |
| 30 | 0.044 | 1.95 | **peptidoglycan biosynthetic process** | | | | |
| 31 | 0.042 | 2.22 | pyrimidine nucleobase biosynthetic process | | | | |
| 32 | 0.039 | 1.51 | **carbohydrate catabolic process** | | | | |
| 33 | 0.035 | 1.47 | **nitrogen compound transport** | | | | |
| 34 | 0.034 | 1.49 | **generation of precursor metabolites and energy** | | | | |
| 35 | 0.030 | 1.59 | **nucleoside diphosphate metabolic process** | | | | |
| 36 | 0.030 | 1.59 | nucleobase metabolic process | | | | |
| 37 | 0.028 | 1.63 | **protein transport** | | | | |
| 38 | 0.025 | 1.52 | **cellular localization** | | | | |
| 39 | 0.022 | 1.42 | **glycolytic process** | | | | |
| 40 | 0.021 | 1.48 | **intracellular transport** | | | | |
| 41 | 0.020 | 1.32 | **ribonucleoprotein complex assembly** | | | | |
| 42 | 0.020 | 1.32 | **anion transport** | | | | |
| 43 | 0.018 | 1.35 | **ribosomal large subunit assembly** | | | | |
| 44 | 0.018 | 1.35 | **cellular protein localization** | | | | |
| 45 | 0.017 | 1.50 | 'de novo' pyrimidine nucleobase biosynthetic process | | | | |
| 46 | 0.017 | 1.50 | **sulfur compound transport** | | | | |
| 47 | 0.015 | 1.33 | **establishment of protein localization to membrane** | | | | |

123

**Table S10** Significantly enriched UniProt keywords of downregulated proteins in Δ*clpP2* and Δ*clpP12* whole proteomes compared to WT at 42 °C identified with aGOtool (Schölz et al., 2015). Terms unique for either whole proteome are highlighted in bold.

| **ΔclpP2/WT, 42 °C** | | | | **ΔclpP12/WT, 42 °C** | | | |
|---|---|---|---|---|---|---|---|
| **Rank** | **s value** | **-log₁₀ (p-value)** | **Description (Keywords)** | **Rank** | **s value** | **-log₁₀ (p-value)** | **Description (Keywords)** |
| 1 | 0.649 | 7.21 | **Ribosomal protein** | 1 | 0.667 | 6.04 | Signal |
| 2 | 0.306 | 3.30 | Membrane | 2 | 0.251 | 2.68 | Membrane |
| 3 | 0.260 | 3.08 | **Transmembrane** | 3 | 0.232 | 2.64 | Transmembrane helix |
| 4 | 0.253 | 3.76 | Signal | 4 | 0.221 | 3.07 | Transport |
| 5 | 0.231 | 2.88 | Transmembrane helix | 5 | 0.143 | 3.69 | Pyrimidine biosynthesis |
| 6 | 0.219 | 3.35 | Transport | 6 | 0.050 | 2.03 | Virulence |
| 7 | 0.116 | 2.80 | **rRNA-binding** | 7 | 0.049 | 2.22 | **Peptidoglycan-anchor** |
| 8 | 0.111 | 3.54 | Pyrimidine biosynthesis | 8 | 0.044 | 2.05 | **Cell wall** |
| 9 | 0.088 | 1.94 | **Cell membrane** | 9 | 0.040 | 1.71 | **Lipoprotein** |
| 10 | 0.076 | 1.90 | **RNA-binding** | 10 | 0.040 | 1.71 | **Secreted** |
| 11 | 0.066 | 2.58 | Virulence | 11 | 0.024 | 1.51 | **Serine protease** |
| 12 | 0.041 | 1.85 | **Cell wall biogenesis/degradation** | 12 | 0.017 | 1.45 | **Lactose metabolism** |
| 13 | 0.025 | 1.41 | **Cell shape** | 13 | 0.017 | 1.45 | **Phosphopantetheine** |
| 14 | 0.022 | 1.42 | **Peptidoglycan synthesis** | | | | |
| 15 | 0.021 | 1.48 | **Protein transport** | | | | |

**Table S11** Putative interaction partners ($log_2$ ratio ≥2 and $-log_{10}$ t-test p-value ≥1.3 in the XL-co-IP experiment, $log_2$ ratio <1 or $-log_{10}$ t-test p-value <1.3 in the whole proteome experiment) of ClpP1 and ClpP2 at 37 °C.

| | | **XL-co-IP of ΔclpP2** | | **Whole proteome of ΔclpP1** | | **XL-co-IP of ΔclpP1** | | **Whole proteome of ΔclpP2** | |
|---|---|---|---|---|---|---|---|---|---|
| **Gene name** | **Uniprot ID** | **$log_2$ ratio (anti-ClpP/isotype control)** | **$-log_{10}$ t-test p-value (anti-ClpP/ isotype control)** | **$log_2$ ratio (Δ*clpP1*/WT)** | **$-log_{10}$ t-test p-value (Δ*clpP1*/WT)** | **$log_2$ ratio (anti-ClpP/isotype control)** | **$-log_{10}$ t-test p-value (anti-ClpP/ isotype control)** | **$log_2$ ratio (Δ*clpP2*/WT)** | **$-log_{10}$ t-test p-value (Δ*clpP2*/WT)** |
| **Putative ClpP1 interactors** | | | | | | | | | |
| cinA;lmo1397 | Q8Y793 | 8.24 | 6.19 | 0.03 | 0.12 | 0.72 | 0.29 | -0.41 | 2.95 |
| mfd | Q8YAD0 | 7.08 | 4.46 | 0.01 | 0.03 | 0.54 | 0.16 | -0.21 | 1.58 |
| mcsB;lmo0231 | Q48759 | 6.46 | 7.02 | 0.12 | 0.90 | 0.00 | NaN | 2.60 | 8.86 |
| polC;lmo1320 | Q8Y7G1 | 6.15 | 4.55 | -0.03 | 0.15 | 2.29 | 1.82 | 1.20 | 7.57 |
| lmo2755 | Q8Y3S6 | 5.98 | 6.11 | 0.12 | 0.34 | 3.85 | 3.18 | 1.25 | 5.30 |
| purL;lmo1769 | Q8Y6C1 | 5.12 | 5.57 | 0.05 | 0.33 | 0.61 | 0.88 | 0.89 | 5.58 |
| dltA;lmo0974 | Q8Y8D4 | 5.11 | 4.79 | -0.01 | 0.01 | 1.20 | 1.69 | 1.54 | 4.50 |
| hemA;lmo1557 | Q8Y6X4 | 5.00 | 6.00 | 0.41 | 0.39 | 0.00 | NaN | 4.07 | 5.27 |
| lmo2182 | Q8Y587 | 4.93 | 3.98 | -0.19 | 0.14 | 2.81 | 1.76 | 1.16 | 1.87 |
| lmo0640 | Q8Y993 | 4.87 | 2.97 | 0.28 | 0.36 | 4.18 | 4.33 | 2.03 | 4.21 |

| | | | | | | | | | |
|---|---|---|---|---|---|---|---|---|---|
| murC;lmo1605 | Q8Y6S8 | 4.82 | 2.20 | 0.10 | 0.45 | 3.96 | 4.89 | 1.15 | 6.95 |
| lmo1358 | Q92BZ6 | 4.81 | 4.13 | NaN | NaN | 1.38 | 1.58 | NaN | NaN |
| lmo0537 | Q8Y9J1 | 4.77 | 1.48 | -0.04 | 0.20 | 3.32 | 0.74 | 1.57 | 7.43 |
| lmo1713 | Q8Y6H3 | 4.71 | 5.50 | 0.00 | 0.00 | 0.00 | NaN | 1.23 | 5.30 |
| lmo0227 | Q8YAB9 | 4.61 | 4.78 | 0.17 | 0.30 | 0.00 | NaN | 4.47 | 9.72 |
| lmo1737 | Q8Y6F0 | 4.57 | 4.37 | 0.10 | 0.39 | 1.81 | 2.38 | -0.55 | 3.91 |
| glpD | Q8Y7I4 | 4.54 | 3.75 | 0.04 | 0.02 | 0.63 | 0.22 | 2.33 | 3.06 |
| lmo0823 | Q8Y8S1 | 4.51 | 3.42 | 0.22 | 1.58 | 4.73 | 5.68 | 1.57 | 9.49 |
| lmo1881 | Q8Y621 | 4.37 | 5.60 | 0.08 | 0.11 | 2.25 | 1.30 | 1.04 | 2.65 |
| lmo1932 | Q8Y5X2 | 4.35 | 3.57 | 0.13 | 0.22 | 2.75 | 3.01 | 1.17 | 3.49 |
| lmo1667 | Q8Y6L7 | 4.21 | 5.15 | 0.06 | 0.30 | -0.19 | 0.09 | 0.53 | 2.76 |
| murB;lmo1420 | Q8Y776 | 4.19 | 5.02 | -0.09 | 0.10 | 0.00 | NaN | -0.06 | 0.08 |
| tal1;lmo2743 | Q8Y3T8 | 4.18 | 1.83 | 0.10 | 0.22 | 5.19 | 3.07 | 1.24 | 4.16 |
| lmo2215 | Q8Y561 | 4.17 | 5.80 | 0.07 | 0.41 | 1.72 | 2.09 | 0.93 | 7.65 |
| lmo1363 | Q8Y7C2 | 4.15 | 3.95 | -0.07 | 0.19 | 1.28 | 0.80 | 0.15 | 0.64 |
| lmo0728 | Q8Y914 | 4.14 | 6.29 | -0.01 | 0.01 | 0.00 | NaN | 1.38 | 3.85 |
| lmo1384 | Q8Y7A4 | 3.98 | 4.24 | 0.09 | 0.15 | 0.00 | NaN | 5.21 | 9.55 |
| lmo0052 | Q8YAR3 | 3.98 | 5.12 | 0.04 | 0.17 | 0.78 | 0.36 | -0.15 | 1.09 |
| lmo0597 | Q8Y9D3 | 3.97 | 4.38 | 0.62 | 1.79 | 1.12 | 0.52 | 1.77 | 6.45 |
| lmo1009 | Q8Y8A3 | 3.94 | 5.03 | -0.08 | 0.12 | 0.38 | 0.20 | 0.20 | 0.43 |
| pflB | Q8Y786 | 3.92 | 4.11 | 0.00 | 0.00 | 3.69 | 3.15 | 2.81 | 8.02 |
| lmo2815 | Q8Y3M0 | 3.91 | 5.09 | 0.09 | 0.62 | 1.01 | 0.91 | 1.40 | 7.28 |
| lmo0305 | Q8YA56 | 3.88 | 2.95 | 0.03 | 0.06 | 0.19 | 0.10 | 0.72 | 2.78 |
| lmo1576 | Q8Y6V5 | 3.86 | 5.28 | 0.11 | 0.92 | 0.62 | 0.31 | 0.10 | 2.04 |
| metE;lmo1681 | Q8Y6K3 | 3.84 | 3.98 | 0.01 | 0.05 | 0.73 | 0.34 | 0.91 | 7.71 |
| fbp;lmo0830 | Q8Y8R5 | 3.81 | 6.40 | -0.06 | 0.04 | 0.00 | NaN | 5.12 | 9.40 |
| lmo0152 | Q8YAH4 | 3.80 | 7.59 | 0.17 | 0.36 | 0.30 | 0.28 | -1.45 | 5.14 |
| lmo1077 | Q8Y841 | 3.78 | 2.77 | 0.03 | 0.13 | 1.82 | 1.23 | -0.62 | 5.42 |
| lmo0234 | Q48762 | 3.75 | 4.17 | 0.02 | 0.09 | 2.14 | 1.12 | -0.42 | 3.05 |
| lmo2712 | Q8Y3W7 | 3.73 | 5.00 | 0.19 | 1.81 | -0.21 | 0.20 | 0.36 | 1.69 |
| lmo1400 | Q8Y792 | 3.73 | 5.50 | -0.38 | 0.25 | 0.00 | NaN | 2.66 | 3.47 |
| pflA | Q8Y5Y6 | 3.71 | 4.84 | 0.19 | 0.39 | 0.00 | NaN | 1.32 | 4.28 |
| tyrA | Q8Y5X9 | 3.69 | 3.11 | 0.03 | 0.17 | 1.92 | 1.41 | -0.54 | 3.33 |
| lmo2844 | Q8Y3J1 | 3.62 | 4.37 | -0.02 | 0.07 | 0.41 | 0.26 | -0.07 | 0.20 |
| sepF;lmo2030 | Q8Y5M7 | 3.60 | 4.17 | -0.01 | 0.02 | 0.00 | NaN | 4.00 | 11.78 |
| lmo1582 | Q8Y6U9 | 3.59 | 3.93 | 0.15 | 0.38 | 1.74 | 2.76 | 0.89 | 3.82 |
| parB | Q8Y3P4 | 3.58 | 4.84 | 0.02 | 0.11 | 1.74 | 0.61 | -0.21 | 1.29 |
| lmo2643 | Q8Y431 | 3.57 | 4.04 | 0.18 | 0.59 | 1.42 | 0.83 | 0.21 | 0.89 |
| uvrA;lmo2488 | Q8Y4F6 | 3.49 | 1.96 | 0.09 | 0.81 | 1.21 | 1.27 | 0.69 | 5.55 |
| rpsU;lmo1469 | P0DJP1 | 3.48 | 5.21 | -0.13 | 0.14 | 0.00 | NaN | -0.86 | 2.87 |
| lmo1078 | Q8Y840 | 3.48 | 3.13 | -0.17 | 0.48 | 1.29 | 0.48 | -0.56 | 3.04 |
| lmo1066 | Q8Y852 | 3.46 | 4.23 | -0.05 | 0.09 | 1.52 | 1.45 | -0.57 | 2.19 |
| lmo2452 | Q8Y4I7 | 3.42 | 4.43 | 0.39 | 1.42 | 0.00 | NaN | 2.61 | 6.91 |
| dacA;cdaA;lmo2120 | Q8Y5E4 | 3.42 | 4.84 | 0.07 | 0.14 | 2.19 | 1.24 | -0.31 | 0.86 |
| lmo1979 | Q8Y5S6 | 3.39 | 3.92 | 0.11 | 0.19 | 0.10 | 0.07 | 0.42 | 1.24 |
| gcvT;lmo1348 | Q8Y7D5 | 3.36 | 4.69 | 0.01 | 0.03 | 1.47 | 0.86 | 0.51 | 5.01 |
| lmo1634 | Q8Y6Q0 | 3.33 | 3.57 | -0.01 | 0.02 | 0.00 | NaN | -0.55 | 0.82 |
| dnaC | Q92FQ6 | 3.33 | 5.03 | 0.09 | 0.68 | 0.44 | 0.32 | 0.52 | 2.85 |
| lmo1866 | Q8Y634 | 3.33 | 4.96 | 0.02 | 0.06 | 0.06 | 0.05 | 1.06 | 6.25 |

| | | | | | | | | | |
|---|---|---|---|---|---|---|---|---|---|
| cshB;lmo1450 | Q8Y755 | 3.32 | 3.76 | -0.18 | 1.50 | 1.30 | 1.08 | -0.71 | 5.61 |
| lmo0785 | Q8Y8V7 | 3.29 | 3.58 | 0.03 | 0.10 | 0.00 | NaN | 1.53 | 6.73 |
| lmo0841 | Q8Y8Q5 | 3.29 | 3.11 | 0.06 | 0.10 | 1.73 | 1.63 | 0.33 | 1.26 |
| purM;lmo1767 | Q8Y6C3 | 3.27 | 3.34 | 0.17 | 0.51 | 1.66 | 1.05 | 0.55 | 2.17 |
| lmo0825 | Q8Y8R9 | 3.21 | 5.87 | -0.04 | 0.19 | 1.68 | 1.10 | -0.12 | 0.79 |
| lmo0261 | Q8YA94 | 3.21 | 5.61 | 0.12 | 1.05 | 0.00 | NaN | 0.51 | 3.23 |
| prs2;lmo0509 | Q8Y9L8 | 3.21 | 4.97 | 0.03 | 0.09 | 1.97 | 1.52 | -0.72 | 4.29 |
| lmo0556 | Q8Y9H2 | 3.19 | 4.19 | 0.00 | NaN | 0.00 | NaN | 5.39 | 9.72 |
| lmo2055 | Q8Y5K5 | 3.17 | 4.39 | -0.34 | 0.44 | 0.00 | NaN | 3.39 | 6.35 |
| alr;dal;lmo0886 | P0DJL8 | 3.17 | 5.71 | 0.03 | 0.08 | 0.00 | NaN | 0.44 | 3.49 |
| dnaX | Q8Y3X5 | 3.16 | 2.74 | -0.03 | 0.18 | 1.21 | 2.15 | -0.46 | 1.83 |
| lmo0554 | Q8Y9H4 | 3.16 | 4.94 | -0.07 | 0.19 | 1.61 | 1.14 | -0.48 | 2.43 |
| lmo2209 | Q8Y567 | 3.13 | 3.66 | 0.06 | 0.15 | 0.29 | 0.25 | 1.10 | 4.53 |
| lmo1710 | Q92AU7 | 3.12 | 2.85 | -0.13 | 0.39 | 0.51 | 0.22 | -1.90 | 4.57 |
| purD;lmo1764 | Q8Y6C6 | 3.08 | 3.70 | 0.08 | 0.32 | 1.33 | 1.32 | 0.16 | 0.91 |
| sul | Q8YAC2 | 3.07 | 3.64 | 0.05 | 0.15 | 0.84 | 0.41 | 0.11 | 0.42 |
| lipL;lmo2566 | Q8Y489 | 3.06 | 3.06 | 0.09 | 0.31 | 0.95 | 0.80 | -0.44 | 2.41 |
| ffh | Q8Y695 | 3.05 | 2.64 | 0.04 | 0.12 | 1.37 | 1.47 | 0.01 | 0.03 |
| nnrD;lmo1622 | Q8Y6R2 | 3.05 | 4.22 | 0.06 | 0.25 | 0.21 | 0.18 | 1.19 | 7.39 |
| murG;lmo2035 | Q8Y5M2 | 3.03 | 2.46 | -0.03 | 0.15 | 1.69 | 2.16 | -0.20 | 1.21 |
| lmo1822 | Q8Y677 | 3.02 | 4.52 | 0.05 | 0.16 | 1.92 | 1.95 | -0.28 | 1.21 |
| lmo0774 | Q8Y8W8 | 3.02 | 4.76 | -0.12 | 0.79 | 2.16 | 1.28 | -0.47 | 4.42 |
| lmo2657 | Q8Y420 | 3.00 | 3.03 | 0.03 | 0.28 | 1.38 | 1.32 | -0.42 | 4.37 |
| fhuC | Q8Y5U5 | 2.99 | 3.00 | 0.15 | 0.08 | 1.05 | 0.85 | 3.98 | 4.47 |
| buk;lmo1370 | Q8Y7B6 | 2.99 | 4.44 | 0.06 | 0.27 | 0.72 | 0.58 | 1.10 | 6.65 |
| lmo0010 | Q8YAV3 | 2.99 | 4.68 | -0.04 | 0.10 | 1.64 | 2.82 | 0.03 | 0.11 |
| lmo0942 | Q8Y8G2 | 2.97 | 4.74 | 0.04 | 0.09 | 1.72 | 2.32 | -0.35 | 1.40 |
| pepC;lmo2338 | O69192 | 2.96 | 4.73 | 0.11 | 0.51 | 1.93 | 2.36 | 1.82 | 7.86 |
| fhs;lmo1877 | Q8Y624 | 2.94 | 5.83 | -0.03 | 0.03 | 0.00 | NaN | 6.20 | 9.70 |
| lmo1949 | Q8Y5V6 | 2.91 | 3.87 | 0.11 | 0.39 | 1.55 | 2.72 | -0.17 | 0.52 |
| lmo0740 | Q8Y902 | 2.90 | 3.08 | -0.08 | 0.06 | 0.00 | NaN | 1.90 | 2.65 |
| lmo0397 | Q8Y9X1 | 2.88 | 4.90 | -0.23 | 0.31 | 0.97 | 0.53 | 0.10 | 0.14 |
| accD;lmo1573 | Q8Y6V8 | 2.88 | 3.69 | 0.04 | 0.08 | 1.51 | 0.88 | 0.18 | 0.88 |
| lmo1867 | Q8Y633 | 2.86 | 3.61 | 0.05 | 0.09 | 0.00 | NaN | 3.03 | 9.31 |
| murA2;murZ;lmo2552 | Q8Y4A2 | 2.86 | 3.15 | 0.06 | 0.19 | 1.63 | 1.83 | 0.18 | 1.06 |
| lmo2550 | Q7AP48 | 2.86 | 2.87 | 0.04 | 0.11 | 1.07 | 1.23 | -0.20 | 1.05 |
| lmo0663 | Q8Y970 | 2.84 | 3.86 | 0.05 | 0.24 | 1.55 | 2.32 | -0.07 | 0.48 |
| lmo0664 | Q8Y969 | 2.83 | 4.94 | -0.11 | 0.13 | 0.00 | NaN | 1.98 | 4.55 |
| addA;lmo2267 | Q8Y511 | 2.80 | 5.46 | 0.15 | 0.69 | 0.00 | NaN | 1.30 | 6.79 |
| lmo0277 | Q8YA82 | 2.79 | 4.51 | 0.02 | 0.06 | 1.60 | 1.92 | 0.51 | 2.62 |
| ispE;lmo0190 | Q8YAE1 | 2.74 | 2.86 | 0.10 | 0.44 | 0.34 | 0.26 | 0.18 | 0.43 |
| dxr;lmo1317 | Q8Y7G4 | 2.74 | 4.08 | -0.07 | 0.34 | -0.05 | 0.03 | 0.13 | 0.53 |
| lmo1966 | Q8Y5T9 | 2.74 | 4.90 | -0.07 | 0.12 | 1.58 | 1.05 | -0.39 | 1.09 |
| pyrB;lmo1838 | Q8Y662 | 2.74 | 3.53 | 0.00 | NaN | 1.07 | 1.36 | 0.28 | 0.43 |
| prsA2;lmo2219 | Q8Y557 | 2.73 | 3.40 | -0.03 | 0.15 | 1.76 | 0.79 | -0.23 | 1.60 |
| dapA;lmo1435 | Q8Y766 | 2.72 | 2.98 | 0.09 | 0.22 | 0.56 | 0.60 | -0.48 | 2.36 |
| parE | Q8Y7J1 | 2.71 | 4.66 | -0.04 | 0.13 | 0.83 | 0.50 | -0.56 | 3.99 |
| lmo2565 | Q8Y490 | 2.67 | 3.55 | 0.05 | 0.13 | 1.12 | 0.53 | -0.26 | 0.94 |
| lmo2374 | Q8Y4R0 | 2.66 | 4.22 | 0.08 | 0.19 | 0.00 | NaN | -0.19 | 1.02 |

| | | | | | | | | | |
|---|---|---|---|---|---|---|---|---|---|
| rpmJ;lmo2609 | P66290 | 2.65 | 1.81 | NaN | NaN | 0.00 | NaN | NaN | NaN |
| lexA;lmo1302 | Q8Y7H7 | 2.65 | 2.80 | 0.27 | 0.53 | 2.58 | 2.67 | 1.51 | 4.13 |
| lmo0898 | Q8Y8K3 | 2.64 | 3.85 | -0.01 | 0.05 | 0.82 | 0.58 | 0.39 | 3.57 |
| lmo1647 | Q8Y6N7 | 2.63 | 3.11 | 0.12 | 0.35 | 0.09 | 0.07 | -0.06 | 0.11 |
| lmo1722 | Q8Y6G5 | 2.61 | 3.52 | -0.03 | 0.27 | 0.00 | NaN | -0.49 | 4.40 |
| dnaJ;lmo1472 | P0DJM1 | 2.61 | 1.59 | -0.01 | 0.04 | 1.72 | 0.89 | -0.09 | 0.24 |
| lmo0229 | Q7AP89 | 2.59 | 2.90 | 0.17 | 0.72 | 1.71 | 1.52 | 2.77 | 9.08 |
| recD2;lmo1509 | Q8Y717 | 2.57 | 3.91 | 0.31 | 0.68 | 0.00 | NaN | 2.34 | 6.32 |
| lmo1515 | Q8Y711 | 2.57 | 4.00 | 0.15 | 0.58 | 3.25 | 4.95 | 1.42 | 4.70 |
| ruvB;lmo1532 | Q8Y6Z8 | 2.55 | 2.71 | -0.07 | 0.19 | 0.07 | 0.03 | 0.90 | 3.52 |
| topA | Q8Y7K2 | 2.55 | 1.98 | 0.06 | 0.45 | 0.36 | 0.32 | 0.12 | 0.86 |
| tmk;lmo2693 | Q8Y3Y6 | 2.54 | 3.10 | 0.16 | 0.45 | 1.47 | 3.32 | 0.02 | 0.05 |
| lmo1825 | Q8Y674 | 2.52 | 3.22 | -0.03 | 0.15 | 1.91 | 1.23 | -0.32 | 2.76 |
| obg;lmo1537 | Q8Y6Z3 | 2.51 | 3.14 | -0.05 | 0.20 | 0.95 | 0.83 | 0.14 | 0.67 |
| msrB;lmo1859 | Q8Y641 | 2.46 | 2.78 | 0.11 | 0.22 | 0.65 | 0.39 | -0.05 | 0.11 |
| lmo0964 | Q8Y8E0 | 2.46 | 4.00 | 0.10 | 0.15 | 0.00 | NaN | 4.44 | 8.81 |
| pheA;lmo1536 | Q8Y6Z4 | 2.45 | 5.02 | 0.02 | 0.04 | 0.92 | 1.08 | -1.02 | 4.71 |
| gbuB | Q7AP75 | 2.45 | 2.53 | 0.19 | 0.55 | 1.92 | 2.60 | -1.09 | 0.62 |
| fmt;lmo1823 | Q8Y676 | 2.44 | 3.87 | -0.09 | 0.45 | 1.77 | 2.29 | -0.60 | 4.52 |
| lmo2705 | Q8Y3X4 | 2.42 | 2.56 | 0.24 | 0.93 | 0.85 | 0.88 | -0.23 | 1.18 |
| cydC | Q8Y3W3 | 2.40 | 4.26 | 0.04 | 0.14 | 1.86 | 1.97 | -0.37 | 0.74 |
| lmo0291 | Q8YA68 | 2.37 | 4.38 | 0.36 | 0.90 | 0.00 | NaN | 5.79 | 11.19 |
| lmo1813 | Q8Y684 | 2.35 | 3.19 | 0.20 | 0.63 | 0.02 | 0.01 | 2.27 | 7.02 |
| lmo1131 | Q8Y7Y8 | 2.34 | 3.39 | 0.07 | 0.09 | 1.69 | 2.61 | 0.00 | 0.01 |
| proB;lmo1260 | Q93Q56 | 2.33 | 3.70 | 0.18 | 0.37 | 0.52 | 0.47 | -0.88 | 3.94 |
| lmo1281 | Q8Y7J6 | 2.32 | 3.75 | 0.24 | 0.66 | 0.32 | 0.36 | 0.42 | 1.35 |
| lmo1351 | Q8Y7D2 | 2.32 | 2.94 | -0.06 | 0.14 | 1.50 | 2.41 | -0.96 | 3.56 |
| tagH;lmo1075 | Q8Y843 | 2.32 | 3.75 | 0.07 | 0.19 | 0.90 | 0.64 | -0.12 | 0.42 |
| lmo1940 | Q8Y5W5 | 2.31 | 3.99 | 0.06 | 0.13 | 1.64 | 2.35 | -0.84 | 3.42 |
| nusG | Q8YAA6 | 2.27 | 3.55 | 0.10 | 0.34 | 1.20 | 0.47 | -0.61 | 3.44 |
| trpS;lmo2198 | Q8Y577 | 2.26 | 3.43 | -0.06 | 0.10 | 0.20 | 0.14 | -0.29 | 0.64 |
| prfA;lmo0200 | P22262 | 2.25 | 3.36 | 0.04 | 0.20 | 1.22 | 3.67 | 0.71 | 5.10 |
| lmo2317 | Q8Y4W2 | 2.23 | 2.75 | NaN | NaN | 0.00 | NaN | NaN | NaN |
| lmo0131 | Q8YAJ4 | 2.22 | 2.39 | 0.03 | 0.16 | 0.00 | NaN | 1.46 | 7.78 |
| mntH;lmo1424 | Q8Y773 | 2.22 | 2.61 | 0.10 | 0.26 | 1.18 | 0.61 | -1.75 | 6.20 |
| ecfA2;cbiO2;lmo2600 | Q8Y455 | 2.21 | 4.33 | 0.14 | 0.81 | 1.90 | 2.04 | 0.01 | 0.03 |
| lmo1930 | Q8Y5X3 | 2.20 | 2.60 | -0.12 | 0.18 | 1.40 | 3.54 | 0.34 | 0.98 |
| prmC;lmo2542 | Q8Y4A9 | 2.20 | 3.81 | 0.01 | 0.03 | 0.00 | NaN | -0.70 | 3.83 |
| lmo0525 | Q8Y9K2 | 2.18 | 1.71 | 0.17 | 1.29 | -0.29 | 0.18 | 0.96 | 6.37 |
| lmo0580 | Q8Y9E9 | 2.18 | 1.88 | -0.12 | 0.14 | 0.00 | NaN | 1.01 | 1.61 |
| lmo2422 | Q8Y4L5 | 2.18 | 2.64 | 0.00 | 0.01 | 0.52 | 0.37 | -0.41 | 1.94 |
| trpC;lmo1630 | Q8Y6Q4 | 2.17 | 3.50 | -0.12 | 0.23 | 1.55 | 1.68 | 0.13 | 0.25 |
| lmo1645 | Q8Y6N9 | 2.17 | 3.32 | 0.07 | 0.29 | -0.38 | 0.35 | 0.45 | 4.09 |
| pcrA | Q8Y6C9 | 2.17 | 3.24 | 0.12 | 1.14 | 0.79 | 0.63 | -0.21 | 2.27 |
| lmo2590 | Q8Y465 | 2.16 | 2.72 | 0.23 | 1.04 | 0.00 | NaN | 1.27 | 6.52 |
| topB | Q8Y3S5 | 2.16 | 3.90 | 0.05 | 0.20 | 0.00 | NaN | 0.37 | 2.76 |
| def;lmo1051 | Q8Y866 | 2.15 | 2.40 | 0.13 | 0.25 | 1.99 | 2.87 | -0.92 | 3.39 |
| rsbU | Q8Y8K7 | 2.14 | 1.93 | -0.14 | 0.50 | 0.00 | NaN | -0.36 | 1.73 |
| relA | Q8Y706 | 2.14 | 3.23 | -0.02 | 0.05 | 0.39 | 0.33 | -0.69 | 3.22 |

| | | | | | | | | | |
|---|---|---|---|---|---|---|---|---|---|
| lmo1507 | Q8Y719 | 2.13 | 3.27 | 0.05 | 0.48 | 0.00 | NaN | 0.09 | 0.57 |
| aroB;lmo1927 | Q8Y5X6 | 2.13 | 4.30 | -0.05 | 0.16 | 1.28 | 1.94 | -0.43 | 2.65 |
| metN2;lmo2419 | Q8Y4L8 | 2.08 | 3.88 | -0.01 | 0.02 | 0.71 | 0.53 | -0.67 | 4.34 |
| gcvPB;lmo1350 | Q8Y7D3 | 2.08 | 1.69 | 0.01 | 0.03 | 2.20 | 2.84 | 1.39 | 7.33 |
| lmo1910 | Q8Y5Z3 | 2.08 | 2.27 | 0.10 | 0.34 | 0.00 | NaN | 0.01 | 0.04 |
| clpB;lmo2206 | Q8Y570 | 2.08 | 3.78 | 0.08 | 0.32 | 0.11 | 0.08 | 2.67 | 11.11 |
| cshA;lmo0866 | Q8Y8N0 | 2.08 | 2.19 | -0.03 | 0.15 | 2.18 | 0.96 | -0.63 | 5.18 |
| mogR;lmo0674 | P0DJO8 | 2.05 | 3.19 | 0.21 | 0.26 | 0.00 | NaN | -0.47 | 0.92 |
| gcvPA;lmo1349 | Q8Y7D4 | 2.04 | 2.65 | -0.03 | 0.10 | 1.41 | 0.92 | 1.35 | 7.24 |
| coaX;lmo0221 | Q8YAC5 | 2.03 | 2.07 | -0.09 | 0.20 | -0.54 | 0.65 | 0.41 | 1.93 |
| lmo0163 | Q8YAG3 | 2.01 | 3.91 | 0.29 | 1.16 | 0.07 | 0.05 | 1.92 | 7.60 |

**Putative ClpP1 and ClpP2 interactors**

| | | | | | | | | | |
|---|---|---|---|---|---|---|---|---|---|
| murD;lmo2036 | Q8Y5M1 | 6.73 | 5.90 | -0.03 | 0.12 | 4.36 | 2.43 | -0.40 | 2.19 |
| lmo0454 | Q8Y9R9 | 6.42 | 4.30 | 0.09 | 0.33 | 2.89 | 3.15 | 0.81 | 4.31 |
| trmFO;gid;lmo1276 | Q8Y7K1 | 6.23 | 5.54 | 0.13 | 1.02 | 4.05 | 5.14 | 0.68 | 5.44 |
| fabG | Q8Y690 | 6.20 | 6.11 | -0.11 | 0.49 | 5.31 | 2.64 | -0.15 | 0.59 |
| carB;pyrAB;lmo1835 | Q8Y665 | 5.99 | 3.85 | -0.15 | 0.22 | 6.30 | 2.34 | -1.33 | 3.62 |
| lmo2592 | Q8Y463 | 5.82 | 6.18 | 0.16 | 0.84 | 4.48 | 4.30 | -0.31 | 1.92 |
| lmo1067 | Q8Y851 | 5.79 | 5.23 | 0.06 | 0.33 | 4.15 | 2.72 | -0.88 | 6.56 |
| ychF;lmo2779 | Q926X1 | 5.78 | 4.62 | 0.04 | 0.24 | 4.28 | 2.11 | -0.97 | 5.41 |
| lmo1815 | Q92AJ3 | 5.76 | 6.46 | 0.17 | 0.29 | 3.97 | 2.82 | -1.11 | 3.16 |
| lmo1636 | Q8Y6P8 | 5.74 | 4.09 | -0.09 | 0.34 | 4.56 | 4.47 | 0.61 | 2.87 |
| metG;metS;lmo0177 | Q8YAF2 | 5.67 | 5.79 | 0.07 | 0.30 | 5.13 | 2.35 | -0.65 | 5.79 |
| recN | Q8Y7B8 | 5.64 | 4.32 | 0.02 | 0.07 | 4.39 | 4.69 | -0.04 | 0.19 |
| thrS;lmo1559 | Q8Y6X2 | 5.62 | 8.78 | 0.00 | 0.01 | 4.72 | 4.71 | -0.66 | 3.67 |
| gshAB;gshF;lmo2770 | Q8Y3R3 | 5.57 | 7.21 | 0.09 | 0.43 | 4.89 | 4.99 | 0.29 | 2.94 |
| lmo0319 | Q7AP84 | 5.55 | 6.23 | 0.02 | 0.11 | 4.43 | 5.05 | -0.49 | 3.47 |
| hemL2;gsaB;lmo1685 | Q8Y6J9 | 5.51 | 2.87 | 0.02 | 0.04 | 5.91 | 4.57 | -0.55 | 3.11 |
| lmo1235 | Q8Y7N9 | 5.46 | 6.66 | -0.04 | 0.27 | 3.98 | 2.94 | -0.23 | 3.26 |
| lmo1782 | Q8Y6A9 | 5.44 | 3.09 | 0.09 | 0.81 | 2.73 | 3.35 | 0.00 | 0.01 |
| glyS;lmo1458 | Q8Y754 | 5.40 | 4.53 | -0.03 | 0.18 | 5.52 | 3.00 | -0.32 | 2.77 |
| lmo2263 | Q8Y515 | 5.40 | 6.43 | -0.29 | 0.33 | 3.73 | 4.53 | 0.29 | 1.25 |
| murI;racE;lmo1237 | Q8Y7N7 | 5.39 | 7.76 | 0.02 | 0.07 | 5.37 | 4.91 | -1.04 | 5.54 |
| lmo0931 | Q8Y8H3 | 5.39 | 5.78 | 0.14 | 0.69 | 3.68 | 1.63 | 0.12 | 0.42 |
| lmo1081 | Q8Y837 | 5.38 | 4.47 | 0.05 | 0.21 | 3.08 | 2.06 | -0.22 | 1.26 |
| purH;lmo1765 | Q8Y6C5 | 5.33 | 4.80 | 0.13 | 0.71 | 3.65 | 3.55 | 0.24 | 1.63 |
| lmo0977 | Q8Y8D1 | 5.28 | 4.01 | 0.10 | 0.41 | 4.97 | 2.63 | -0.22 | 1.60 |
| lmo2677 | Q8Y401 | 5.28 | 6.10 | 0.11 | 0.39 | 2.56 | 2.44 | 0.21 | 0.81 |
| lmo2720 | Q8Y3W1 | 5.24 | 4.17 | 0.08 | 0.43 | 3.63 | 2.06 | 0.81 | 4.19 |
| lmo1726 | Q8Y6G1 | 5.24 | 2.68 | -0.04 | 0.14 | 5.15 | 2.98 | 0.04 | 0.31 |
| lmo0521 | Q8Y9K6 | 5.22 | 5.18 | 0.10 | 1.01 | 3.46 | 2.76 | 0.17 | 1.49 |
| murE;lmo2038 | Q8Y5L9 | 5.21 | 4.43 | 0.05 | 0.15 | 4.39 | 5.48 | 0.05 | 0.19 |
| lmo1436 | Q8Y765 | 5.20 | 2.67 | 0.03 | 0.08 | 4.25 | 1.83 | -0.36 | 1.97 |
| tsaD;gcp;lmo2075 | Q8Y5I7 | 5.18 | 3.83 | 0.05 | 0.07 | 3.45 | 3.60 | 0.90 | 2.85 |
| mreB | Q8Y6Y3 | 5.16 | 2.91 | 0.04 | 0.11 | 2.85 | 1.69 | -0.61 | 3.34 |
| cmk;lmo1939 | Q8Y5W6 | 5.15 | 4.82 | 0.11 | 1.50 | 4.75 | 2.97 | -0.96 | 5.18 |
| lmo1744 | Q8Y6E3 | 5.08 | 9.47 | -0.11 | 0.75 | 3.35 | 4.30 | -0.46 | 2.06 |
| queA;lmo1531 | Q8Y6Z9 | 5.08 | 3.94 | 0.15 | 0.88 | 3.19 | 4.47 | 0.08 | 0.44 |

| | | | | | | | | | |
|---|---|---|---|---|---|---|---|---|---|
| pheT;lmo1222 | Q8Y7Q1 | 5.07 | 4.46 | -0.07 | 0.39 | 4.51 | 3.94 | -0.68 | 4.97 |
| lmo2155 | Q8Y5B2 | 5.05 | 3.90 | 0.10 | 0.56 | 3.99 | 2.64 | -0.33 | 2.85 |
| lmo1652 | Q8Y6N2 | 5.05 | 4.35 | 0.12 | 0.39 | 4.91 | 3.85 | -0.84 | 3.42 |
| lmo1812 | Q8Y685 | 5.00 | 4.36 | 0.08 | 0.48 | 4.62 | 4.30 | 0.87 | 4.79 |
| minD | Q8Y6Y7 | 5.00 | 5.65 | 0.02 | 0.06 | 2.71 | 1.71 | -0.02 | 0.06 |
| mnmE;trmE;lmo2811 | Q8Y3M4 | 5.00 | 4.56 | 0.20 | 0.52 | 3.81 | 2.75 | -0.09 | 0.26 |
| ftsE | Q8Y4E0 | 5.00 | 6.01 | 0.05 | 0.09 | 3.05 | 2.55 | 0.34 | 1.37 |
| lmo1935 | Q8Y5X0 | 4.99 | 3.09 | 0.11 | 0.25 | 3.82 | 3.36 | 0.71 | 3.26 |
| tyrS;lmo1598 | Q8Y6T4 | 4.94 | 3.06 | 0.11 | 1.42 | 3.82 | 2.91 | -0.21 | 2.91 |
| nadK2;lmo1586 | P65770 | 4.91 | 7.38 | 0.04 | 0.18 | 4.13 | 3.47 | -0.13 | 0.96 |
| azoR1;lmo0611 | Q8Y9C1 | 4.90 | 2.49 | 0.11 | 0.41 | 5.26 | 3.68 | -0.30 | 1.33 |
| gyrB | Q8YAV7 | 4.88 | 2.93 | 0.06 | 0.26 | 4.20 | 4.30 | -0.37 | 2.93 |
| ecfA1;cbiO1;lmo2601 | Q8Y454 | 4.87 | 3.09 | 0.16 | 0.44 | 3.90 | 3.24 | 0.54 | 2.23 |
| lmo0773 | Q8Y8W9 | 4.85 | 5.17 | 0.22 | 0.59 | 2.90 | 2.69 | -0.62 | 3.29 |
| lmo1717 | Q8Y6G9 | 4.85 | 6.23 | 0.04 | 0.16 | 3.12 | 3.96 | -0.75 | 4.59 |
| lysS;lmo0228 | Q8YAB8 | 4.84 | 5.41 | 0.10 | 1.52 | 5.72 | 2.23 | -1.02 | 7.01 |
| lmo0267 | Q8YA92 | 4.82 | 4.28 | 0.21 | 0.88 | 4.82 | 4.77 | -0.48 | 2.92 |
| lmo1006 | Q8Y8A4 | 4.82 | 5.38 | 0.00 | 0.00 | 4.17 | 4.87 | -1.12 | 4.25 |
| alaS;lmo1504 | Q8Y722 | 4.78 | 3.93 | 0.07 | 0.23 | 5.19 | 2.52 | -0.88 | 4.98 |
| aroC;aroF;lmo1928 | Q8Y5X5 | 4.74 | 5.44 | -0.06 | 0.38 | 3.00 | 2.99 | -0.46 | 4.65 |
| lmo1080 | Q8Y838 | 4.73 | 5.60 | -0.03 | 0.20 | 2.46 | 2.33 | 0.04 | 0.25 |
| lmo0387 | Q8Y9Y0 | 4.70 | 2.82 | -0.09 | 0.20 | 4.95 | 2.90 | -0.60 | 1.78 |
| lmo1084 | Q8Y834 | 4.68 | 2.65 | 0.04 | 0.14 | 3.69 | 2.31 | -0.12 | 0.60 |
| recA;lmo1398 | P0DJP0 | 4.63 | 3.08 | 0.10 | 0.41 | 3.63 | 2.25 | -0.20 | 0.84 |
| lmo1863 | Q8Y637 | 4.63 | 5.04 | 0.04 | 0.12 | 3.71 | 4.15 | -0.39 | 2.22 |
| lmo1357 | Q8Y7C7 | 4.63 | 3.00 | 0.05 | 0.15 | 4.51 | 2.18 | -0.11 | 0.45 |
| lmo1401 | Q8Y791 | 4.61 | 5.68 | 0.04 | 0.16 | 2.26 | 1.68 | -0.61 | 4.79 |
| lmo2474 | Q8Y4G9 | 4.59 | 4.33 | 0.18 | 1.76 | 2.10 | 1.51 | 0.32 | 3.28 |
| lysA | Q8Y5V3 | 4.58 | 6.78 | 0.07 | 0.27 | 2.37 | 2.61 | -0.59 | 4.11 |
| lmo1820 | Q8Y679 | 4.58 | 4.31 | -0.04 | 0.26 | 3.19 | 2.22 | -0.43 | 3.54 |
| lmo1431 | Q8Y770 | 4.52 | 5.02 | 0.10 | 0.34 | 3.98 | 2.62 | -0.33 | 1.98 |
| lmo1372 | Q8Y7B4 | 4.51 | 3.60 | 0.02 | 0.11 | 6.13 | 3.15 | 0.22 | 1.39 |
| lmo0287 | Q8YA72 | 4.51 | 6.12 | -0.02 | 0.05 | 3.93 | 2.66 | -0.27 | 0.93 |
| lmo1457 | P67195 | 4.50 | 5.17 | -0.02 | 0.09 | 2.80 | 2.33 | 0.06 | 0.25 |
| pycA | Q8Y846 | 4.49 | 2.78 | -0.02 | 0.15 | 5.66 | 2.03 | 0.04 | 0.25 |
| lmo2114 | Q8Y5F0 | 4.49 | 5.53 | 0.10 | 0.48 | 3.31 | 1.88 | -0.75 | 4.63 |
| lmo1057 | Q8Y860 | 4.48 | 4.72 | 0.15 | 0.80 | 3.55 | 3.31 | 0.03 | 0.07 |
| atpG;lmo2530 | Q927W3 | 4.48 | 4.58 | 0.04 | 0.10 | 4.01 | 2.29 | -0.17 | 0.65 |
| asnS;lmo1896 | P58695 | 4.47 | 5.35 | -0.02 | 0.10 | 4.57 | 1.80 | -0.61 | 4.23 |
| rnr;lmo2449 | Q8Y4J0 | 4.46 | 4.36 | 0.16 | 0.76 | 3.54 | 4.10 | -0.65 | 6.19 |
| mutS;lmo1403 | Q8Y789 | 4.44 | 4.40 | 0.08 | 0.86 | 3.13 | 2.58 | -0.43 | 2.74 |
| qoxA | Q8YAV0 | 4.42 | 4.93 | -0.09 | 1.24 | 3.94 | 2.25 | 0.59 | 5.44 |
| nusA | Q8Y7F9 | 4.42 | 4.62 | 0.08 | 0.21 | 4.71 | 2.51 | -0.62 | 2.74 |
| proA;lmo1259 | Q93Q55 | 4.41 | 5.08 | 0.05 | 0.17 | 3.37 | 2.45 | -0.31 | 1.81 |
| lmo0132 | Q8YAJ3 | 4.39 | 4.49 | 0.07 | 0.49 | 2.39 | 2.19 | 0.90 | 5.40 |
| rex;lmo2072 | P60384 | 4.38 | 3.50 | 0.04 | 0.19 | 5.56 | 2.08 | -0.15 | 1.06 |
| lmo1389 | Q8Y7A1 | 4.34 | 3.99 | -0.07 | 0.29 | 3.39 | 4.66 | -0.02 | 0.11 |
| aspS;lmo1519 | Q8Y709 | 4.33 | 5.34 | -0.02 | 0.14 | 5.80 | 2.68 | -0.75 | 6.03 |
| hemB | Q8Y6X7 | 4.32 | 5.66 | 0.05 | 0.19 | 3.02 | 1.85 | 0.06 | 0.29 |

| | | | | | | | | | |
|---|---|---|---|---|---|---|---|---|---|
| lmo1258 | Q8Y7L7 | 4.30 | 5.31 | 0.06 | 0.16 | 2.22 | 2.36 | 0.52 | 2.79 |
| ackA1;lmo1581 | Q8Y6V0 | 4.29 | 6.35 | 0.10 | 0.19 | 4.12 | 2.28 | -0.23 | 0.82 |
| lmo1387 | Q8Y7A2 | 4.28 | 2.45 | -0.14 | 0.44 | 4.16 | 3.08 | 0.84 | 2.91 |
| alsS | Q8Y5Q0 | 4.27 | 2.27 | 0.00 | 0.01 | 2.73 | 1.79 | -2.59 | 7.76 |
| lmo2401 | Q8Y4N4 | 4.26 | 3.18 | 0.10 | 0.25 | 4.77 | 5.25 | -0.18 | 0.57 |
| lmo0965 | Q8Y8D9 | 4.25 | 3.66 | -0.05 | 0.17 | 3.21 | 2.28 | 0.64 | 3.34 |
| citB | Q8Y6P3 | 4.24 | 3.28 | 0.13 | 1.18 | 3.77 | 2.56 | 0.31 | 2.20 |
| lmo1513 | Q8Y713 | 4.23 | 4.12 | 0.08 | 0.24 | 2.55 | 2.21 | 0.33 | 1.20 |
| lmo1236 | Q8Y7N8 | 4.16 | 3.49 | -0.05 | 0.37 | 3.95 | 4.07 | -0.53 | 3.84 |
| resD | Q8Y5V7 | 4.15 | 3.51 | 0.01 | 0.04 | 4.77 | 4.73 | 0.37 | 1.98 |
| ansB | Q8Y6M1 | 4.12 | 4.97 | -0.03 | 0.08 | 3.79 | 2.41 | -0.45 | 2.35 |
| gpsA;lmo1936 | Q8Y5W9 | 4.11 | 2.70 | -0.06 | 0.22 | 4.25 | 3.17 | -0.76 | 4.69 |
| lmo1718 | Q8Y6G8 | 4.08 | 3.76 | 0.02 | 0.09 | 4.22 | 3.28 | -0.31 | 2.75 |
| ddl;ddlA;lmo0855 | Q8Y8P1 | 4.07 | 3.85 | 0.11 | 0.43 | 4.23 | 1.80 | -0.94 | 6.16 |
| lmo1414 | Q8Y782 | 4.07 | 1.58 | 0.04 | 0.17 | 5.15 | 2.20 | -0.36 | 2.33 |
| thyA;lmo1874 | Q8Y626 | 4.04 | 1.97 | 0.13 | 0.66 | 4.79 | 2.65 | 0.62 | 2.84 |
| tarJ;lmo1087 | Q8Y831 | 4.04 | 4.47 | 0.12 | 0.57 | 4.72 | 4.63 | 0.14 | 1.01 |
| prfC;lmo0988 | Q8Y8C0 | 4.04 | 4.22 | 0.07 | 0.25 | 4.41 | 4.05 | -0.97 | 5.05 |
| lmo0356 | Q8YA10 | 4.03 | 3.00 | -0.13 | 0.43 | 3.92 | 3.94 | 0.25 | 1.15 |
| lmo2372 | Q8Y4R2 | 4.02 | 3.31 | -0.03 | 0.12 | 4.39 | 1.95 | 0.58 | 4.16 |
| rpoZ;lmo1826 | Q8Y673 | 4.02 | 4.44 | 0.01 | 0.01 | 2.71 | 1.63 | -1.08 | 3.30 |
| polA | Q8Y6W6 | 4.00 | 2.41 | -0.02 | 0.08 | 4.65 | 5.14 | 0.29 | 1.85 |
| lmo2031 | Q8Y5M6 | 3.99 | 4.57 | -0.02 | 0.07 | 3.14 | 3.61 | -0.64 | 3.58 |
| parA | Q926W7 | 3.98 | 7.30 | 0.13 | 0.64 | 4.62 | 4.66 | 0.21 | 1.29 |
| dapF;lmo2018 | Q8Y5N9 | 3.97 | 3.78 | 0.24 | 0.74 | 3.28 | 3.95 | 0.97 | 3.77 |
| dat;daaA;lmo1619 | P0DJL9 | 3.97 | 3.65 | 0.03 | 0.09 | 4.50 | 2.41 | -0.66 | 3.39 |
| rnj;lmo1434 | Q8Y767 | 3.95 | 3.58 | 0.00 | 0.01 | 3.60 | 1.91 | -0.30 | 2.18 |
| RsbR | Q8Y8K9 | 3.93 | 4.16 | 0.09 | 0.18 | 4.50 | 2.61 | -0.41 | 1.55 |
| pyrH;smbA;lmo1313 | P65927 | 3.93 | 4.72 | 0.04 | 0.05 | 4.02 | 5.01 | -0.16 | 0.25 |
| lmo1283 | Q8Y7J4 | 3.90 | 4.17 | -0.03 | 0.15 | 5.38 | 5.18 | 0.47 | 5.68 |
| minC;lmo1545 | Q8Y6Y6 | 3.90 | 4.68 | -0.08 | 0.29 | 3.20 | 6.18 | 0.62 | 3.88 |
| tkt | Q8Y7H4 | 3.89 | 4.70 | -0.01 | 0.04 | 7.29 | 2.46 | -0.39 | 3.14 |
| lmo1919 | Q8Y5Y4 | 3.89 | 3.15 | 0.01 | 0.02 | 3.43 | 2.01 | 0.10 | 0.24 |
| lmo2262 | Q8Y516 | 3.88 | 3.13 | -0.01 | 0.03 | 3.22 | 3.95 | -0.04 | 0.24 |
| lepA;lmo1479 | Q8Y742 | 3.87 | 5.79 | 0.04 | 0.13 | 2.45 | 2.75 | -0.73 | 4.66 |
| ileS;lmo2019 | Q8Y5N8 | 3.87 | 3.61 | -0.04 | 0.35 | 4.43 | 2.46 | -0.32 | 2.24 |
| lmo1705 | Q8Y6H9 | 3.87 | 5.06 | 0.09 | 0.34 | 4.14 | 6.48 | -0.83 | 5.55 |
| mnmG;gidA;lmo2810 | Q8Y3M5 | 3.85 | 5.09 | 0.08 | 0.64 | 2.96 | 3.08 | 0.26 | 3.81 |
| aroE;lmo1490 | Q8Y733 | 3.84 | 3.65 | 0.00 | 0.00 | 2.95 | 2.03 | -0.55 | 4.34 |
| metK;lmo1664 | Q8Y6M0 | 3.84 | 1.72 | 0.03 | 0.08 | 4.68 | 4.56 | -0.47 | 2.23 |
| lmo1371 | Q8Y7B5 | 3.82 | 4.98 | 0.10 | 0.44 | 4.12 | 4.43 | 0.90 | 4.61 |
| menF | Q8Y6K8 | 3.81 | 4.51 | 0.13 | 0.91 | 2.32 | 1.99 | -0.32 | 2.68 |
| lmo2247 | Q8Y529 | 3.81 | 2.46 | 0.09 | 0.71 | 4.52 | 4.70 | 0.76 | 5.43 |
| purK | Q8Y6B7 | 3.79 | 3.78 | 0.17 | 0.55 | 4.35 | 5.06 | 0.36 | 1.35 |
| lmo1092 | Q8Y826 | 3.78 | 4.20 | -0.08 | 0.42 | 2.80 | 3.59 | 0.00 | 0.00 |
| lmo0110 | Q8YAK8 | 3.77 | 5.28 | 0.09 | 0.37 | 3.32 | 3.83 | 0.92 | 4.92 |
| lmo1992 | Q8Y5R4 | 3.76 | 3.41 | 0.12 | 0.34 | 4.20 | 2.16 | -1.85 | 6.96 |
| lmo2406 | Q8Y4N0 | 3.76 | 1.85 | 0.05 | 0.12 | 3.56 | 2.66 | 0.96 | 4.66 |
| lmo0653 | Q8Y980 | 3.76 | 5.51 | 0.00 | 0.01 | 3.16 | 1.67 | -0.77 | 2.18 |

| | | | | | | | | | |
|---|---|---|---|---|---|---|---|---|---|
| lmo2700 | Q8Y3X9 | 3.76 | 5.40 | 0.04 | 0.20 | 5.06 | 2.31 | 0.42 | 2.22 |
| parC | Q8Y7J0 | 3.75 | 4.58 | 0.06 | 0.29 | 2.45 | 3.65 | -0.57 | 4.59 |
| murF | Q8Y8P0 | 3.74 | 3.84 | 0.07 | 0.22 | 3.67 | 3.95 | -1.21 | 6.39 |
| lmo2390 | Q8Y4P5 | 3.73 | 3.52 | 0.14 | 1.04 | 6.25 | 5.31 | 0.19 | 2.00 |
| mutL;lmo1404 | Q8Y788 | 3.72 | 6.64 | -0.02 | 0.05 | 2.70 | 3.57 | -0.38 | 1.79 |
| mbl | Q8Y4C5 | 3.72 | 3.78 | 0.06 | 0.44 | 2.95 | 2.01 | -0.59 | 4.75 |
| lmo2824 | Q8Y3L1 | 3.71 | 4.56 | 0.02 | 0.11 | 5.04 | 3.43 | 0.39 | 3.16 |
| zwf;lmo1978 | Q8Y5S7 | 3.69 | 2.62 | 0.10 | 0.80 | 5.03 | 2.40 | -0.31 | 1.97 |
| pnp;lmo1331 | Q8Y7F1 | 3.69 | 1.36 | 0.09 | 1.07 | 2.73 | 1.96 | -0.20 | 3.38 |
| trpA;lmo1627 | Q8Y6Q7 | 3.68 | 4.78 | -0.15 | 0.16 | 2.89 | 3.41 | -0.26 | 0.38 |
| mnmA;trmU;lmo1512 | Q8Y714 | 3.67 | 4.07 | 0.07 | 0.58 | 3.81 | 4.08 | -0.15 | 1.19 |
| lmo1611 | Q8Y6S2 | 3.67 | 5.28 | -0.05 | 0.25 | 6.21 | 2.50 | -0.48 | 3.47 |
| der;engA;lmo1937 | Q8Y5W8 | 3.66 | 2.99 | -0.01 | 0.05 | 2.74 | 3.48 | -0.68 | 4.86 |
| pdxT;lmo2102 | Q8Y5G1 | 3.64 | 4.15 | 0.01 | 0.03 | 2.95 | 2.32 | -1.09 | 5.25 |
| lmo0935 | Q8Y8G9 | 3.63 | 5.81 | 0.04 | 0.09 | 2.40 | 4.11 | -0.32 | 1.61 |
| purA;lmo0055 | Q8YAR1 | 3.63 | 3.15 | 0.04 | 0.23 | 3.49 | 2.48 | -0.23 | 3.23 |
| hom | Q8Y4A4 | 3.62 | 4.82 | -0.08 | 0.30 | 5.47 | 2.29 | -1.23 | 6.01 |
| pyrR;lmo1840 | Q8Y660 | 3.62 | 4.53 | -0.03 | 0.10 | 3.56 | 2.28 | -0.59 | 2.77 |
| gmk;lmo1827 | Q8Y672 | 3.60 | 6.02 | 0.08 | 0.42 | 3.41 | 4.48 | -0.62 | 3.45 |
| lmo0906 | Q8Y8J5 | 3.60 | 3.54 | 0.04 | 0.19 | 3.59 | 6.86 | 0.53 | 2.81 |
| lmo2823 | Q8Y3L2 | 3.59 | 2.97 | 0.04 | 0.09 | 3.04 | 4.50 | 0.53 | 2.33 |
| lmo1855 | Q8Y645 | 3.58 | 3.96 | 0.01 | 0.02 | 3.10 | 3.78 | -0.80 | 3.79 |
| lmo2403 | Q8Y4N3 | 3.58 | 3.40 | -0.08 | 0.40 | 2.51 | 3.83 | -0.27 | 2.66 |
| argS;lmo2561 | Q8Y493 | 3.57 | 1.98 | 0.06 | 0.19 | 4.71 | 2.63 | -0.53 | 3.29 |
| gatB;lmo1754 | Q8Y6D3 | 3.54 | 2.08 | 0.05 | 0.29 | 2.81 | 1.71 | -0.21 | 2.36 |
| lmo2518 | Q8Y4D2 | 3.53 | 4.61 | 0.07 | 0.17 | 3.31 | 4.28 | -0.72 | 3.16 |
| apt;lmo1524 | P0A2X5 | 3.53 | 6.37 | 0.00 | 0.01 | 2.24 | 3.85 | -0.50 | 3.06 |
| fruB | Q8Y4U5 | 3.53 | 2.98 | 0.10 | 0.17 | 3.66 | 2.31 | -0.65 | 2.29 |
| folD;lmo1360 | Q8Y7C5 | 3.52 | 4.04 | 0.02 | 0.04 | 3.81 | 3.17 | 0.52 | 2.27 |
| nadE;lmo1093 | Q8Y825 | 3.51 | 1.53 | -0.02 | 0.14 | 4.67 | 1.85 | 0.08 | 0.46 |
| folA | Q8YAC1 | 3.51 | 3.56 | 0.29 | 1.07 | 3.29 | 3.62 | 0.02 | 0.05 |
| lmo0407 | Q8Y9W2 | 3.51 | 2.90 | 0.40 | 0.96 | 3.93 | 3.17 | 0.97 | 1.93 |
| ezrA;lmo1594 | Q8Y6T8 | 3.50 | 3.17 | -0.04 | 0.22 | 4.44 | 3.80 | -0.72 | 6.15 |
| secA1;lmo2510 | P47847 | 3.49 | 2.83 | -0.04 | 0.38 | 2.85 | 2.73 | -0.06 | 0.46 |
| rnj;lmo1027 | Q92CZ5 | 3.48 | 3.10 | -0.06 | 0.35 | 3.44 | 1.80 | -0.30 | 2.74 |
| lmo2831 | Q8Y3K4 | 3.47 | 3.53 | 0.08 | 0.34 | 3.58 | 3.86 | -0.34 | 2.57 |
| glmM;lmo2118 | Q8Y5E6 | 3.46 | 5.97 | 0.05 | 0.54 | 5.93 | 2.72 | -0.08 | 0.60 |
| lmo2411 | Q928M6 | 3.45 | 5.33 | 0.02 | 0.20 | 3.72 | 2.30 | 0.41 | 4.52 |
| lmo1423 | Q8Y774 | 3.44 | 1.99 | 0.03 | 0.20 | 2.82 | 1.99 | 0.01 | 0.06 |
| lmo1354 | Q8Y7C9 | 3.40 | 2.75 | -0.03 | 0.16 | 5.24 | 2.43 | 0.05 | 0.16 |
| pnp | Q8Y5V2 | 3.40 | 1.64 | 0.05 | 0.20 | 5.25 | 2.45 | -0.32 | 1.70 |
| ribC | Q8Y7F2 | 3.38 | 2.76 | -0.10 | 0.33 | 2.69 | 2.94 | 0.16 | 0.44 |
| lmo0970 | Q8Y8D5 | 3.36 | 3.75 | 0.06 | 0.24 | 5.28 | 1.98 | 0.14 | 0.84 |
| infB;lmo1325 | Q8Y7F6 | 3.34 | 1.50 | 0.01 | 0.07 | 4.07 | 2.29 | -0.40 | 5.25 |
| lmo2194 | Q8Y580 | 3.34 | 4.27 | -0.02 | 0.10 | 4.27 | 6.75 | -0.77 | 4.64 |
| lmo1922 | Q8Y5Y1 | 3.33 | 2.70 | -0.10 | 0.35 | 5.06 | 2.78 | -0.57 | 2.56 |
| lmo1068 | Q8Y850 | 3.32 | 3.74 | -0.01 | 0.05 | 3.77 | 1.83 | -1.56 | 6.61 |
| lisR | Q92BX8 | 3.31 | 5.20 | -0.01 | 0.01 | 2.63 | 2.05 | 0.17 | 0.84 |
| ftsA | Q8Y5M4 | 3.30 | 4.08 | 0.01 | 0.06 | 4.01 | 2.21 | -0.20 | 1.48 |

| | | | | | | | | | |
|---|---|---|---|---|---|---|---|---|---|
| trpD;lmo1631 | Q8Y6Q3 | 3.30 | 3.55 | -0.42 | 0.30 | 2.15 | 3.12 | 0.92 | 1.06 |
| dnaA;lmo0001 | Q8YAW2 | 3.29 | 5.24 | 0.07 | 0.31 | 2.22 | 2.46 | -0.18 | 1.50 |
| lmo1493 | Q8Y730 | 3.27 | 2.58 | 0.06 | 0.36 | 4.85 | 2.89 | -0.06 | 0.55 |
| lmo1886 | Q8Y616 | 3.26 | 2.12 | -0.06 | 0.33 | 5.57 | 2.69 | 0.11 | 0.78 |
| lmo2577 | Q8Y478 | 3.26 | 4.12 | 0.26 | 1.70 | 3.88 | 4.34 | -0.22 | 1.59 |
| lmo1814 | Q8Y683 | 3.25 | 1.53 | -0.06 | 0.19 | 4.42 | 2.92 | -0.49 | 2.91 |
| lmo1013 | Q8Y899 | 3.25 | 3.19 | 0.22 | 0.87 | 2.67 | 1.82 | -0.28 | 1.48 |
| uvrB;lmo2489 | Q8Y4F5 | 3.24 | 2.99 | 0.18 | 0.89 | 2.07 | 2.16 | 0.93 | 5.56 |
| lmo1339 | Q8Y7E4 | 3.22 | 2.31 | 0.00 | 0.00 | 2.79 | 2.97 | -0.39 | 1.85 |
| lmo1612 | Q8Y6S1 | 3.21 | 6.76 | -0.03 | 0.04 | 3.19 | 4.79 | 0.12 | 0.37 |
| azoR2;lmo0786 | Q8Y8V6 | 3.19 | 1.56 | 0.11 | 0.34 | 5.41 | 3.06 | -0.85 | 3.53 |
| lmo1005 | Q8Y8A5 | 3.19 | 1.76 | -0.08 | 0.54 | 2.57 | 2.80 | -0.08 | 0.41 |
| nagB;lmo0957 | Q8Y8E7 | 3.18 | 4.71 | 0.06 | 0.28 | 3.66 | 5.62 | -0.97 | 5.12 |
| dapB;lmo1907 | Q8Y5Z6 | 3.18 | 3.31 | 0.14 | 0.37 | 5.90 | 2.59 | -0.10 | 0.32 |
| tarI;lmo1086 | Q8Y832 | 3.17 | 4.31 | -0.03 | 0.17 | 4.08 | 2.39 | 0.01 | 0.02 |
| proC;lmo0396 | Q8Y9X2 | 3.16 | 3.29 | 0.03 | 0.12 | 2.28 | 1.48 | -0.30 | 1.30 |
| lmo1976 | Q8Y5S9 | 3.16 | 5.47 | -0.02 | 0.09 | 3.34 | 4.06 | -0.75 | 5.65 |
| panB;lmo1902 | Q8Y601 | 3.15 | 2.43 | 0.05 | 0.06 | 4.91 | 2.60 | -0.83 | 1.95 |
| lmo0848 | Q8Y8P8 | 3.14 | 4.89 | 0.11 | 0.73 | 2.75 | 4.74 | -0.32 | 1.47 |
| pdp | Q8Y5R3 | 3.13 | 2.07 | 0.01 | 0.02 | 3.36 | 3.88 | -0.03 | 0.11 |
| lmo0607 | Q8Y9C5 | 3.13 | 3.54 | -0.05 | 0.18 | 2.23 | 3.29 | -0.68 | 3.75 |
| plsX;lmo1809 | Q8Y688 | 3.13 | 3.35 | -0.06 | 0.15 | 3.45 | 4.08 | -0.40 | 1.97 |
| gltX;lmo0237 | Q8YAB3 | 3.13 | 1.90 | -0.05 | 0.25 | 4.46 | 1.68 | -0.18 | 1.60 |
| gatA;lmo1755 | Q8Y6D2 | 3.13 | 3.40 | 0.03 | 0.16 | 5.86 | 1.72 | -0.12 | 1.02 |
| hisS;lmo1520 | Q8Y708 | 3.09 | 2.63 | 0.08 | 0.34 | 3.48 | 1.97 | -0.49 | 4.48 |
| lmo1795 | Q8Y6A0 | 3.05 | 6.42 | 0.22 | 1.07 | 2.11 | 1.92 | -0.05 | 0.11 |
| prmA;lmo1471 | P0DJO9 | 3.04 | 1.93 | -0.06 | 0.25 | 3.82 | 3.19 | 0.00 | 0.00 |
| lmo1745 | Q8Y6E2 | 3.01 | 3.98 | 0.00 | 0.01 | 2.05 | 2.73 | -0.59 | 5.04 |
| yhaM;lmo2220 | Q8Y556 | 2.99 | 5.11 | 0.03 | 0.06 | 3.34 | 1.77 | 0.20 | 0.51 |
| smc | Q8Y692 | 2.99 | 3.91 | 0.05 | 0.16 | 2.79 | 2.86 | -0.12 | 0.43 |
| lmo1946 | Q8Y5V9 | 2.98 | 3.74 | -0.10 | 0.58 | 2.60 | 4.65 | 0.67 | 3.79 |
| menD;lmo1675 | Q8Y6K9 | 2.98 | 4.71 | 0.04 | 0.20 | 2.09 | 1.55 | -0.03 | 0.17 |
| lmo0271 | Q8YA88 | 2.97 | 3.25 | 0.07 | 0.45 | 4.06 | 3.91 | -0.01 | 0.05 |
| lmo1373 | Q8Y7B3 | 2.96 | 4.41 | 0.02 | 0.06 | 4.30 | 3.43 | 0.23 | 1.63 |
| leuS;lmo1660 | Q8Y6M4 | 2.94 | 3.51 | 0.06 | 0.28 | 4.84 | 4.65 | -0.43 | 3.19 |
| prfB;lmo2509 | Q8Y4D8 | 2.94 | 3.00 | 0.07 | 0.18 | 4.13 | 4.70 | -0.78 | 3.60 |
| rny;lmo1399 | P0DJP2 | 2.93 | 3.60 | 0.12 | 0.32 | 3.90 | 2.04 | -0.26 | 1.44 |
| lmo0983 | Q8Y8C5 | 2.91 | 5.26 | -0.18 | 0.33 | 2.75 | 3.16 | 0.98 | 4.36 |
| lmo2248 | Q929C7 | 2.85 | 3.47 | 0.02 | 0.12 | 3.63 | 3.81 | -0.86 | 6.74 |
| lmo0241 | Q92F34 | 2.84 | 3.20 | 0.11 | 0.36 | 3.58 | 2.12 | -0.26 | 1.57 |
| panC;lmo1901 | Q8Y602 | 2.82 | 2.92 | -0.02 | 0.02 | 2.03 | 3.27 | -0.70 | 1.62 |
| ligA;lmo1758 | Q8Y6D0 | 2.78 | 3.71 | 0.05 | 0.15 | 3.76 | 5.03 | -0.01 | 0.03 |
| rph;lmo1238 | Q8Y7N6 | 2.78 | 2.96 | -0.04 | 0.12 | 3.66 | 2.11 | -1.08 | 5.36 |
| asd;lmo1437 | Q8Y764 | 2.77 | 2.79 | 0.00 | 0.01 | 4.95 | 2.27 | -0.16 | 0.52 |
| lmo2537 | Q8Y4B4 | 2.77 | 2.68 | 0.04 | 0.14 | 3.82 | 4.81 | -0.53 | 3.29 |
| lmo1967 | Q8Y5T8 | 2.75 | 2.45 | 0.00 | 0.00 | 4.08 | 3.58 | -0.09 | 0.19 |
| carA;pyrAA;lmo1836 | Q8Y664 | 2.73 | 2.98 | 0.14 | 0.16 | 3.46 | 2.15 | -1.23 | 3.40 |
| lmo1753 | Q8Y6D4 | 2.72 | 3.03 | -0.13 | 0.33 | 2.64 | 3.29 | -0.75 | 3.47 |
| lmo2462 | Q8Y4H9 | 2.71 | 3.75 | -0.11 | 0.76 | 3.46 | 3.59 | -1.38 | 8.59 |

| | | | | | | | | | |
|---|---|---|---|---|---|---|---|---|---|
| lmo1595 | Q8Y6T7 | 2.71 | 1.56 | 0.27 | 0.54 | 3.37 | 4.03 | -0.11 | 0.37 |
| lmo2371 | Q8Y4R3 | 2.70 | 1.83 | 0.09 | 0.32 | 4.07 | 2.16 | 0.55 | 4.36 |
| lmo1254 | Q8Y7M0 | 2.69 | 1.92 | 0.08 | 0.38 | 4.36 | 2.89 | 0.74 | 4.38 |
| hslU;clpY;lmo1279 | Q8Y7J8 | 2.68 | 2.09 | -0.02 | 0.06 | 2.58 | 1.80 | -0.22 | 1.26 |
| rsbV;lmo0893 | P0A4J8 | 2.68 | 1.38 | -0.09 | 0.33 | 4.27 | 5.50 | -0.28 | 1.79 |
| lmo0553 | Q8Y9H5 | 2.66 | 5.07 | 0.01 | 0.03 | 4.94 | 2.05 | -0.12 | 0.33 |
| ldh1;ldh;lmo0210 | P33380 | 2.66 | 3.58 | 0.11 | 0.62 | 5.99 | 2.31 | -0.89 | 5.91 |
| comEB | Q8Y739 | 2.65 | 2.29 | -0.28 | 1.07 | 2.38 | 2.58 | -0.83 | 4.00 |
| lmo2358 | Q8Y4S4 | 2.64 | 2.81 | 0.02 | 0.05 | 2.71 | 1.80 | 0.10 | 0.30 |
| era;lmo1462 | Q8Y750 | 2.59 | 2.98 | -0.13 | 0.40 | 2.11 | 3.16 | -0.08 | 0.26 |
| gbuA | Q7AP76 | 2.58 | 2.50 | 0.06 | 0.30 | 3.84 | 2.57 | 0.70 | 6.58 |
| trpB;lmo1628 | Q8Y6Q6 | 2.57 | 5.26 | -0.08 | 0.10 | 2.39 | 3.59 | -0.16 | 0.30 |
| cysS;lmo0239 | Q8YAB1 | 2.57 | 3.45 | 0.12 | 0.55 | 5.21 | 2.01 | 0.39 | 2.29 |
| lmo2433 | Q8Y4K5 | 2.56 | 1.36 | -0.07 | 0.22 | 3.74 | 2.39 | -0.57 | 2.79 |
| rpoC;lmo0259 | Q8YA96 | 2.51 | 2.76 | 0.03 | 0.16 | 3.73 | 1.71 | -0.71 | 6.41 |
| xpt;lmo1885 | Q8Y617 | 2.50 | 4.25 | 0.10 | 0.15 | 3.77 | 3.77 | -0.88 | 4.00 |
| tagD | Q8Y829 | 2.49 | 2.63 | 0.11 | 0.53 | 2.23 | 2.66 | -0.28 | 0.65 |
| ccpA | Q8Y6T3 | 2.49 | 2.63 | -0.02 | 0.05 | 5.10 | 1.87 | -0.58 | 1.78 |
| lmo2426 | Q8Y4L1 | 2.48 | 2.11 | -0.04 | 0.11 | 2.49 | 4.76 | -0.49 | 1.46 |
| mtnN;lmo1494 | Q8Y729 | 2.47 | 2.45 | 0.05 | 0.18 | 3.48 | 4.49 | -0.20 | 1.61 |
| lmo1083 | Q8Y835 | 2.44 | 4.26 | 0.06 | 0.31 | 5.42 | 2.14 | 0.00 | 0.00 |
| ftsY;lmo1803 | Q8Y693 | 2.40 | 2.78 | 0.11 | 0.38 | 2.57 | 4.04 | -0.92 | 4.68 |
| lmo1392 | Q8Y798 | 2.40 | 2.83 | -0.12 | 0.46 | 3.35 | 2.61 | -0.13 | 0.48 |
| aroD;lmo0491 | Q8Y9N4 | 2.36 | 3.85 | 0.12 | 0.40 | 2.57 | 2.14 | -0.41 | 3.35 |
| lmo1467 | Q7AP63 | 2.36 | 3.18 | -0.01 | 0.03 | 2.93 | 3.54 | 0.24 | 0.79 |
| lmo0494 | Q8Y9N1 | 2.34 | 2.32 | -0.15 | 0.55 | 3.27 | 5.28 | -0.34 | 0.41 |
| lmo1257 | Q92CE7 | 2.33 | 2.99 | 0.15 | 0.67 | 3.31 | 3.67 | -1.84 | 6.12 |
| lmo2450 | Q8Y4I9 | 2.33 | 3.36 | 0.19 | 0.98 | 3.08 | 4.41 | -0.71 | 4.16 |
| argR;lmo1367 | Q8Y7B9 | 2.32 | 2.24 | 0.17 | 1.10 | 2.13 | 2.64 | 0.13 | 0.44 |
| lmo0608 | Q8Y9C4 | 2.30 | 3.11 | 0.04 | 0.22 | 2.15 | 3.74 | -0.42 | 3.38 |
| lmo1082 | Q8Y836 | 2.29 | 1.51 | -0.08 | 0.60 | 3.78 | 2.15 | -0.16 | 1.85 |
| lmo2487 | Q8Y4F7 | 2.28 | 2.93 | -0.18 | 1.41 | 3.21 | 2.72 | -0.72 | 5.05 |
| lmo2853 | Q8Y3I3 | 2.26 | 4.23 | -0.05 | 0.21 | 4.22 | 2.09 | -0.22 | 0.63 |
| lmo2208 | Q8Y568 | 2.25 | 3.24 | -0.03 | 0.29 | 2.26 | 2.48 | -0.86 | 8.13 |
| lmo1012 | Q8Y8A0 | 2.24 | 1.38 | -0.11 | 0.91 | 5.93 | 4.06 | 0.37 | 2.76 |
| psuG;lmo2340 | Q8Y4U2 | 2.23 | 3.74 | 0.10 | 0.23 | 5.72 | 2.66 | -0.21 | 0.66 |
| fumC;citG;lmo2225 | Q8Y551 | 2.23 | 1.46 | 0.08 | 0.30 | 3.49 | 6.26 | 0.00 | 0.01 |
| lmo1022 | Q8Y892 | 2.22 | 2.57 | 0.02 | 0.05 | 3.06 | 5.45 | -0.46 | 2.86 |
| lmo2515 | Q927X8 | 2.20 | 1.58 | -0.06 | 0.28 | 3.36 | 1.75 | -0.28 | 1.68 |
| lmo1217 | Q8Y7Q5 | 2.17 | 3.49 | 0.05 | 0.11 | 7.28 | 2.69 | 0.03 | 0.07 |
| lmo2389 | Q8Y4P6 | 2.15 | 2.72 | 0.03 | 0.20 | 2.98 | 1.34 | 0.27 | 2.89 |
| valS;lmo1552 | Q8Y6X9 | 2.13 | 2.22 | -0.05 | 0.36 | 5.53 | 3.12 | -0.51 | 5.36 |
| pheS;lmo1221 | Q8Y7Q2 | 2.13 | 3.36 | 0.05 | 0.18 | 2.77 | 3.40 | -0.61 | 4.08 |
| sigA;rpoD;lmo1454 | P52331 | 2.13 | 2.21 | 0.14 | 0.67 | 3.29 | 3.32 | -0.41 | 1.95 |
| lmo0847 | Q8Y8P9 | 2.12 | 2.78 | 0.03 | 0.18 | 2.82 | 1.62 | -0.55 | 5.48 |
| accA;lmo1572 | Q8Y6V9 | 2.11 | 2.36 | 0.06 | 0.29 | 3.35 | 3.69 | 0.35 | 2.75 |
| rpoB;lmo0258 | Q9RLT9 | 2.09 | 2.36 | 0.03 | 0.27 | 3.47 | 2.21 | -0.40 | 4.81 |
| lmo2089 | Q8Y5H3 | 2.07 | 3.73 | 0.08 | 0.52 | 4.18 | 1.69 | 0.65 | 4.18 |
| deoB;drm;lmo1954 | Q8Y5V1 | 2.07 | 3.22 | 0.05 | 0.23 | 4.62 | 2.50 | -0.55 | 3.42 |

| | | | | | | | | | |
|---|---|---|---|---|---|---|---|---|---|
| lmo0341 | Q8YA24 | 2.06 | 2.82 | 0.20 | 0.99 | 3.54 | 4.30 | -0.37 | 2.06 |
| **Putative ClpP2 interactors** | | | | | | | | | |
| lmo2167 | Q8Y5A2 | 0.00 | NaN | -0.07 | 0.14 | 7.27 | 5.36 | 0.03 | 0.04 |
| lmo1579 | Q8Y6V2 | 1.10 | 1.06 | -0.01 | 0.03 | 6.77 | 2.05 | 0.96 | 5.45 |
| lmo0560 | Q8Y9G8 | 1.60 | 2.99 | 0.05 | 0.26 | 6.70 | 2.26 | -0.68 | 5.97 |
| cadA;lmo1100 | P58414 | 3.81 | 0.91 | -0.02 | 0.04 | 6.28 | 1.35 | -1.58 | 5.33 |
| lmo1510 | Q8Y716 | 1.91 | 1.19 | 0.10 | 0.19 | 6.26 | 3.40 | -0.41 | 1.15 |
| guaA;lmo1096 | Q8Y822 | 1.11 | 0.87 | 0.04 | 0.24 | 5.99 | 2.07 | -1.07 | 7.08 |
| pyrC;lmo1837 | Q8Y663 | 0.09 | 0.04 | -0.17 | 0.25 | 5.89 | 2.85 | -1.65 | 4.17 |
| thrC | Q8Y4A5 | 1.39 | 0.94 | 0.18 | 0.76 | 5.79 | 2.87 | -1.28 | 5.24 |
| lmo2569 | Q8Y486 | 1.63 | 0.39 | -0.06 | 0.21 | 5.62 | 1.55 | -1.08 | 4.51 |
| lmo0191 | Q8YAE0 | 1.44 | 1.28 | 0.12 | 0.55 | 5.59 | 2.54 | 0.17 | 0.56 |
| proS;lmo1319 | Q8Y7G2 | 0.85 | 1.71 | 0.07 | 0.27 | 5.52 | 2.12 | -0.23 | 1.31 |
| mntB;lmo1849 | Q8Y651 | 1.84 | 2.61 | -0.02 | 0.09 | 5.41 | 1.82 | -0.37 | 3.33 |
| fabH;lmo2202 | Q8Y573 | 1.58 | 3.70 | 0.07 | 0.36 | 5.40 | 1.76 | -0.54 | 2.98 |
| lmo2113 | Q8Y5F1 | 0.09 | 0.06 | 0.27 | 1.91 | 5.36 | 4.58 | 0.28 | 1.31 |
| hslO;lmo0222 | Q8YAC4 | 1.72 | 2.38 | -0.02 | 0.04 | 5.36 | 5.36 | -0.26 | 1.01 |
| lmo0907 | Q8Y8J4 | 1.50 | 2.25 | 0.00 | 0.00 | 5.35 | 2.79 | -0.09 | 0.36 |
| lmo1376 | Q8Y7B0 | -0.37 | 0.33 | 0.04 | 0.31 | 5.26 | 2.23 | -0.41 | 5.09 |
| glnA | Q8Y7I0 | -0.62 | 1.10 | 0.16 | 2.72 | 5.25 | 1.94 | 0.00 | 0.01 |
| lmo2475 | Q8Y4G8 | 0.49 | 0.50 | 0.06 | 0.36 | 5.24 | 3.41 | 0.63 | 5.93 |
| lmo0982 | Q8Y8C6 | -0.76 | 1.29 | 0.11 | 0.75 | 5.14 | 2.46 | 0.11 | 0.47 |
| dnaN | Q8YAW1 | 1.65 | 3.58 | 0.01 | 0.03 | 5.10 | 1.73 | -0.49 | 3.07 |
| deoD;lmo1856 | Q8Y644 | -0.39 | 0.53 | 0.16 | 0.74 | 5.09 | 2.31 | 0.01 | 0.02 |
| lmo0534 | Q8Y9J3 | 0.00 | NaN | 0.10 | 0.95 | 5.01 | 4.89 | -0.86 | 3.71 |
| glyA;lmo2539 | Q8Y4B2 | 0.09 | 0.06 | 0.14 | 0.41 | 4.97 | 1.99 | 0.96 | 3.96 |
| lmo0978 | Q8Y8D0 | 1.79 | 3.90 | -0.01 | 0.05 | 4.92 | 2.02 | 0.18 | 0.92 |
| hfq;lmo1295 | Q92C58 | 1.22 | 1.14 | NaN | NaN | 4.89 | 3.45 | NaN | NaN |
| ami | Q8Y496 | 0.00 | NaN | 0.03 | 0.15 | 4.87 | 3.21 | -0.56 | 3.99 |
| lmo2491 | Q8Y4F3 | 1.57 | 0.97 | 0.21 | 0.83 | 4.83 | 5.99 | 0.26 | 1.19 |
| lmo2413 | Q8Y4M4 | 0.00 | NaN | 0.02 | 0.05 | 4.75 | 3.41 | 0.01 | 0.03 |
| fabD | Q8Y689 | 0.26 | 0.17 | 0.02 | 0.09 | 4.65 | 3.17 | -0.23 | 1.48 |
| lmo2192 | Q8Y581 | 1.87 | 1.97 | 0.18 | 0.41 | 4.54 | 1.66 | 0.23 | 0.63 |
| pyrF;lmo1832 | P58641 | 1.72 | 1.02 | -0.38 | 0.36 | 4.52 | 2.60 | -1.25 | 1.78 |
| lmo0098 | Q8YAM0 | 1.55 | 1.97 | -0.13 | 0.21 | 4.52 | 1.72 | -0.57 | 1.67 |
| citC | Q8Y6W5 | 0.49 | 1.44 | 0.12 | 0.42 | 4.51 | 5.28 | 0.88 | 6.07 |
| tpx;lmo1583 | Q8Y6U8 | 0.49 | 0.43 | 0.15 | 0.15 | 4.48 | 2.59 | -0.06 | 0.06 |
| lmo0078 | Q8YAN9 | 2.11 | 0.98 | 0.01 | 0.02 | 4.44 | 5.02 | -0.50 | 3.06 |
| pfkA;pfk;lmo1571 | Q8Y6W0 | -1.27 | 2.28 | -0.09 | 0.48 | 4.42 | 2.08 | -0.03 | 0.05 |
| lmo1688 | Q8Y6J6 | 1.83 | 3.84 | 0.10 | 0.23 | 4.38 | 2.10 | 0.53 | 2.11 |
| pyrG;lmo2559 | Q8Y495 | 0.32 | 0.28 | 0.02 | 0.12 | 4.36 | 1.65 | -0.40 | 3.20 |
| rsfS;lmo1486 | Q92BM7 | 0.00 | NaN | -0.09 | 0.13 | 4.35 | 2.34 | 0.62 | 1.59 |
| aroA;aroE;lmo1923 | Q8Y5Y0 | 1.45 | 1.74 | 0.00 | 0.01 | 4.33 | 5.23 | -0.62 | 5.99 |
| glmS;lmo0727 | Q8Y915 | 0.40 | 1.22 | 0.01 | 0.05 | 4.32 | 2.21 | 0.28 | 1.98 |
| aspB | Q8Y606 | 1.57 | 1.17 | 0.05 | 0.14 | 4.32 | 2.80 | -0.32 | 2.09 |
| lmo1017 | Q8Y897 | 1.58 | 0.73 | 0.03 | 0.14 | 4.31 | 2.25 | 0.28 | 2.62 |
| codY;lmo1280 | Q8Y7J7 | 1.53 | 2.16 | -0.03 | 0.13 | 4.23 | 2.13 | -0.43 | 3.54 |
| lmo1603 | Q8Y6T0 | 0.72 | 0.47 | 0.13 | 0.70 | 4.22 | 2.53 | -0.36 | 2.30 |

| | | | | | | | | | |
|---|---|---|---|---|---|---|---|---|---|
| cydA | Q927C3 | 1.48 | 0.87 | 0.31 | 0.75 | 4.18 | 5.70 | -1.03 | 3.46 |
| lmo0592 | Q8Y9D8 | 1.19 | 1.90 | -0.03 | 0.04 | 4.17 | 2.53 | 0.46 | 1.03 |
| lmo2638 | Q8Y435 | 1.68 | 2.18 | 0.15 | 0.43 | 4.17 | 1.58 | -0.81 | 3.87 |
| pepT;lmo1780 | Q8Y6B1 | -0.96 | 1.15 | 0.04 | 0.24 | 4.14 | 1.82 | 0.87 | 6.78 |
| lmo0595 | Q8Y9D5 | 0.00 | NaN | 0.09 | 0.37 | 4.13 | 3.55 | -0.64 | 3.30 |
| lmo1858 | Q8Y642 | -0.39 | 0.25 | 0.01 | 0.04 | 4.13 | 3.30 | 0.25 | 1.40 |
| lmo1621 | Q8Y6R3 | -0.76 | 0.70 | 0.04 | 0.11 | 4.13 | 2.42 | -0.02 | 0.08 |
| lmo1684 | Q8Y6K0 | 1.36 | 0.84 | 0.15 | 0.94 | 4.12 | 2.31 | -0.18 | 0.83 |
| lmo2354 | Q8Y4S8 | 1.52 | 0.85 | 0.11 | 0.25 | 4.11 | 3.88 | -0.10 | 0.33 |
| gyrA | Q8YAV6 | 2.39 | 0.71 | 0.06 | 0.24 | 4.08 | 3.21 | -0.29 | 2.27 |
| qoxB | Q8YAU9 | 1.25 | 1.01 | -0.01 | 0.04 | 4.07 | 3.92 | 0.42 | 1.19 |
| lmo0759 | Q8Y8Y3 | 1.36 | 1.02 | 0.25 | 0.55 | 4.04 | 4.80 | 0.43 | 1.47 |
| tilS/hprT;lmo0219 | Q8YAC7 | 1.32 | 0.82 | 0.09 | 0.19 | 4.01 | 2.50 | -0.33 | 1.25 |
| lmo1711 | Q92AU6 | 0.30 | 0.27 | 0.01 | 0.03 | 3.94 | 1.84 | 0.64 | 3.62 |
| lmo2188 | Q8Y583 | -0.21 | 0.13 | 0.06 | 0.38 | 3.94 | 2.09 | 0.78 | 4.47 |
| serC;lmo2825 | Q8Y3L0 | 1.74 | 1.29 | 0.11 | 1.08 | 3.93 | 3.29 | 0.56 | 5.16 |
| lmo1620 | Q8Y6R4 | -1.27 | 3.86 | 0.06 | 0.24 | 3.91 | 2.80 | 0.31 | 2.51 |
| atpD2;lmo2529 | Q8Y4C1 | 1.55 | 3.45 | 0.09 | 0.39 | 3.89 | 2.54 | 0.00 | 0.01 |
| secD;secF;lmo1527 | Q8Y703 | 2.35 | 1.10 | -0.02 | 0.08 | 3.87 | 4.61 | -0.24 | 2.11 |
| lmo1415 | Q8Y781 | -0.55 | 0.57 | 0.00 | 0.01 | 3.85 | 1.95 | -0.32 | 2.13 |
| lmo2193 | Q7AP53 | 2.94 | 1.23 | 0.14 | 0.32 | 3.84 | 1.68 | 0.08 | 0.20 |
| ade;adeC;lmo1742 | Q8Y6E5 | 0.00 | NaN | 0.09 | 0.19 | 3.83 | 3.81 | -0.21 | 0.49 |
| lmo0392 | Q92EP8 | 1.84 | 3.77 | 0.12 | 0.77 | 3.83 | 2.35 | 0.13 | 1.25 |
| lmo2692 | Q8Y3Y7 | 0.00 | NaN | 0.02 | 0.03 | 3.80 | 1.45 | -0.13 | 0.21 |
| aroA | Q8Y6T2 | 0.17 | 0.14 | 0.11 | 0.36 | 3.80 | 1.42 | -1.16 | 5.38 |
| dps;flp;fri;lmo0943 | Q8Y8G1 | 0.35 | 0.64 | 0.05 | 0.27 | 3.74 | 4.22 | -1.02 | 7.24 |
| lmo1393 | Q8Y797 | 1.75 | 2.51 | 0.03 | 0.27 | 3.73 | 1.78 | 0.06 | 0.41 |
| trxB;lmo2478 | O32823 | 1.26 | 3.21 | 0.07 | 0.19 | 3.70 | 2.92 | -0.29 | 1.01 |
| rpoA;lmo2606 | P66699 | 1.15 | 2.61 | -0.04 | 0.31 | 3.70 | 2.50 | -0.74 | 6.49 |
| ptsI;lmo1003 | O31149 | 1.40 | 2.15 | 0.04 | 0.10 | 3.65 | 3.38 | -0.12 | 0.32 |
| lmo2201 | Q8Y574 | 1.34 | 2.74 | -0.02 | 0.14 | 3.64 | 2.53 | 0.31 | 4.24 |
| ldh2;lmo1534 | Q8Y6Z6 | 1.72 | 0.81 | -0.06 | 0.24 | 3.61 | 2.85 | 0.49 | 3.73 |
| stp;lmo1821 | Q8Y678 | 1.07 | 0.56 | -0.04 | 0.24 | 3.61 | 4.91 | -0.30 | 1.30 |
| lmo2414 | Q8Y4M3 | 1.92 | 3.51 | 0.03 | 0.10 | 3.56 | 1.96 | 0.08 | 0.49 |
| lmo1578 | Q8Y6V3 | 1.58 | 2.52 | 0.03 | 0.05 | 3.52 | 2.30 | -0.02 | 0.04 |
| lmo2195 | Q8Y579 | 1.47 | 0.86 | 0.01 | 0.06 | 3.47 | 1.77 | -0.79 | 3.46 |
| lemA | Q8Y8E2 | 1.50 | 3.46 | 0.01 | 0.02 | 3.47 | 1.62 | -0.91 | 5.54 |
| pgi;lmo2367 | Q8Y4R7 | -1.35 | 2.36 | -0.02 | 0.09 | 3.46 | 2.18 | -0.41 | 2.38 |
| lmo1240 | Q8Y7N4 | 0.00 | NaN | 0.10 | 0.45 | 3.44 | 1.66 | -1.27 | 6.06 |
| ppaC;lmo1448 | Q8Y757 | -0.29 | 0.29 | 0.00 | 0.00 | 3.41 | 2.87 | -0.48 | 2.64 |
| hemL1;lmo1553 | Q8Y6X8 | 1.90 | 2.14 | 0.03 | 0.12 | 3.32 | 2.45 | 0.32 | 2.24 |
| pta | Q8Y5G0 | -0.25 | 0.46 | 0.04 | 0.09 | 3.32 | 1.33 | -0.19 | 0.55 |
| lmo0185 | Q8YAE5 | 1.98 | 3.35 | 0.05 | 0.17 | 3.31 | 3.38 | -0.15 | 0.68 |
| lmo2792 | Q8Y3P3 | 1.96 | 3.98 | 0.07 | 0.12 | 3.31 | 1.74 | 0.31 | 1.00 |
| rpsP;lmo1797 | Q8Y699 | 1.29 | 0.48 | 0.15 | 0.24 | 3.31 | 3.65 | -0.60 | 1.08 |
| ndk;lmo1929 | Q8Y5X4 | -0.15 | 0.05 | 0.09 | 0.67 | 3.29 | 2.54 | -0.28 | 2.08 |
| lmo2399 | Q8Y4N6 | 0.00 | NaN | -0.30 | 0.49 | 3.26 | 2.17 | 0.06 | 0.06 |
| lmo0437 | Q8Y9T6 | 0.00 | NaN | 0.26 | 2.29 | 3.25 | 3.61 | -2.04 | 7.26 |
| lmo1395 | Q8Y795 | 0.90 | 0.77 | -0.01 | 0.01 | 3.25 | 4.63 | -0.18 | 0.18 |

| | | | | | | | | | |
|---|---|---|---|---|---|---|---|---|---|
| lmo1374 | Q8Y7B2 | 1.24 | 0.39 | -0.06 | 0.26 | 3.24 | 3.98 | -0.07 | 0.25 |
| gap | Q8Y4I1 | -0.99 | 2.87 | 0.09 | 0.47 | 3.14 | 3.06 | -0.44 | 2.95 |
| fruA | Q8Y4U6 | 0.77 | 2.37 | 0.03 | 0.04 | 3.12 | 2.27 | -0.86 | 3.12 |
| lmo1255 | Q8Y7L9 | 1.19 | 1.48 | -0.05 | 0.06 | 3.11 | 2.27 | 0.26 | 0.37 |
| serS;lmo2747 | Q8Y3T4 | -0.02 | 0.01 | 0.05 | 0.29 | 3.11 | 1.86 | -0.02 | 0.07 |
| lmo0936 | Q8Y8G8 | 1.91 | 2.38 | 0.03 | 0.10 | 3.09 | 3.48 | -0.23 | 1.38 |
| fusA;fus;lmo2654 | Q8Y421 | 1.82 | 3.64 | 0.04 | 0.09 | 3.08 | 2.86 | -0.97 | 4.96 |
| lmo1739 | Q8Y6E8 | 0.00 | NaN | -0.03 | 0.05 | 3.04 | 3.99 | -1.65 | 3.86 |
| PdhD | Q8Y862 | 0.99 | 3.23 | -0.07 | 0.13 | 3.04 | 2.79 | -0.35 | 1.31 |
| menB | Q8Y6L1 | 0.16 | 0.23 | 0.04 | 0.22 | 2.99 | 2.48 | 0.15 | 1.08 |
| glmU;lmo0198 | Q8YAD4 | 1.71 | 3.65 | 0.13 | 0.58 | 2.99 | 3.03 | 0.05 | 0.31 |
| rpsR;lmo0046 | P66461 | 0.30 | 1.05 | -0.09 | 0.15 | 2.98 | 4.07 | -0.57 | 1.47 |
| hslV;clpQ;lmo1278 | Q8Y7J9 | 0.43 | 0.33 | -0.12 | 0.32 | 2.97 | 2.70 | -0.47 | 1.94 |
| lmo1738 | Q8Y6E9 | 0.00 | NaN | -0.02 | 0.06 | 2.96 | 1.79 | -2.63 | 8.06 |
| lmo1356 | Q8Y7C8 | 0.73 | 0.32 | 0.04 | 0.05 | 2.95 | 3.53 | -0.84 | 3.07 |
| purB | Q8Y6B8 | 1.83 | 1.19 | 0.14 | 0.67 | 2.95 | 2.32 | -0.17 | 1.17 |
| namA;lmo2471 | Q8Y4H1 | -1.00 | 1.23 | 0.01 | 0.02 | 2.94 | 1.84 | 0.61 | 3.42 |
| PdhB | Q8Y864 | 1.38 | 3.03 | 0.06 | 0.22 | 2.91 | 2.27 | -0.47 | 2.04 |
| lmo2767 | Q8Y3R6 | 1.48 | 3.29 | 0.08 | 0.15 | 2.89 | 2.17 | -0.73 | 2.16 |
| hemE;lmo2212 | Q8Y564 | -0.36 | 0.28 | 0.09 | 0.24 | 2.88 | 1.57 | 0.53 | 2.80 |
| pdxS;lmo2101 | Q8Y5G2 | 0.80 | 1.94 | 0.14 | 0.90 | 2.87 | 2.52 | -1.10 | 6.51 |
| ssb2;lmo2308;ssb1;lmo0045 | Q8YAR8;Q8Y4X1 | 1.48 | 1.67 | -0.11 | 0.15 | 2.87 | 1.64 | 0.09 | 0.16 |
| prs1;lmo0199 | Q48793 | 1.08 | 1.96 | 0.04 | 0.10 | 2.87 | 1.31 | -0.14 | 0.54 |
| rsbW;lmo0894 | Q8Y8K6 | 0.22 | 0.31 | 0.10 | 0.18 | 2.84 | 1.70 | 0.76 | 2.38 |
| cysK | Q8YAC3 | 0.26 | 0.61 | -0.04 | 0.12 | 2.81 | 3.68 | -0.61 | 3.15 |
| glyQ;lmo1459 | Q8Y753 | 1.09 | 2.50 | 0.02 | 0.04 | 2.80 | 2.06 | -0.80 | 3.73 |
| rbfA;lmo1327 | Q8Y7F4 | 0.00 | NaN | 0.03 | 0.12 | 2.76 | 2.90 | -1.05 | 5.41 |
| lmo2216 | Q8Y560 | 1.48 | 1.10 | -0.14 | 0.37 | 2.72 | 2.89 | 0.64 | 2.44 |
| lmo1818 | Q8Y681 | -0.44 | 0.77 | 0.02 | 0.04 | 2.67 | 1.64 | 0.01 | 0.03 |
| ftsH | Q8YAC6 | 0.88 | 1.30 | 0.03 | 0.17 | 2.65 | 1.70 | -0.05 | 0.44 |
| clpX;lmo1268 | Q8Y7K9 | 1.84 | 4.21 | -0.01 | 0.06 | 2.62 | 3.19 | 0.74 | 6.05 |
| pykA | Q8Y6W1 | -1.24 | 2.84 | 0.02 | 0.10 | 2.61 | 1.74 | 0.18 | 1.58 |
| lmo0406 | Q8Y9W3 | 0.00 | NaN | -0.13 | 0.20 | 2.51 | 2.38 | -0.06 | 0.10 |
| atpA2;lmo2531 | Q8Y4C0 | 1.44 | 4.06 | -0.04 | 0.15 | 2.47 | 2.41 | -0.29 | 1.88 |
| lmo0281 | Q8YA78 | 1.90 | 5.45 | 0.10 | 0.43 | 2.40 | 3.42 | 0.33 | 1.33 |
| lmo2077 | Q8Y5I5 | 0.00 | NaN | -0.02 | 0.03 | 2.37 | 3.16 | 0.56 | 1.70 |
| lmo0613 | Q8Y9B9 | 0.57 | 0.19 | 0.06 | 0.09 | 2.34 | 2.61 | -1.14 | 3.74 |
| lmo0533 | Q8Y9J4 | 0.00 | NaN | 0.40 | 1.49 | 2.33 | 3.44 | -1.24 | 3.15 |
| lmo0888 | Q8Y8L0 | -0.13 | 0.10 | 0.12 | 0.59 | 2.32 | 3.29 | -0.22 | 1.20 |
| lmo1701 | Q8Y6I3 | 0.00 | NaN | -0.04 | 0.11 | 2.31 | 2.80 | -0.86 | 4.27 |
| nusB;lmo1359 | Q8Y7C6 | 0.72 | 0.42 | 0.01 | 0.02 | 2.31 | 4.47 | -0.67 | 2.04 |
| lmo1691 | Q8Y6J3 | -1.58 | 3.31 | 0.03 | 0.13 | 2.30 | 3.42 | 0.04 | 0.13 |
| rimM;lmo1793 | Q8Y6A2 | 0.00 | NaN | 0.00 | 0.01 | 2.29 | 2.18 | -0.33 | 1.66 |
| lmo1065 | Q8Y853 | 2.20 | 1.19 | 0.11 | 0.53 | 2.28 | 1.41 | 0.11 | 0.45 |
| rpsE;lmo2615 | Q8Y446 | 0.80 | 4.11 | -0.20 | 1.37 | 2.28 | 2.69 | -0.68 | 2.98 |
| lmo2115 | Q8Y5E9 | 0.49 | 0.26 | 0.01 | 0.02 | 2.27 | 2.79 | -1.57 | 5.75 |
| lmo2636 | Q8Y437 | 0.00 | NaN | -0.01 | 0.02 | 2.25 | 2.44 | -0.95 | 4.08 |
| thrB;lmo2545 | Q8Y4A6 | 1.74 | 3.25 | 0.16 | 0.51 | 2.23 | 2.31 | -1.43 | 5.42 |
| rpmF2;lmo2047 | P66207 | 0.73 | 1.24 | -0.18 | 0.05 | 2.21 | 5.84 | -1.83 | 0.69 |

| | | | | | | | | | |
|---|---|---|---|---|---|---|---|---|---|
| atpH;lmo2532 | Q8Y4B9 | 0.00 | NaN | 0.14 | 0.49 | 2.20 | 2.93 | -0.50 | 3.14 |
| lmo0278 | Q8YA81 | 1.99 | 2.72 | -0.01 | 0.04 | 2.19 | 4.06 | -0.10 | 0.56 |
| lmo0813 | Q8Y8T1 | 1.43 | 1.84 | 0.09 | 0.81 | 2.17 | 2.86 | -0.67 | 3.66 |
| lmo0758 | Q8Y8Y4 | 1.07 | 0.64 | 0.10 | 0.30 | 2.14 | 4.45 | 0.31 | 1.88 |
| lmo1604 | Q8Y6S9 | 1.20 | 1.46 | 0.14 | 0.30 | 2.14 | 1.31 | 0.19 | 0.60 |
| thiI;lmo1592 | Q8Y6U0 | 1.97 | 2.60 | 0.11 | 0.33 | 2.13 | 1.58 | -0.89 | 5.35 |
| lmo1231 | Q8Y7P2 | 1.71 | 3.05 | 0.04 | 0.28 | 2.12 | 2.58 | -0.17 | 1.51 |
| lmo0775 | Q8Y8W7 | 0.00 | NaN | 0.03 | 0.05 | 2.09 | 3.23 | 0.12 | 0.35 |
| pdhA | Q8Y865 | 1.22 | 2.81 | -0.13 | 0.66 | 2.08 | 1.99 | -0.77 | 4.81 |
| lmo0208 | P0A4Q8 | 1.54 | 4.46 | 0.03 | 0.03 | 2.08 | 2.89 | 0.28 | 0.45 |
| lmo0273 | Q8YA86 | 1.94 | 1.91 | -0.04 | 0.09 | 2.07 | 3.48 | -0.28 | 1.42 |
| lmo1637 | Q8Y6P7 | 0.81 | 0.67 | 0.03 | 0.09 | 2.05 | 2.92 | 0.10 | 0.40 |
| lmo2674 | Q8Y404 | -0.51 | 0.96 | -0.06 | 0.51 | 2.05 | 1.64 | -0.19 | 0.67 |
| rplW;lmo2630 | Q8Y441 | 0.28 | 0.35 | -0.10 | 0.41 | 2.04 | 1.65 | -0.39 | 1.63 |
| rplS;lmo1787 | O53083 | -0.23 | 0.55 | -0.19 | 0.61 | 2.00 | 1.46 | -1.19 | 6.53 |

**Table S12** Putative interaction partners ($log_2$ ratio ≥2 and $-log_{10}$ t-test p-value ≥1.3 in the XL-co-IP experiment, $log_2$ ratio <1 or $-log_{10}$ t-test p-value <1.3 in the whole proteome experiment) of ClpP1 and ClpP2 at 42 °C.

| | | XL-co-IP of ΔclpP2 | | Whole proteome of ΔclpP1 | | XL-co-IP of ΔclpP1 | | Whole proteome of ΔclpP2 | |
|---|---|---|---|---|---|---|---|---|---|
| **Gene name** | **Uniprot ID** | **$log_2$ ratio (anti-ClpP/isotype control)** | **$-log_{10}$ t-test p-value (anti-ClpP/ isotype control)** | **$log_2$ ratio (Δ*clpP1*/WT)** | **$-log_{10}$ t-test p-value (Δ*clpP1*/WT)** | **$log_2$ ratio (anti-ClpP/isotype control)** | **$-log_{10}$ t-test p-value (anti-ClpP/ isotype control)** | **$log_2$ ratio (Δ*clpP2*/WT)** | **$-log_{10}$ t-test p-value (Δ*clpP2*/WT)** |
| **Putative ClpP1 interactors** | | | | | | | | | |
| lmo2113 | Q8Y5F1 | 11.07 | 6.52 | 0.15 | 1.19 | -7.99 | 1.51 | -0.67 | 6.00 |
| lmo2723 | Q8Y3V8 | 9.41 | 6.59 | 0.14 | 0.14 | 0.00 | NaN | 4.34 | 9.06 |
| lmo0537 | Q8Y9J1 | 9.06 | 7.84 | 0.00 | NaN | 0.00 | NaN | 3.56 | 4.43 |
| polC | Q8Y7G1 | 8.50 | 7.38 | -0.68 | 0.40 | 0.00 | NaN | 3.83 | 3.81 |
| hrcA | P0DJM4 | 7.09 | 4.66 | 0.12 | 0.58 | 3.37 | 3.38 | 1.71 | 7.91 |
| lmo0230 | Q8YAB7 | 6.49 | 6.42 | 0.46 | 0.97 | 0.00 | NaN | 4.03 | 9.05 |
| lmo1866 | Q8Y634 | 6.48 | 3.78 | -0.19 | 1.08 | 0.00 | NaN | 1.75 | 8.90 |
| lmo1384 | Q8Y7A4 | 6.37 | 4.92 | -0.39 | 0.52 | 0.00 | NaN | 8.02 | 12.32 |
| lmo1684 | Q8Y6K0 | 6.22 | 4.28 | 0.02 | 0.02 | 1.62 | 0.90 | -0.17 | 0.50 |
| fhs | Q8Y624 | 6.19 | 5.66 | -0.36 | 1.85 | 0.00 | NaN | 3.41 | 9.96 |
| lmo0485 | Q8Y9P0 | 6.02 | 4.64 | 0.99 | 0.59 | 0.07 | 0.05 | 7.80 | 6.71 |
| lmo0964 | Q8Y8E0 | 5.96 | 5.78 | -0.14 | 0.71 | 0.00 | NaN | 3.83 | 11.37 |
| mcsB | Q48759 | 5.94 | 3.99 | -0.07 | 0.31 | 6.07 | 5.72 | 2.23 | 10.21 |
| hemH | Q8Y565 | 5.92 | 7.71 | -0.06 | 0.59 | -0.47 | 0.97 | 0.38 | 4.09 |
| lmo0898 | Q8Y8K3 | 5.82 | 6.47 | -0.21 | 0.81 | 0.00 | NaN | 1.91 | 8.58 |
| lmo0229 | Q7AP89 | 5.71 | 4.68 | -0.24 | 0.47 | 2.39 | 3.16 | 4.39 | 10.71 |
| lmo0931 | Q8Y8H3 | 5.49 | 4.23 | 0.06 | 0.54 | -0.94 | 0.43 | -0.30 | 2.25 |
| yhaM | Q8Y556 | 5.38 | 7.23 | -0.01 | 0.07 | 0.00 | NaN | -0.87 | 4.98 |
| fbp | Q8Y8R5 | 5.31 | 7.54 | 0.04 | 0.02 | 0.00 | NaN | 7.33 | 7.82 |
| lmo1875 | Q8Y625 | 5.14 | 5.47 | -0.24 | 0.33 | 0.00 | NaN | 3.70 | 7.41 |
| uvrB | Q8Y4F5 | 5.13 | 3.89 | 0.01 | 0.03 | 2.98 | 3.08 | 1.97 | 6.96 |

| | | | | | | | | | |
|---|---|---|---|---|---|---|---|---|---|
| lexA | Q8Y7H7 | 4.99 | 5.17 | -0.78 | 1.08 | 3.25 | 2.10 | 2.48 | 8.72 |
| lmo1067 | Q8Y851 | 4.87 | 5.32 | -0.08 | 0.35 | 0.57 | 0.45 | -0.85 | 5.65 |
| lmo1713 | Q8Y6H3 | 4.79 | 4.93 | -0.05 | 0.03 | 2.85 | 4.37 | 1.73 | 1.90 |
| prfC | Q8Y8C0 | 4.77 | 7.36 | 0.16 | 0.43 | 1.31 | 3.24 | 0.60 | 3.56 |
| lmo0785 | Q8Y8V7 | 4.77 | 4.45 | -0.33 | 0.31 | 4.32 | 7.26 | 2.89 | 6.81 |
| lmo1393 | Q8Y797 | 4.75 | 5.54 | -0.10 | 0.78 | -1.49 | 0.64 | 0.39 | 3.68 |
| lmo0983 | Q8Y8C5 | 4.72 | 5.42 | -0.15 | 0.44 | 0.00 | NaN | 2.13 | 8.65 |
| lmo0305 | Q8YA56 | 4.56 | 5.56 | -0.69 | 0.75 | 0.00 | NaN | 0.94 | 3.94 |
| lmo1431 | Q8Y770 | 4.54 | 5.64 | -0.67 | 0.66 | 0.00 | NaN | 5.43 | 7.07 |
| murC | Q8Y6S8 | 4.53 | 5.12 | 0.20 | 0.80 | 0.19 | 0.19 | 0.22 | 1.22 |
| mnmA | Q8Y714 | 4.53 | 4.96 | -0.31 | 1.74 | -1.27 | 0.72 | -0.01 | 0.03 |
| queA | Q8Y6Z9 | 4.50 | 4.39 | -0.08 | 0.30 | -0.33 | 0.16 | 0.39 | 3.59 |
| lmo0454 | Q8Y9R9 | 4.47 | 2.66 | -0.27 | 1.78 | 3.79 | 2.79 | 1.18 | 9.30 |
| dps | Q8Y8G1 | 4.47 | 4.27 | -0.09 | 0.25 | 0.90 | 0.35 | -1.66 | 4.94 |
| lmo2720 | Q8Y3W1 | 4.43 | 3.50 | 0.15 | 0.47 | -0.19 | 0.11 | 1.55 | 6.46 |
| recN | Q8Y7B8 | 4.40 | 2.44 | -0.07 | 0.33 | -0.11 | 0.05 | 0.93 | 8.22 |
| lmo2828 | Q8Y3K7 | 4.25 | 5.78 | 0.36 | 1.57 | 0.00 | NaN | 2.18 | 8.01 |
| lmo1084 | Q8Y834 | 4.25 | 5.22 | -0.01 | 0.02 | -0.82 | 0.37 | 0.16 | 1.53 |
| lmo2166 | Q8Y5A3 | 4.24 | 6.06 | 0.07 | 0.23 | 1.10 | 1.04 | -0.21 | 1.45 |
| metE | Q8Y6K3 | 4.21 | 7.85 | -0.52 | 0.28 | 0.00 | NaN | 2.67 | 3.62 |
| glmU | Q8YAD4 | 4.19 | 6.90 | -0.04 | 0.24 | -0.25 | 0.66 | -0.23 | 3.32 |
| pepC | O69192 | 4.19 | 5.52 | -0.16 | 0.14 | 0.00 | NaN | 5.17 | 10.63 |
| pflB | Q8Y786 | 4.13 | 6.64 | -0.26 | 1.35 | 1.28 | 1.47 | 1.70 | 7.93 |
| pheT | Q8Y6S6 | 4.10 | 4.08 | -0.01 | 0.04 | -2.08 | 1.05 | 0.27 | 1.71 |
| addA | Q8Y511 | 4.08 | 5.18 | -0.31 | 0.16 | 3.01 | 1.70 | 5.54 | 6.48 |
| lmo1921 | Q8Y5Y2 | 4.06 | 2.29 | -0.60 | 0.64 | 0.00 | NaN | 7.75 | 10.22 |
| lmo1782 | Q8Y6A9 | 4.02 | 3.86 | -0.09 | 0.43 | 0.00 | NaN | 0.22 | 1.21 |
| lmo1825 | Q8Y674 | 3.99 | 5.85 | 0.04 | 0.23 | 2.22 | 1.09 | -0.56 | 5.50 |
| lmo1922 | Q8Y5Y1 | 3.92 | 3.18 | -0.11 | 0.62 | 1.45 | 1.10 | 0.42 | 3.56 |
| lmo0227 | Q8YAB9 | 3.92 | 6.37 | -0.29 | 1.39 | 2.95 | 3.39 | 4.01 | 11.10 |
| lmo1867 | Q8Y633 | 3.89 | 2.81 | -0.74 | 1.32 | 0.00 | NaN | 4.48 | 7.63 |
| kat | Q8Y3P9 | 3.75 | 6.64 | -0.22 | 1.12 | 0.31 | 0.56 | 2.67 | 7.93 |
| ssb1;ssb2 | Q8YAR8;Q8Y4X1 | 3.71 | 3.68 | -0.22 | 0.71 | -0.39 | 0.28 | -1.67 | 6.63 |
| nusA | Q8Y7F9 | 3.71 | 2.45 | 0.06 | 0.52 | -0.36 | 0.41 | -0.45 | 6.51 |
| tyrS | Q8Y6T4 | 3.69 | 5.35 | 0.03 | 0.13 | 1.53 | 0.84 | -0.14 | 1.23 |
| obg | Q8Y6Z3 | 3.69 | 4.53 | -0.46 | 0.38 | 0.00 | NaN | 1.61 | 5.92 |
| tal1 | Q8Y3T8 | 3.62 | 4.07 | -0.08 | 0.11 | 0.86 | 0.32 | -0.66 | 2.07 |
| lmo1217 | Q8Y7Q5 | 3.58 | 4.51 | 0.00 | 0.00 | 0.00 | NaN | -0.12 | 0.51 |
| lmo0291 | Q8YA68 | 3.55 | 5.26 | -0.20 | 0.12 | 0.00 | NaN | 6.14 | 7.83 |
| minC | Q8Y6Y6 | 3.54 | 5.42 | 0.09 | 0.33 | 1.34 | 1.58 | 0.15 | 1.34 |
| glpD | Q8Y7I4 | 3.49 | 4.23 | 0.11 | 0.40 | 2.13 | 1.89 | 1.09 | 5.05 |
| lmo2759 | Q8Y3S3 | 3.44 | 1.74 | 0.03 | 0.16 | -0.56 | 0.22 | 0.61 | 6.39 |
| pyrH | P65927 | 3.44 | 3.54 | 0.03 | 0.17 | 2.53 | 1.28 | -0.48 | 5.06 |
| fabH | Q8Y573 | 3.44 | 1.67 | 0.05 | 0.21 | 0.70 | 0.62 | -0.02 | 0.09 |
| lmo2168 | Q8Y5A1 | 3.43 | 4.66 | -0.23 | 0.52 | 0.00 | NaN | 3.09 | 7.62 |
| msrB | Q8Y641 | 3.43 | 4.88 | -0.30 | 1.40 | 0.00 | NaN | 0.74 | 3.66 |
| mnmG | Q8Y3M5 | 3.42 | 5.03 | -0.18 | 1.99 | 0.44 | 0.86 | 0.29 | 3.67 |

| | | | | | | | | | |
|---|---|---|---|---|---|---|---|---|---|
| lmo1814 | Q8Y683 | 3.38 | 4.17 | -0.03 | 0.15 | 1.54 | 1.34 | -0.49 | 4.08 |
| rpsQ | Q927L6 | 3.34 | 3.51 | -0.03 | 0.03 | -0.72 | 2.67 | 0.75 | 3.81 |
| topB | Q8Y3S5 | 3.32 | 4.26 | 0.02 | 0.01 | 0.00 | NaN | 5.96 | 5.54 |
| addB | Q8Y510 | 3.30 | 3.12 | -2.72 | 1.91 | 0.00 | NaN | 5.61 | 4.17 |
| trpD | Q8Y6Q3 | 3.30 | 3.92 | 0.52 | 0.24 | 2.61 | 3.73 | 2.86 | 2.51 |
| lmo1414 | Q8Y782 | 3.29 | 3.23 | -0.14 | 0.35 | 0.40 | 0.52 | -0.22 | 0.96 |
| hslV | Q8Y7J9 | 3.27 | 6.49 | 0.11 | 0.32 | 1.10 | 0.59 | -0.53 | 2.96 |
| deoC | Q8Y5R1 | 3.27 | 4.10 | 0.09 | 0.39 | 0.00 | NaN | 1.81 | 8.20 |
| lmo0978 | Q8Y8D0 | 3.22 | 1.41 | -0.31 | 1.72 | 0.31 | 1.12 | -0.40 | 4.89 |
| purL | Q8Y6C1 | 3.22 | 4.78 | -0.91 | 1.92 | 0.00 | NaN | 2.50 | 7.48 |
| pheT | Q8Y7Q1 | 3.21 | 2.90 | -0.05 | 0.21 | 1.03 | 0.54 | -0.86 | 6.22 |
| lmo1812 | Q8Y685 | 3.21 | 3.76 | -0.31 | 0.62 | 0.00 | NaN | 0.19 | 0.93 |
| pcrA | Q8Y6C9 | 3.20 | 2.42 | 0.07 | 0.40 | 0.91 | 0.74 | 0.00 | 0.01 |
| glyS | Q8Y754 | 3.19 | 2.71 | 0.14 | 0.79 | 1.33 | 1.64 | -0.47 | 5.16 |
| lmo0823 | Q8Y8S1 | 3.15 | 5.00 | -0.04 | 0.19 | -0.76 | 0.38 | 0.99 | 6.87 |
| murD | Q8Y5M1 | 3.13 | 3.07 | -0.04 | 0.24 | -0.09 | 0.03 | -0.44 | 5.06 |
| opuCA | Q7AP65 | 3.12 | 5.92 | -0.21 | 2.44 | 1.95 | 1.03 | -0.47 | 5.68 |
| smc | Q8Y692 | 3.11 | 3.98 | 0.11 | 0.71 | 0.00 | NaN | 0.34 | 3.07 |
| mutS | Q8Y789 | 3.10 | 3.25 | -0.08 | 0.53 | 1.28 | 0.71 | -0.60 | 6.71 |
| lmo2048 | Q8Y5L1 | 3.07 | 3.06 | 0.01 | 0.06 | -1.48 | 0.93 | -0.77 | 4.59 |
| lmo1028 | Q92CZ4 | 3.05 | 4.34 | 0.11 | 0.51 | -2.03 | 1.15 | -1.04 | 5.95 |
| citB | Q8Y6P3 | 3.04 | 3.72 | 0.04 | 0.18 | -0.24 | 0.09 | 0.32 | 2.94 |
| lmo1283 | Q8Y7J4 | 3.02 | 5.44 | 0.07 | 0.32 | -0.85 | 0.45 | 0.48 | 3.92 |
| lmo2707 | Q8Y3X2 | 3.01 | 4.54 | 0.16 | 0.57 | -0.69 | 0.79 | -0.98 | 3.47 |
| lmo0356 | Q8YA10 | 3.00 | 2.61 | 0.10 | 0.48 | 0.84 | 0.29 | -0.01 | 0.17 |
| lmo1609 | Q8Y6S4 | 2.99 | 2.25 | -1.21 | 0.56 | -0.27 | 0.14 | -0.11 | 0.57 |
| lmo1078 | Q8Y840 | 2.98 | 6.53 | -0.14 | 0.99 | 1.14 | 4.87 | -0.55 | 4.87 |
| menE | P58730 | 2.95 | 6.73 | -0.91 | 0.88 | 0.00 | NaN | 2.92 | 3.93 |
| lmo2426 | Q8Y4L1 | 2.94 | 3.42 | 0.02 | 0.04 | -1.06 | 0.71 | -0.51 | 2.42 |
| lmo1457 | P67195 | 2.93 | 4.76 | 0.23 | 1.79 | 1.90 | 4.96 | 0.20 | 2.66 |
| lmo2712 | Q8Y3W7 | 2.92 | 2.39 | 0.00 | NaN | 2.61 | 4.62 | 1.41 | 3.25 |
| lmo2437 | Q8Y4K1 | 2.92 | 5.14 | -0.67 | 1.49 | 0.00 | NaN | 2.38 | 5.28 |
| lmo0273 | Q8YA86 | 2.91 | 2.94 | -1.20 | 1.50 | 0.00 | NaN | 1.57 | 7.63 |
| lmo1634 | Q8Y6Q0 | 2.90 | 5.16 | -0.01 | 0.01 | 0.00 | NaN | -3.50 | 8.97 |
| lmo0774 | Q8Y8W8 | 2.87 | 1.91 | 0.08 | 0.27 | 0.45 | 0.27 | -0.57 | 3.76 |
| ecfA2 | Q8Y455 | 2.87 | 2.49 | 0.16 | 0.73 | 1.04 | 1.07 | -0.53 | 5.62 |
| lmo1710 | Q92AU7 | 2.85 | 3.31 | 0.06 | 0.13 | 1.96 | 3.64 | -1.23 | 5.51 |
| lmo2411 | Q928M6 | 2.84 | 5.29 | 0.07 | 0.74 | 0.31 | 1.68 | -0.37 | 4.34 |
| iap | P21171 | 2.80 | 4.47 | 0.44 | 1.56 | 0.00 | NaN | 1.42 | 6.58 |
| lmo1813 | Q8Y684 | 2.80 | 1.84 | -0.36 | 1.24 | 3.15 | 2.81 | 1.30 | 7.18 |
| lmo0387 | Q8Y9Y0 | 2.80 | 1.62 | 0.27 | 1.07 | -3.30 | 2.03 | 0.25 | 1.09 |
| mbl | Q8Y4C5 | 2.78 | 2.72 | 0.10 | 0.42 | 1.89 | 3.90 | -1.32 | 10.06 |
| lmo0267 | Q8YA92 | 2.77 | 4.33 | -0.08 | 0.28 | 0.00 | NaN | 0.45 | 2.83 |
| ruvB | Q8Y6Z8 | 2.77 | 4.22 | 0.10 | 0.35 | 0.00 | NaN | 0.95 | 6.59 |
| purH | Q8Y6C5 | 2.75 | 3.60 | -0.16 | 0.69 | 0.00 | 0.00 | 0.28 | 2.52 |
| lmo0287 | Q8YA72 | 2.72 | 3.93 | 0.04 | 0.32 | -1.11 | 0.55 | -0.13 | 0.91 |
| prmA | P0DJO9 | 2.70 | 3.02 | -0.02 | 0.06 | 1.71 | 2.56 | 0.30 | 2.26 |

| | | | | | | | | | |
|---|---|---|---|---|---|---|---|---|---|
| map | Q8Y6H5 | 2.70 | 1.50 | -0.08 | 0.36 | 0.80 | 2.13 | -0.48 | 3.65 |
| tilS/hprT | Q8YAC7 | 2.68 | 4.92 | -0.14 | 1.10 | 0.73 | 0.39 | -0.77 | 5.92 |
| lmo1087 | Q8Y831 | 2.68 | 1.71 | -0.04 | 0.14 | 0.22 | 0.33 | 0.25 | 1.80 |
| serS | Q8Y3T4 | 2.68 | 2.95 | -0.05 | 0.16 | -3.09 | 3.80 | -0.88 | 5.16 |
| dat | P0DJL9 | 2.67 | 4.93 | -0.18 | 0.89 | 0.00 | NaN | 0.09 | 0.37 |
| dapF | Q8Y5N9 | 2.64 | 1.75 | 0.03 | 0.11 | -0.89 | 0.80 | 0.44 | 5.31 |
| recD2 | Q8Y717 | 2.63 | 3.63 | -0.30 | 0.19 | 0.00 | NaN | 4.01 | 4.92 |
| lmo1022 | Q8Y892 | 2.62 | 4.55 | -0.34 | 3.00 | 0.94 | 0.52 | -0.25 | 1.93 |
| azoR2 | Q8Y8V6 | 2.62 | 6.27 | 0.14 | 0.61 | -1.50 | 0.87 | -1.37 | 7.34 |
| lmo2217 | Q8Y559 | 2.61 | 2.50 | 0.11 | 0.32 | -0.99 | 0.67 | -0.33 | 3.12 |
| lmo0241 | Q92F34 | 2.61 | 4.84 | -0.09 | 0.52 | 0.13 | 0.04 | -0.99 | 5.88 |
| trmFO | Q8Y7K1 | 2.59 | 1.52 | -0.23 | 1.25 | 1.58 | 3.76 | 0.04 | 0.26 |
| gshAB | Q8Y3R3 | 2.57 | 4.17 | -0.23 | 0.66 | 0.00 | NaN | 1.25 | 7.13 |
| mecA | Q9RGW9 | 2.56 | 1.62 | 0.19 | 0.23 | 0.00 | NaN | 3.91 | 7.26 |
| lmo2358 | Q8Y4S4 | 2.51 | 3.14 | -0.22 | 1.30 | 0.00 | NaN | 1.54 | 8.56 |
| tsaD | Q8Y5I7 | 2.51 | 5.54 | -0.02 | 0.07 | 1.90 | 2.94 | 0.36 | 2.91 |
| hemL2 | Q8Y6J9 | 2.50 | 2.16 | 0.04 | 0.14 | -0.21 | 0.10 | -0.15 | 1.06 |
| clpE | Q8Y8B1 | 2.50 | 5.11 | 0.27 | 1.23 | 0.33 | 1.71 | 5.63 | 13.27 |
| lmo2031 | Q8Y5M6 | 2.50 | 4.33 | -0.13 | 0.77 | 0.00 | NaN | -0.69 | 4.32 |
| dltA | Q8Y8D4 | 2.46 | 3.72 | 1.44 | 0.75 | 0.00 | NaN | 3.52 | 2.47 |
| ligA | Q8Y6D0 | 2.45 | 4.31 | -0.08 | 0.50 | 0.00 | NaN | -0.10 | 0.71 |
| lmo0936 | Q8Y8G8 | 2.40 | 3.04 | -0.09 | 0.59 | 1.28 | 0.93 | -0.56 | 4.51 |
| rimM | Q8Y6A2 | 2.39 | 3.67 | -0.02 | 0.07 | 0.00 | NaN | -0.15 | 0.86 |
| lmo0941 | Q8Y8G3 | 2.38 | 2.63 | 0.01 | 0.01 | 0.00 | NaN | 1.64 | 5.69 |
| lmo1083 | Q8Y835 | 2.37 | 3.00 | -0.10 | 0.59 | 0.54 | 0.20 | -0.47 | 3.62 |
| rsmH | Q8Y5L7 | 2.36 | 2.07 | -0.14 | 0.64 | 0.00 | NaN | 0.07 | 0.52 |
| pnp | Q8Y5V2 | 2.36 | 2.78 | -0.18 | 0.61 | -0.43 | 0.41 | -1.23 | 6.01 |
| deoB | Q8Y5V1 | 2.36 | 4.43 | -0.28 | 1.06 | -0.76 | 0.40 | -0.69 | 3.28 |
| gcvPB | Q8Y7D3 | 2.36 | 2.83 | -0.52 | 0.91 | 0.00 | NaN | 0.85 | 2.16 |
| lmo0884 | Q8Y8L3 | 2.35 | 3.56 | -0.12 | 0.79 | 0.19 | 0.13 | 0.21 | 1.70 |
| lmo1042 | Q8Y875 | 2.33 | 3.49 | -0.25 | 0.29 | 0.00 | NaN | 4.17 | 6.29 |
| lmo1543 | Q8Y6Y8 | 2.33 | 2.88 | -0.45 | 0.60 | 0.00 | NaN | 0.63 | 1.32 |
| atpG | Q927W3 | 2.32 | 1.58 | -0.04 | 0.12 | 1.64 | 1.02 | -0.34 | 3.39 |
| aspS | Q8Y709 | 2.32 | 1.51 | 0.07 | 0.36 | 0.23 | 0.56 | -1.18 | 7.40 |
| lmo2590 | Q8Y465 | 2.31 | 2.90 | 0.08 | 0.19 | 0.00 | NaN | 1.01 | 5.03 |
| isdG | Q92EH3 | 2.31 | 2.40 | 0.00 | NaN | 0.00 | NaN | 5.63 | 7.43 |
| lmo2853 | Q8Y3I3 | 2.28 | 3.49 | 0.23 | 1.26 | -0.95 | 2.84 | -0.38 | 2.25 |
| lmo0483 | Q8Y9P1 | 2.27 | 4.56 | -0.23 | 1.04 | -1.17 | 0.85 | -0.77 | 3.38 |
| cysS | Q8YAB1 | 2.26 | 1.92 | -0.02 | 0.21 | -1.32 | 0.47 | -0.95 | 9.51 |
| lmo1680 | Q8Y6K4 | 2.23 | 2.78 | 0.01 | 0.02 | 1.12 | 2.01 | 0.48 | 1.52 |
| lmo1239 | Q8Y7N5 | 2.22 | 3.53 | -0.17 | 0.53 | 0.00 | NaN | -0.41 | 3.09 |
| lmo1231 | Q8Y7P2 | 2.20 | 4.29 | -0.15 | 1.82 | 0.00 | NaN | -0.24 | 2.73 |
| lmo1223 | Q8Y7Q0 | 2.20 | 2.99 | -0.59 | 0.65 | 1.12 | 1.18 | 0.47 | 2.72 |
| murI | Q8Y7N7 | 2.19 | 3.08 | -0.13 | 0.57 | 1.37 | 2.24 | -1.22 | 8.61 |
| pheS | Q8Y7Q2 | 2.18 | 2.70 | -0.05 | 0.16 | -0.12 | 0.06 | -0.88 | 6.74 |
| upp | Q8Y4B3 | 2.18 | 4.37 | -0.02 | 0.11 | 0.21 | 0.33 | -0.97 | 7.04 |
| ychF | Q926X1 | 2.15 | 4.05 | 0.00 | 0.01 | -0.21 | 0.14 | -0.98 | 9.29 |

| | | | | | | | | | |
|---|---|---|---|---|---|---|---|---|---|
| lmo1514 | Q8Y712 | 2.13 | 2.17 | 0.18 | 0.09 | 0.00 | NaN | 3.47 | 3.90 |
| metG | Q8YAF2 | 2.13 | 1.65 | -0.05 | 0.32 | 0.19 | 0.07 | -0.86 | 6.30 |
| msrA | Q8Y640 | 2.13 | 2.38 | -1.99 | 1.91 | 0.00 | NaN | 3.51 | 8.70 |
| lmo1091 | Q8Y827 | 2.13 | 2.08 | 0.14 | 0.49 | 0.69 | 0.55 | -0.27 | 0.96 |
| lmo1336 | Q8Y7E7 | 2.12 | 4.43 | -0.62 | 0.44 | 0.00 | NaN | 1.71 | 6.78 |
| ami | Q8Y496 | 2.11 | 2.16 | -0.18 | 0.66 | 0.00 | NaN | -0.96 | 6.42 |
| lmo0557 | Q8Y9H1 | 2.08 | 4.55 | 0.00 | NaN | 0.00 | NaN | 6.49 | 8.77 |
| nrdR | Q8Y6W9 | 2.08 | 2.46 | -0.12 | 0.45 | 0.39 | 0.17 | -0.42 | 2.92 |
| lmo0811 | Q8Y8T3 | 2.07 | 6.84 | 0.13 | 0.24 | -0.82 | 3.41 | 3.78 | 9.29 |
| lmo0191 | Q8YAE0 | 2.05 | 1.61 | 0.05 | 0.18 | -1.04 | 0.31 | -0.54 | 2.99 |
| hisC | Q8Y5X8 | 2.05 | 4.96 | 0.02 | 0.02 | 0.00 | NaN | 5.10 | 9.41 |

**Putative ClpP1 and ClpP2 interactors**

| | | | | | | | | | |
|---|---|---|---|---|---|---|---|---|---|
| nadK2 | P65770 | 9.68 | 5.08 | -0.07 | 0.20 | 6.63 | 4.47 | -0.73 | 3.71 |
| lmo1021 | Q8Y893 | 7.77 | 7.04 | -0.24 | 1.04 | 5.97 | 6.12 | -0.79 | 3.18 |
| parB | Q8Y3P4 | 7.30 | 5.09 | 0.08 | 0.58 | 7.79 | 4.44 | -0.20 | 2.39 |
| lmo1576 | Q8Y6V5 | 6.41 | 6.58 | -0.08 | 0.12 | 3.53 | 3.01 | -0.23 | 0.66 |
| lmo1389 | Q8Y7A1 | 6.15 | 5.89 | 0.00 | 0.00 | 4.95 | 4.14 | -0.10 | 0.43 |
| lepA | Q8Y742 | 5.73 | 5.42 | -0.07 | 0.35 | 3.31 | 4.05 | -0.30 | 2.86 |
| mfd | Q8YAD0 | 5.72 | 4.88 | 0.01 | 0.03 | 4.51 | 5.72 | 0.06 | 0.83 |
| mutL | Q8Y788 | 5.41 | 5.60 | -0.04 | 0.13 | 4.23 | 3.48 | 0.50 | 4.77 |
| ansB | Q8Y6M1 | 5.39 | 1.92 | 0.01 | 0.02 | 2.45 | 3.44 | -0.31 | 3.39 |
| noc | Q8Y3P1 | 4.65 | 3.61 | 0.14 | 0.81 | 5.29 | 6.62 | 0.02 | 0.11 |
| dnaC | Q92FQ6 | 4.51 | 5.97 | 0.05 | 0.18 | 5.03 | 6.04 | 0.61 | 3.88 |
| prs2 | Q8Y9L8 | 4.47 | 5.30 | 0.03 | 0.09 | 2.43 | 6.86 | -0.46 | 4.57 |
| lmo2473 | P58588 | 4.35 | 4.67 | -0.13 | 0.56 | 3.01 | 2.80 | 0.38 | 3.32 |
| mnmE | Q8Y3M4 | 4.31 | 3.88 | 0.02 | 0.05 | 2.81 | 1.41 | 0.14 | 0.87 |
| lmo2263 | Q8Y515 | 4.17 | 5.09 | -0.52 | 0.70 | 4.38 | 5.67 | 0.02 | 0.02 |
| lmo1258 | Q8Y7L7 | 4.07 | 5.44 | 0.04 | 0.07 | 2.91 | 1.86 | 0.95 | 6.75 |
| gyrB | Q8YAV7 | 4.04 | 2.99 | -0.04 | 0.19 | 4.94 | 3.95 | -0.86 | 7.75 |
| proA | Q93Q55 | 3.95 | 2.77 | -0.08 | 0.57 | 4.74 | 5.01 | -0.35 | 3.03 |
| dnaA | Q8YAW2 | 3.84 | 5.63 | 0.32 | 1.80 | 2.04 | 5.56 | -0.48 | 3.32 |
| aroA | Q8Y5Y0 | 3.77 | 6.00 | -0.22 | 0.80 | 2.49 | 3.00 | 0.91 | 4.93 |
| murE | Q8Y5L9 | 3.75 | 2.32 | -0.08 | 0.34 | 2.65 | 1.91 | 0.21 | 1.87 |
| rnj | Q8Y767 | 3.69 | 3.49 | -0.08 | 0.58 | 2.03 | 2.87 | -0.47 | 5.17 |
| dnaX | Q8Y3X5 | 3.65 | 7.11 | 0.10 | 0.34 | 3.02 | 5.62 | 0.38 | 2.48 |
| ribC | Q8Y7F2 | 3.59 | 3.52 | 0.13 | 0.75 | 2.73 | 4.21 | 0.26 | 2.07 |
| lmo2247 | Q8Y529 | 3.58 | 4.42 | 0.03 | 0.16 | 2.66 | 2.67 | 0.11 | 0.78 |
| parE | Q8Y7J1 | 3.47 | 3.59 | -0.07 | 0.31 | 2.58 | 1.58 | -0.56 | 4.66 |
| lmo1577 | Q8Y6V4 | 3.44 | 4.97 | -0.32 | 0.98 | 2.43 | 5.04 | -0.94 | 3.91 |
| lmo1235 | Q8Y7N9 | 3.40 | 2.60 | -0.05 | 0.31 | 3.89 | 2.69 | -0.80 | 6.37 |
| lmo1721 | Q8Y6G6 | 3.27 | 4.24 | -0.12 | 0.59 | 2.47 | 4.64 | -0.23 | 1.61 |
| gcvPA | Q8Y7D4 | 3.25 | 2.54 | -0.41 | 0.87 | 2.88 | 4.49 | 0.65 | 2.03 |
| pnp | Q8Y7F1 | 3.24 | 4.86 | 0.04 | 0.24 | 2.66 | 2.44 | -0.55 | 4.97 |
| lmo1611 | Q8Y6S2 | 3.21 | 4.90 | -0.05 | 0.17 | 2.08 | 2.40 | -0.24 | 1.24 |
| rpmG1 | P66219 | 3.19 | 5.73 | 0.07 | 0.14 | 2.28 | 4.67 | -2.08 | 5.60 |
| lmo1647 | Q8Y6N7 | 3.09 | 4.12 | 0.20 | 0.80 | 2.84 | 1.84 | 0.04 | 0.21 |

| | | | | | | | | | |
|---|---|---|---|---|---|---|---|---|---|
| lmo1285 | Q8Y7J2 | 2.96 | 3.85 | -0.10 | 0.22 | 2.29 | 3.13 | -1.04 | 3.81 |
| azoR1 | Q8Y9C1 | 2.90 | 2.16 | 0.11 | 0.49 | 4.86 | 4.97 | 0.73 | 6.13 |
| proC | Q8Y9X2 | 2.89 | 4.18 | -0.14 | 0.63 | 3.99 | 6.51 | -0.73 | 6.47 |
| thrB | Q8Y4A6 | 2.84 | 4.44 | -0.11 | 0.30 | 3.87 | 5.64 | -1.24 | 4.90 |
| lmo0282 | Q8YA77 | 2.81 | 4.30 | 0.16 | 0.63 | 2.92 | 2.60 | 0.51 | 2.14 |
| parA | Q926W7 | 2.80 | 2.97 | -0.01 | 0.05 | 3.39 | 1.74 | -0.04 | 0.35 |
| dnaE | Q8Y6V7 | 2.78 | 3.20 | -0.06 | 0.26 | 4.12 | 4.91 | -0.27 | 2.67 |
| rnj | Q92CZ5 | 2.73 | 2.49 | -0.06 | 0.28 | 3.32 | 2.64 | -0.56 | 5.49 |
| lmo2114 | Q8Y5F0 | 2.73 | 3.80 | -0.08 | 0.55 | 2.73 | 2.54 | -0.11 | 0.50 |
| tagH | Q8Y843 | 2.65 | 1.89 | -0.05 | 0.17 | 4.72 | 3.16 | -0.19 | 1.52 |
| lmo2215 | Q8Y561 | 2.51 | 1.39 | -0.22 | 0.63 | 3.19 | 5.69 | 0.89 | 4.40 |
| lmo2390 | Q8Y4P5 | 2.45 | 4.37 | -0.01 | 0.04 | 3.35 | 2.32 | 0.78 | 7.09 |
| tdk | Q8Y4A7 | 2.42 | 2.64 | -0.11 | 0.64 | 2.03 | 2.62 | -1.01 | 6.84 |
| lmo0857 | Q8Y8N9 | 2.40 | 2.78 | 0.12 | 0.31 | 3.32 | 2.41 | 0.11 | 0.12 |
| trpB | Q8Y6Q6 | 2.38 | 1.48 | 0.10 | 0.27 | 3.07 | 5.91 | -0.94 | 3.92 |
| lmo0663 | Q8Y970 | 2.38 | 2.60 | -0.03 | 0.09 | 3.08 | 2.81 | 0.14 | 1.12 |
| lmo1612 | Q8Y6S1 | 2.36 | 2.95 | -0.13 | 0.31 | 3.01 | 3.77 | 0.33 | 2.70 |
| lmo1976 | Q8Y5S9 | 2.31 | 1.92 | -0.01 | 0.02 | 5.36 | 2.91 | -0.73 | 4.53 |
| lmo2557 | Q8Y497 | 2.28 | 3.69 | -0.16 | 1.32 | 3.57 | 3.93 | -0.27 | 1.31 |
| ispE | Q8YAE1 | 2.20 | 3.04 | 0.24 | 1.15 | 4.29 | 3.90 | 0.06 | 0.19 |
| dnaB | Q8Y6X0 | 2.17 | 2.44 | -0.10 | 0.51 | 3.04 | 2.44 | 0.18 | 1.72 |
| radA | Q48761 | 2.10 | 3.75 | 0.23 | 1.59 | 3.43 | 4.92 | 0.25 | 1.21 |
| ecfA1 | Q8Y454 | 2.00 | 4.28 | 0.01 | 0.03 | 3.49 | 2.38 | 0.22 | 2.10 |
| **Putative ClpP2 interactors** | | | | | | | | | |
| lmo2646 | Q8Y429 | 0.00 | NaN | -0.67 | 1.07 | 5.45 | 4.83 | -0.28 | 0.89 |
| thiM | Q8YA46 | 0.00 | NaN | -0.14 | 0.25 | 5.35 | 2.44 | -0.43 | 1.55 |
| lmo2643 | Q8Y431 | 1.69 | 0.91 | -0.05 | 0.16 | 5.26 | 3.14 | -0.04 | 0.14 |
| lmo1529 | Q8Y701 | -0.90 | 0.47 | 0.27 | 1.22 | 4.64 | 3.39 | -0.40 | 2.58 |
| lmo0737 | Q8Y905 | 0.00 | NaN | -0.03 | 0.05 | 4.47 | 4.38 | -4.93 | 6.86 |
| lmo0641 | Q8Y992 | 0.96 | 1.24 | -0.03 | 0.08 | 4.40 | 3.84 | -3.70 | 6.17 |
| lmo1652 | Q8Y6N2 | 0.00 | NaN | 0.42 | 0.63 | 4.32 | 5.24 | -0.24 | 0.33 |
| lmo0352 | Q8YA14 | 1.62 | 1.24 | -0.05 | 0.21 | 3.90 | 3.73 | -0.96 | 7.13 |
| aroB | Q8Y5X6 | 0.00 | NaN | -0.04 | 0.18 | 3.78 | 4.63 | 0.46 | 4.94 |
| lmo1919 | Q8Y5Y4 | 0.97 | 4.08 | -0.07 | 0.45 | 3.66 | 2.18 | -0.36 | 2.58 |
| lmo1358 | Q92BZ6 | 1.73 | 1.43 | 0.24 | 1.03 | 3.65 | 2.87 | -0.02 | 0.06 |
| lmo2404 | Q8Y4N2 | 0.00 | NaN | 0.21 | 0.95 | 3.60 | 4.80 | 0.46 | 3.02 |
| rpmG2 | P66221 | 1.18 | 0.80 | NaN | NaN | 3.60 | 4.32 | NaN | NaN |
| lmo2554 | Q8Y4A0 | 0.00 | NaN | 0.78 | 1.14 | 3.27 | 3.92 | 0.41 | 0.51 |
| lmo1511 | Q8Y715 | 0.00 | NaN | -0.11 | 0.19 | 3.22 | 4.36 | 0.19 | 1.11 |
| lmo0052 | Q8YAR3 | 1.71 | 1.37 | 0.13 | 0.63 | 3.17 | 4.60 | -0.11 | 0.93 |
| lmo2337 | Q8Y4U4 | 0.96 | 1.50 | 0.02 | 0.12 | 3.07 | 3.91 | -0.37 | 2.12 |
| lmo1930 | Q8Y5X3 | 0.00 | NaN | 0.00 | 0.00 | 3.05 | 3.52 | -0.10 | 0.25 |
| lmo1081 | Q8Y837 | 1.61 | 3.88 | -0.10 | 0.39 | 3.05 | 1.58 | 0.02 | 0.07 |
| dxr | Q8Y7G4 | 0.17 | 0.09 | -0.11 | 0.61 | 2.87 | 3.66 | 0.32 | 2.10 |
| lmo0763 | Q8Y8X9 | 0.00 | NaN | -0.15 | 0.28 | 2.84 | 3.63 | 0.36 | 2.23 |
| accA | Q8Y6V9 | 1.63 | 1.91 | 0.00 | 0.00 | 2.82 | 1.50 | -0.05 | 0.32 |

| | | | | | | | | | |
|---|---|---|---|---|---|---|---|---|---|
| glmS | Q8Y915 | 0.56 | 2.58 | 0.00 | 0.01 | 2.82 | 8.40 | 0.77 | 9.09 |
| polA | Q8Y6W6 | 1.57 | 1.94 | -0.02 | 0.10 | 2.75 | 3.65 | 0.39 | 4.26 |
| fruA | Q8Y4U6 | 0.52 | 2.19 | -0.16 | 1.23 | 2.75 | 5.01 | -0.70 | 4.98 |
| lmo0847 | Q8Y8P9 | 0.00 | NaN | 0.05 | 0.41 | 2.71 | 4.43 | -0.68 | 6.63 |
| lmo0739 | Q8Y903 | 0.00 | NaN | -0.11 | 0.32 | 2.69 | 2.32 | -2.10 | 6.13 |
| lmo2486 | Q8Y4F8 | 0.45 | 0.37 | -0.35 | 1.71 | 2.68 | 3.44 | -1.05 | 5.33 |
| rny | P0DJP2 | 1.47 | 1.69 | 0.06 | 0.58 | 2.62 | 3.18 | 0.07 | 0.63 |
| lmo2769 | Q8Y3R4 | 0.00 | NaN | -0.15 | 0.21 | 2.60 | 3.91 | -0.38 | 1.76 |
| lmo1369 | Q8Y7B7 | 1.26 | 0.93 | 0.04 | 0.12 | 2.60 | 1.95 | 0.28 | 1.33 |
| lmo2194 | Q8Y580 | 0.00 | NaN | -0.03 | 0.08 | 2.59 | 2.10 | -1.24 | 5.67 |
| lmo0077 | Q8YAP0 | 0.00 | NaN | -0.53 | 0.49 | 2.58 | 3.54 | -0.66 | 2.88 |
| birA | Q8Y5Z9 | 0.77 | 1.52 | 0.10 | 0.65 | 2.56 | 3.86 | -0.30 | 2.24 |
| plsX | Q8Y688 | 0.90 | 2.01 | -0.06 | 0.22 | 2.55 | 5.36 | -0.88 | 5.69 |
| lmo1373 | Q8Y7B3 | 1.24 | 4.40 | -0.16 | 0.73 | 2.54 | 6.12 | 0.00 | 0.01 |
| lmo0967 | Q92D54 | 0.00 | NaN | 0.09 | 0.70 | 2.53 | 1.77 | -0.05 | 0.55 |
| lmo1219 | Q8Y7Q3 | 0.00 | NaN | -0.13 | 0.38 | 2.52 | 5.98 | -1.09 | 5.47 |
| hom | Q8Y4A4 | 1.65 | 1.34 | 0.13 | 0.94 | 2.52 | 5.69 | -1.31 | 7.81 |
| ftsK | Q8Y7A3 | 1.49 | 2.94 | 0.27 | 1.12 | 2.51 | 3.49 | -0.66 | 6.72 |
| lmo2738 | Q8Y3U3 | 1.27 | 2.76 | NaN | NaN | 2.48 | 4.49 | NaN | NaN |
| tagD | Q8Y829 | 1.87 | 1.18 | 0.38 | 1.04 | 2.46 | 3.52 | 0.10 | 0.21 |
| lmo0826 | Q8Y8R8 | 0.00 | NaN | -0.46 | 0.49 | 2.46 | 4.07 | -1.07 | 4.22 |
| lmo1351 | Q8Y7D2 | 1.60 | 1.13 | 0.06 | 0.26 | 2.40 | 1.59 | -1.23 | 7.33 |
| lmo2705 | Q8Y3X4 | 1.90 | 2.22 | 0.62 | 2.36 | 2.38 | 1.34 | -0.11 | 0.34 |
| lmo1057 | Q8Y860 | 0.82 | 1.62 | -0.07 | 0.10 | 2.36 | 2.71 | -0.64 | 2.11 |
| lmo0317 | Q8YA45 | 0.00 | NaN | 0.52 | 0.44 | 2.35 | 1.67 | 0.26 | 0.13 |
| hly | P13128 | 0.83 | 0.96 | 1.24 | 4.14 | 2.35 | 4.55 | -1.57 | 5.03 |
| lmo1395 | Q8Y795 | 1.95 | 4.33 | 0.01 | 0.03 | 2.33 | 2.07 | 0.04 | 0.24 |
| lmo0292 | Q8YA67 | 0.00 | NaN | -0.55 | 1.77 | 2.27 | 1.65 | -0.89 | 4.55 |
| hemL1 | Q8Y6X8 | 1.22 | 0.74 | 0.01 | 0.02 | 2.26 | 1.47 | -0.23 | 2.98 |
| fni | Q8Y7A5 | 0.00 | NaN | 0.02 | 0.08 | 2.25 | 2.37 | -0.09 | 0.61 |
| ilvC | Q8Y5S0 | 0.00 | NaN | -0.88 | 0.45 | 2.24 | 2.71 | -2.24 | 0.99 |
| metN1 | Q8YA75 | 0.00 | NaN | 0.32 | 1.54 | 2.23 | 1.47 | -1.41 | 5.53 |
| lmo1741 | Q8Y6E6 | 0.00 | NaN | 0.11 | 0.44 | 2.20 | 4.39 | -0.45 | 3.89 |
| lipL | Q8Y489 | 0.00 | NaN | 0.07 | 0.26 | 2.18 | 4.32 | -0.74 | 4.17 |
| lmo2258 | Q8Y520 | 0.00 | NaN | -2.62 | 1.43 | 2.15 | 1.89 | -5.01 | 5.42 |
| lmo0667 | Q8Y966 | 0.54 | 1.53 | -0.18 | 0.78 | 2.14 | 3.33 | 0.96 | 5.11 |
| lmo1066 | Q8Y852 | 0.93 | 0.66 | 0.09 | 0.34 | 2.14 | 3.39 | -0.30 | 1.51 |
| lmo1401 | Q8Y791 | 0.22 | 0.13 | -0.10 | 0.56 | 2.13 | 3.90 | -0.59 | 6.22 |
| lmo0825 | Q8Y8R9 | 0.97 | 1.26 | -0.09 | 0.50 | 2.09 | 4.14 | -0.05 | 0.27 |
| dnaJ | P0DJM1 | 1.21 | 2.87 | 0.04 | 0.12 | 2.09 | 4.88 | -0.08 | 0.26 |
| lmo2550 | Q7AP48 | 0.00 | NaN | -0.09 | 0.22 | 2.06 | 2.16 | -0.36 | 1.13 |
| rpsE | Q8Y446 | 0.38 | 0.97 | 0.18 | 1.15 | 2.04 | 6.29 | -1.14 | 6.91 |
| lmo1438 | Q8Y763 | 0.00 | NaN | -0.14 | 0.65 | 2.03 | 2.92 | -0.95 | 6.67 |
| lmo2474 | Q8Y4G9 | 1.35 | 2.37 | 0.16 | 0.97 | 2.02 | 5.08 | -0.18 | 1.10 |
